# RNA decay defines the response to transcriptional perturbation in leukaemia

**DOI:** 10.1101/2022.04.06.487057

**Authors:** Izabela Todorovski, Breon Feran, Zheng Fan, Sreeja Gadipally, David Yoannidis, Isabella Y Kong, Stefan Bjelosevic, Magnus Zethoven, Edwin D Hawkins, Kaylene J Simpson, Gisela Mir Arnau, Anthony T Papenfuss, Ricky W Johnstone, Stephin J Vervoort

**Affiliations:** Peter MacCallum Cancer Centre, Melbourne, Victoria 3000, Australia; Sir Peter MacCallum Department of Oncology, The University of Melbourne, Victoria 3010, Australia; The Walter and Eliza Hall Institute of Medical Research, Parkville, Australia; Department of Medical Biology, The University of Melbourne, Parkville, Australia

## Abstract

Therapeutic targeting of dysregulated transcriptional programs has arisen as a promising strategy for the treatment of leukaemias. The therapeutic response to small molecule inhibitors of Bromodomain-Containing Proteins (BRD), such as BRD2 and BRD4, P300/cAMP-response element binding protein (CBP) and Cyclin Dependent Kinases (CDKs), is generally attributed to the selective disruption of oncogenic gene expression networks driven by enhancers, super-enhancers (SEs) and lineage-specific transcription factors (TFs), including the *c-MYC* oncogene. Using technologies such as thiol (SH)-linked alkylation for the metabolic sequencing of RNA sequencing (SLAM-seq) to profile messenger RNA (mRNA) decay and production rates, we demonstrate that gene intrinsic properties largely govern the selectivity associated with transcriptional inhibition, where total mRNA response signatures are dominated with genes that have short transcript half-lives, including those regulated by SEs and oncogenic TFs. Further highlighting that gene sensitivities only occur in the context of short transcript half-lives, stabilisation of the *c-MYC* transcript through changes in the 3’ UTR rendered it insensitive to transcriptional targeting. However, this was not sufficient to rescue *c-MYC* target gene transcription and anti-leukaemia effects following transcriptional inhibition. Importantly, long-lived mRNAs encoding essential genes that evade transcriptional targeting can be rendered sensitive via modulation of mRNA decay kinetics through inhibition of the RNA Binding Protein (RBP), ELAV Like RNA binding protein 1 (ELAVL1)/ Human Antigen R (HuR). Taken together, these data demonstrate that mRNA decay shapes the therapeutic response to transcriptional perturbation and can be modulated for novel therapeutic outcomes using transcriptional agents in leukaemia.

## Introduction

Genetic alterations in cancer can affect proteins involved in almost all regulatory steps of RNA Polymerase II (RNAPII) -driven gene expression. Dysregulated transcription ultimately contributes to cellular transformation and the development of malignant phenotypes (*1*, *2*). This includes leukaemias, where several chromosomal rearrangements, such as those involving the Mixed Lineage Leukemia (MLL) H3K4 methyltransferase, can drive the aberrant expression of leukaemogenic proteins (*3*).

As these transcriptionally dysregulated cancers exhibit a critical dependency on components of the core transcription machinery that regulate RNAPII, small molecule inhibitors that target core transcriptional enzymes and structural proteins have been developed for therapeutic use (*4*, *5*). In leukaemia treatment, this includes compounds that target the transcriptional co-activator p300/cAMP-response element binding protein (CBP), Bromodomain and Extra Terminal (BET) family of epigenetic reader proteins such as Bromodomain-Containing Protein 4 (BRD4) and transcriptional Cyclin Dependent Kinases (t-CDKs) (*6*–*10*).

A large number of publications studying the effects of small molecules targeting the core RNAPII machinery or epigenetic regulatory proteins show that they result in selective gene expression responses. The selectivity is generally attributed to gene-specific chromatin features, including occupancy of the targeted factor, cell-type specific core regulatory transcription factors (CR-TFs), enhancers and super-enhancers (SEs) (*1*, *11*, *12*), all of which impact the *de novo* production of mRNA. In the context of leukaemia this has been described to primarily affect oncogenic networks driven by key TFs such as *c-MYC* (*11*, *13*–*15*). Indeed, the sensitivity of *c-MYC* itself to transcriptional perturbation has largely been associated with a cluster of SEs 1.7 mega bases (Mb) from its transcription start site (TSS) and CR-TFs such as PU.1, MYB and FLI1 (*16*, *17*). Recent work has also demonstrated that co-factors can possess more broad or selective chromatin specificities, including CDK9 and p300/CBP and BRD4, respectively (*12*). Again, both broad and selective cofactor regulatory targets were associated with histone modifications, DNA sequences and TF motifs (*12*).

As the chromatin landscape is the focus in evaluating gene responses to transcriptional therapies, the impact of different stages of the mRNA life cycle to gene expression changes is rarely evaluated. Obtaining information about mRNA life cycle stages can be achieved by leveraging recent technological advances that enable the high-confidence measurement of mRNA production and decay rates genome-wide. This involves metabolic labelling of RNA with the nucleotide analog 4-thiouridine (4sU), and its quantification using affinity- or conversion-based technologies (*18*–*23*). For example, thiol (SH)-linked alkylation for the metabolic sequencing of RNA (SLAM-seq) is a conversion-based approach that uses 4sU dependent thymine-to-cytosine (T > C) conversions for the *in silico* separation of newly synthesized and pre-existing transcripts (*18*).

Using SLAM-seq to assess mRNA kinetics, we demonstrate that transcript decay rates largely dictate the response to transcriptional and epigenetic therapies. Importantly, we further highlight that genetic and small-molecule modulation of mRNA half-lives can alter the effects on transcript levels seen following inhibition of mRNA production and can provide enhanced anti-cancer responses when combined with agents that perturb the core RNAPII machinery.

## Results

### The anti-leukaemia responses to transcriptional inhibition occurs concomitantly with selective reduction in total mRNA levels

To investigate the role of mRNA kinetics in shaping responses to agents that perturb cofactors with selective (p300/CBP and BRD4; class II) and broad (CDK9 and RNAPII; class I) chromatin specificities (*12*), the biological response to small molecule inhibitors of p300/CBP (A-485), BRD4 (JQ1), CDK9 (AZ-5576) and RNAPII (actinomycin-D; ACTD) in the K562 Chronic Myeloid Leukaemia (CML) cell line was first assessed (Fig. 1A and S1). All agents reduced the number of K562 cells grown *in vitro* in a concentration and time-dependent manner (Fig. S1A). Moreover, the small molecule concentrations used to assess the anti-leukaemic effects were consistent with on-target activities, as demonstrated by a reduction in H3K18ac, c-MYC protein and RNAPII carboxy-terminal domain (CTD) Serine 2 (Ser2) phosphorylation following treatment with A-485, JQ1 and AZ-5576, respectively (Fig. S1. B) (*8*, *24*, *25*). Using the same leukaemia model and small molecule inhibitors, SLAM-seq was next performed to evaluate responses to transcriptional targeting (*18*) (Fig. 1A-B). Differential gene expression analysis (DGEA) of total mRNA read counts revealed that all transcriptional inhibitors affected the levels of only a subset of expressed genes (Fig. 1C, S2. A). Consistent with previous studies (*26*, *27*), 785 SE-associated genes defined using publicly available H3K27ac ChIP-seq data in K562 cells (Fig. S2. B) and 83 CR SE-associated TFs identified using the Coltron algorithm (Fig. S2. C-D) were significantly more down-regulated compared to all other genes across all treatments (Fig. 1D, S2. E-F). Transcripts repressed following treatment with A-485, JQ1, AZ-5576 and ACTD were enriched for genes encoding several cancer and inflammatory hallmark pathways, including c-MYC targets and cytokine signalling, respectively (Fig. S2. G). This selectivity was observed despite A-485, JQ1, AZ-5576 and ACTD causing largely distinct total mRNA changes, whereby CDK9i and ACTD signatures co-clustered and exhibited the highest degree of similarity (Fig. S2. H). However, the vast majority of genes were refractory to changes in total mRNA levels at these timepoints and inhibitor concentrations (Fig. 1C), including several SE associated genes and CR TFs (Fig. S2. E-F). Overall, these data validate that on the total mRNA level, transcriptional targeting largely perturbs the expression of SE- and CR-TF oncogenic gene networks.

**Figure 1.**
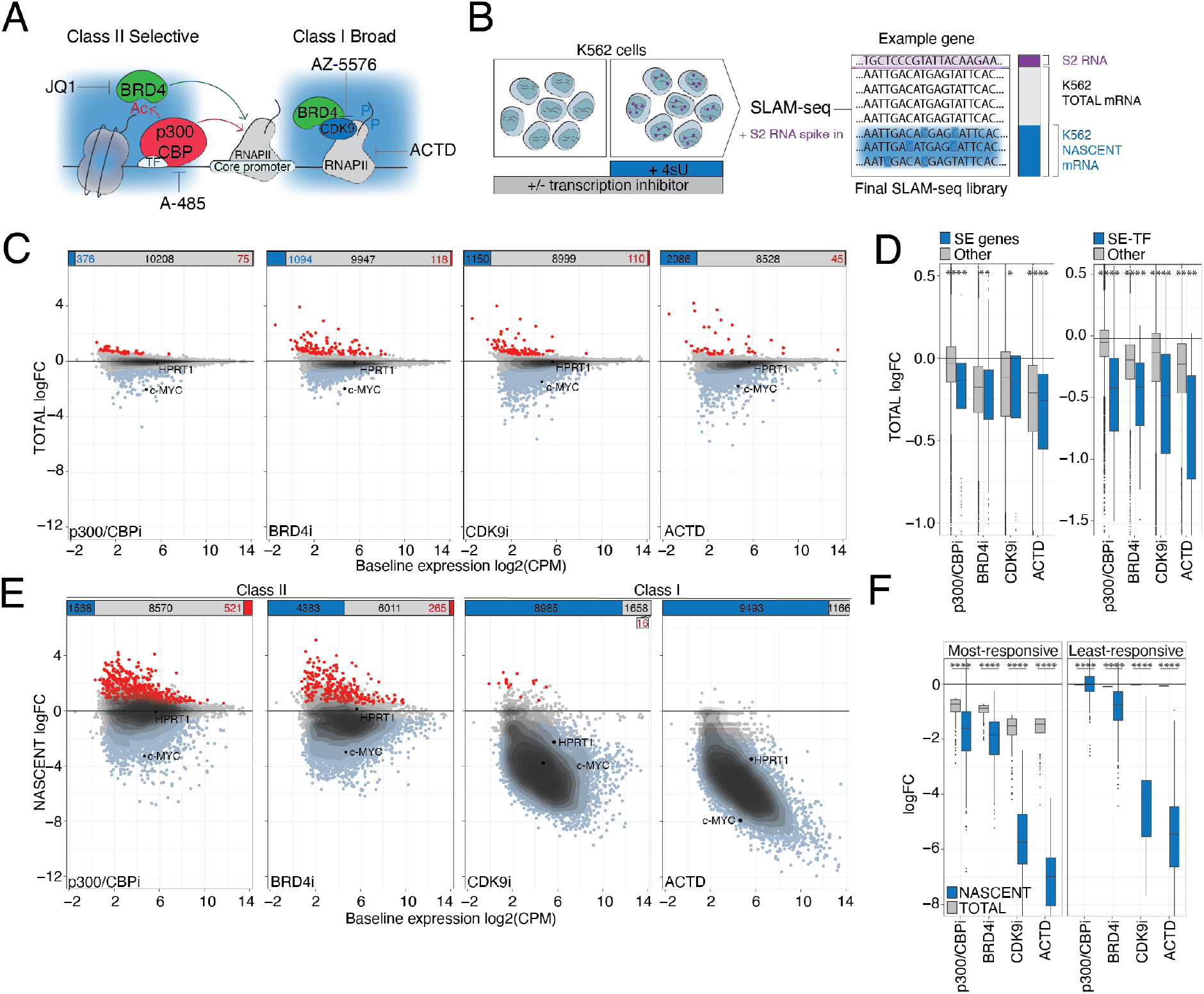
A defined subset of genes determines the therapeutic response to transcription inhibition. **(A)** Simplified schematic of transcription inhibition by A-485 (p300/CBPi), JQ1 (BRD4i), AZ-5576 (CDK9i) and Actinomycin-D (ACTD). **(B)** Schematic of SLAM-seq experimental procedure. Nascent reads are defined as RNA containing at least two thymine-to-cytosine (T>C) conversions and total reads are defined as the sum of unconverted and converted RNA. Schneider 2 (S2) RNA was spiked-in as an external reference control. **(C)** Scatterplot of baseline total mRNA expression versus change in spike-in normalized total gene expression upon two hours of transcription inhibition. Significantly up- or down-regulated genes highlighted in red and blue, respectively. Sum of significantly differentially altered events indicated in bar chart above. **(D)** Change in spike-in normalized total expression of **(left)** genes regulated by super-enhancers (SEs) and **(right)** core-regulatory transcription factors (SE-TF). Remaining genes indicated in grey. **(E)** Scatterplot of baseline nascent mRNA expression versus change in spike-in normalized total gene expression upon two hours of transcription inhibition. Significantly up- or down-regulated genes highlighted in red and blue, respectively. Sum of significantly differentially altered events indicated in bar chart above. (**F)** Change in spike-in normalized total and nascent expression of **(left)** most- and **(right)** least-responsive genes. p300/CBP: p300/cAMP-response element binding protein (CBP). BRD4: Bromodomain-Containing Protein 4. CDK9: Cyclin Dependent Kinase 9. RNAPII: RNA Polymerase II. TF: Transcription factor. Ac: Acetyl group. P: Phosphate group. 4sU: 4-thiouridine. H: Hour. 3’UTR: 3’ Untranslated Region. LogFC: log2 fold change relative to DMSO. Baseline expression: spike-in normalized total log2 CPM in DMSO-treated conditions. CPM: Counts Per Million. Significantly up-regulated: P Value < 0.05 & logFC > 0.5. Significantly down-regulated: P Value < 0.05 & logFC < −0.5. P Value < 0.0001 using an unpaired Wilcoxon test. Most-responsive: top 200 most significantly down-regulated genes (logFC < −0.5 and P Value < 0.05) using spike-in normalized total reads. Least-responsive: 200 unaltered (−0.25 < logFC < 0.25 and P Value > 0.05) genes using spike-in normalized total reads. AUC: Area Under the Curve.

### Nascent and total mRNA responses to transcriptional inhibition are disconnected

To investigate whether selective changes in total mRNA levels following small molecule inhibition of p300/CBP, BRD4, CDK9 and RNAPII are due to a concordant change in mRNA production, DGEA was performed on nascent read counts (Fig. 1B). In contrast to the effect seen on total mRNA (Fig. 1C, S2. A), nascent mRNA was more broadly repressed in response to all inhibitors (Fig. 1E, S3. A-C). The greatest effect on nascent mRNA expression was observed following treatment with AZ-5576 and ACTD and globally down-regulated the *de novo* mRNA synthesis of > 80% of genes (Fig. 1E, S3. A-C). In contrast, transcriptional inhibition with p300/CBPi and BRD4i, was selective and significantly reduced the mRNA synthesis of < 50% of expressed genes (Fig. 1E, S3. A-C). Importantly, changes in nascent gene expression with class II inhibitor treatment was still significantly greater in comparison to associated total mRNA changes (Fig. S3. C). In agreement with these data, comparison between nascent and total mRNA read counts revealed only a modest Pearson’s correlation coefficient (Fig. S3. D). Correlations were lower with class I inhibitors (0.17-0.3) in comparison to more selective transcriptional perturbation with class II compounds (0.43-0.46) (Fig. S3. D). As an example of the disconnect between changes in nascent and total mRNA abundance, genes such as *HPRT1* that were the least responsive (P-value > 0.05, −0.25 < logFC < 0.25; 200 genes closest to the mean logFC) to changes in total mRNA had significantly down-regulated nascent gene expression, especially in response to class I compounds (Fig. 1F, S3. E). In contrast, the proto-oncogene *c-MYC* was amongst the most responsive to transcriptional perturbation (top 200 genes most significantly down-regulated) on both total and nascent mRNA levels (Fig. 1F, S3. F). This highlights that in a subset of genes, down-regulation of *de novo* mRNA synthesis can result in a concomitant reduction in total mRNA. However, despite global down-regulation of nascent transcripts, most genes appear to be refractory to changes in total mRNA following acute transcriptional inhibition. Taken together, these data suggest that selective targeting of mRNA production is more limited than currently appreciated and infrequently translates into alterations in the global mRNA pool. Moreover, it further indicates that other parameters that define total mRNA levels underpin the observed responses to transcriptional perturbation.

### Gene-intrinsic mRNA decay properties shape the response to transcriptional inhibitors

Total transcript levels are critically determined by the decay component of the mRNA life-cycle (*28*). To investigate whether mRNA stability influences changes in total mRNA levels following transcriptional inhibition, 4sU pulse-chase combined with SLAM-seq was performed in K562 cells (Fig. 2A), with a concentration of 4sU that retained viability (Fig. S4. A) and proliferation (Fig. S4. B) throughout the experiment. This enabled the generation of estimated mRNA half-life (t_1/2_) parameters for 6580 genes (Fig. S4. C), including for short-lived genes such as *c-MYC* (t_1/2_ = 1.3 hours) (Fig. S4. D), which were concordant with published decay measurements obtained in K562 cells with conversion-based protocols (Fig. S4. E) (*20*, *29*). Functional annotation of genes according to their mRNA half-lives in combination with mRNA production rates, obtained using transient transcriptome (TT) -seq (*21*), revealed that leukaemogenic proteins, namely CR-TFs, were highly labile (Fig. 2B, S4. F). Similarly, high-expression of genes encoding only short-lived mRNAs correlated with poorer Acute Myeloid Leukaemia (AML) patient overall survival (Fig. S4. G-J). These data therefore suggest that fast mRNA kinetics are associated with oncogenes that are clinically relevant for AML.

**Figure 2.**
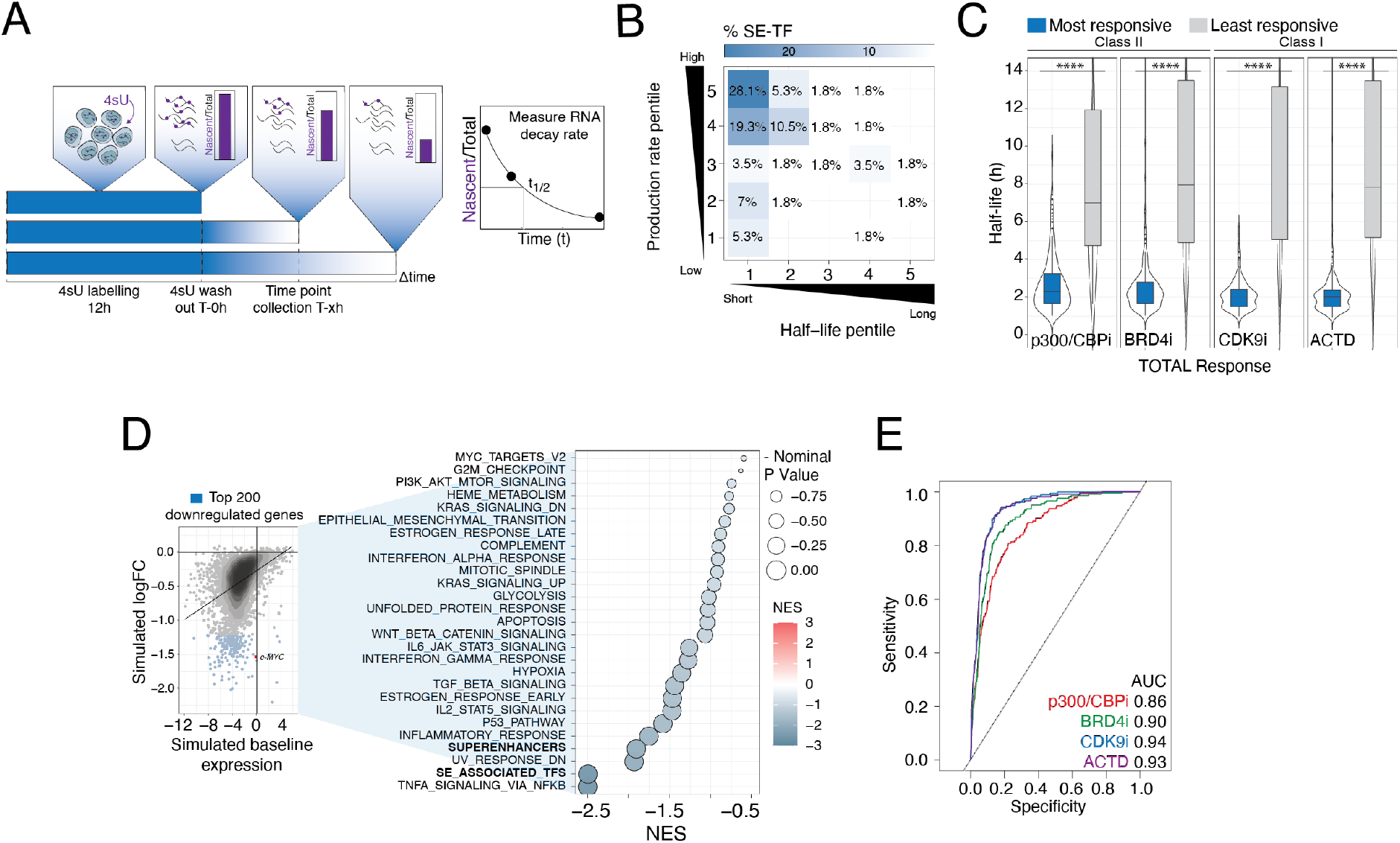
Gene intrinsic RNA decay properties shape the therapeutic response to transcriptional inhibitors. **(A)** Schematic of 4-thiouridine (4sU) pulse labelling and chase. T > C conversion rates were calculated for each time point, normalized to 0-h and fit with exponential decay functions derive half-life (t_1/2_). **(B)** Heatmap of percentage of CR TFs present within each mRNA decay and production pentiles from Fig. S4. F. **(C)** Boxplot of mRNA half-lives of genes most- and least-responsive to treatments indicated. **(D) (Right)** Simulation of total mRNA levels with complete transcription shutdown after two hours and **(left)** associated GSEA analysis of hallmark, SE and SE-associated TF gene sets using ranked z-scored logFC values. Signatures with Normalized Enrichment Scores (NES) < 0 shown. **(E)** Receiver Operator Characteristic (ROC) analysis of total logFC with treatments indicated and simulated total logFC after complete transcription shutdown for 2 hours. logFC values were binarized according to whether genes were most-responsive to treatments indicated. AUC: Area Under the Curve. Most-responsive: top 200 most significantly down-regulated genes (logFC < −0.5 and P Value < 0.05) using spike-in normalized reads. Least-responsive: 200 unaltered (−0.25 < logFC < 0.25 and P Value > 0.05) genes using spike-in normalized reads.

In light of our observation that transcripts within these oncogenic networks are rapidly turned over through a cycle of rapid production and decay (Fig. 2B, Fig. S4. F), we hypothesized that gene-intrinsic parameters are the causative factor driving selective total transcriptional changes to perturbation of the core-transcriptional machinery. Indeed, the most responsive genes to class I and II inhibitors had half-lives approximately 3-4 times shorter than those least responsive to RNAPII-targeting, and the median half-life of each category varied little between class I- or II-mediated transcriptional inhibition (Fig. 2C). Conversely, genes with short mRNA half-lives were significantly down-regulated in comparison to longer-lived gene groups on the total mRNA level with all compounds tested (Fig. S5. A) and were enriched for genes defined as the most responsive (Fig. S5. B). This was more evident with class I compounds, where >90% of most responsive genes were in the shortest t_1/2_ group (Fig. S5. B). In contrast, genes that were least responsive to RNAPII-targeting were over-represented for longer t_1/2_ gene categories (Fig. S5. C), indicating that transcripts with slow decay kinetics maintained total mRNA levels following acute transcriptional inhibition. In addition, mRNA production rates were predictive of total mRNA down-regulation following transcriptional inhibition only in the context of transcript half-lives, where genes with the highest mRNA production rates (pentile 5; 4.2e-02-6.7 FPKM h^-1^) and shortest half-lives (pentile 1; t_1/2_ 0.9-3.3 hours) were most strongly repressed following treatment with transcriptional inhibitors (Fig. S5. D). Moreover, the most-responsive genes (Fig. 1F) were enriched (25.5-37%) in the shortest lived and most highly produced subset (Fig. S5. D), where this was not observed for genes least-responsive (Fig. 1F, S5. E).

To quantitatively assess the impact of transcript half-lives to total mRNA abundance upon transcriptional inhibition *in silico*, we developed a mono-exponential model of steady state gene expression using only mRNA production and decay rates (Fig. S5. F). Production rates (*k_1_*) and decay rates (*k_2_*) were determined using TT-seq and SLAM-seq (Fig. 2A) measurements obtained in K562 cells, respectively. Predicted steady state mRNA levels significantly correlated with experimental measurements of baseline mRNA abundance (Fig. S5. G), indicating that the model was sufficiently accurate to estimate equilibrium mRNA levels with a variety of initial conditions and time intervals. *In silico* modelling of total mRNA abundance following two hours of complete abrogation of *de novo* mRNA synthesis (Fig. S5. F, H) and subsequent functional analysis, revealed significant negative enrichment of TFs, cytokine signalling, inflammatory and *c-MYC* target cancer hallmark pathways and SE-transcriptional programs, where oncogenes such *c-MYC* were amongst the most reduced (Fig. 2D, S5. I-J). This is in line with data obtained using total mRNA changes to small molecules targeting RNAPII (Fig. 1C, S2. G). Consistently, comparison of *in silico* transcription shutdown and experimental total mRNA measurements revealed highly similar and significant gene expression changes, albeit more strikingly with class I inhibitors (Fig. S5. K). In agreement with these results, short mRNA t_1/2_ genes were strongly predicted to be more responsive to RNAPII targeting, where predictive accuracy was higher with class I inhibitors (AUC 0.93-0.94) in comparison to class II (AUC 0.86-0.90) (Fig. 2E). This firstly highlights that any change in total mRNA levels following global perturbation of mRNA production with class I inhibitors is strongly determined by mRNA half-lives. Secondly, these data also suggest that changes in total mRNA levels of distinct genes affected by class II compounds stems from a combination of selective inhibition of *de novo* mRNA synthesis and gene intrinsic mRNA t_1/2_. Whilst reciprocal adjustments in mRNA synthesis and decay rates to maintain total cellular mRNA concentrations can occur, a phenomenon called ‘transcriptional buffering’, modelling of complete transcriptional shutdown using treatment-specific mRNA decay rates (Fig. S6. A-D) correlated significantly with experimentally measured total mRNA changes (Fig. S6. E). This indicates that transcriptional buffering does not negate the importance of mRNA decay rates in shaping the total mRNA response to the class of therapeutic transcriptional inhibitors.

Analysis of an independent SLAM-seq dataset from K562 cells (Fig. S7. A-D) (*19*) revealed that transcript half-life also defines the response to transcriptional disruption through small molecule inhibition of BRD4 or CDK9, as well as, acute protein degradation of c-MYC and BRD4 (Fig. S7. E-I). In addition, short-lived transcripts were most sensitive to inhibition of BCR-ABL1 signalling using nilotinib (Fig. S7. E-I). Concordant with our observations using K562 cells, the magnitude of nascent mRNA responses greatly exceeded those on the total mRNA level, indicating that transcript stability regulates this response, which has therapeutic implications. To extend our findings to another leukaemic model, we measured mRNA decay parameters (Fig. S8. A-B) and transcriptional responses to various small molecule transcriptional inhibitors (Fig. S8. C-D) in MLL-rearranged THP-1 cells. Moreover, we also included an independent and previously published SLAM-seq dataset assessing responses to transcriptional perturbation in THP-1 cells in our analysis (Fig. S8. C-D) (*30*). This revealed that similar to K562 cells, the total mRNA response in THP-1 cells to distinct BET, CDK9, CDK12/13 and CDK7 inhibitors was largely dependent on gene-intrinsic decay properties, where a much larger transcriptional response on the nascent mRNA level was observed (Fig. S8. E-H).

Taken together, these data demonstrate that mRNA decay parameters can isolate a group of responsive genes to all forms of transcription abrogation in manner that is completely agnostic to the chromatin landscape or the molecular consequences of the cofactor targeted.

### MAC-seq defines the role of mRNA decay properties to changes in gene expression with 73 epigenetic and transcriptional compounds

To assess the role of mRNA t_1/2_ parameters to gene expression responses across a wider range of clinically relevant inhibitors, we used Multiplexed Analysis of Cells sequencing (MAC-seq) to profile total mRNA changes in response to 73 small molecule inhibitors targeting epigenetic and transcriptional proteins in K562 cells, where previously tested drugs (Fig. 1A) were included as controls (Fig. 3A, S9. A). DGEA revealed a number of differentially expressed events (Fig. 3B), where only eight out the 73 compounds tested globally repressed transcription and significantly down-regulated > 1000 genes (Fig. 3C). This first group of inhibitors (class I) target cofactors described to exhibit broad chromatin specificities (*12*), including those able to perturb t-CDKs and the core RNAPII machinery (Pan-Tx) (Fig. 3C, S9. A). Moreover, a second group (class II) of an additional six drugs targeting BRD4, p300/CBP, histone deacetylases (HDACs), and protein phosphatases (PP), were able to selectively down-regulate a subset of least 200 genes, consistent with their role as regulating only a part of the genome (*12*) (Fig. 3C, S9. A). Despite technical and experimental differences between the MAC- and SLAM-seq protocols, MAC-seq total mRNA data recapitulated the previous segregation of the tested compounds into class I and II (Fig. 1A, 3C). Remaining compounds, defined as class III, that targeted histone demethylases (HDMe) and methyltransferases (HMe), DNA alkyl- (DNA AGT) and methyl-transferases (DNAMe), sirituin (SIRT), PARP, ELAVL1/HuR and topoisomerases (Topo) either repressed the expression of a minimal number of genes (< 200) or primarily up-regulated transcription (Fig. 3C). This indicates that gene-intrinsic mRNA stability most likely has minimal influence on the total transcriptome for class III compounds.

**Figure 3.**
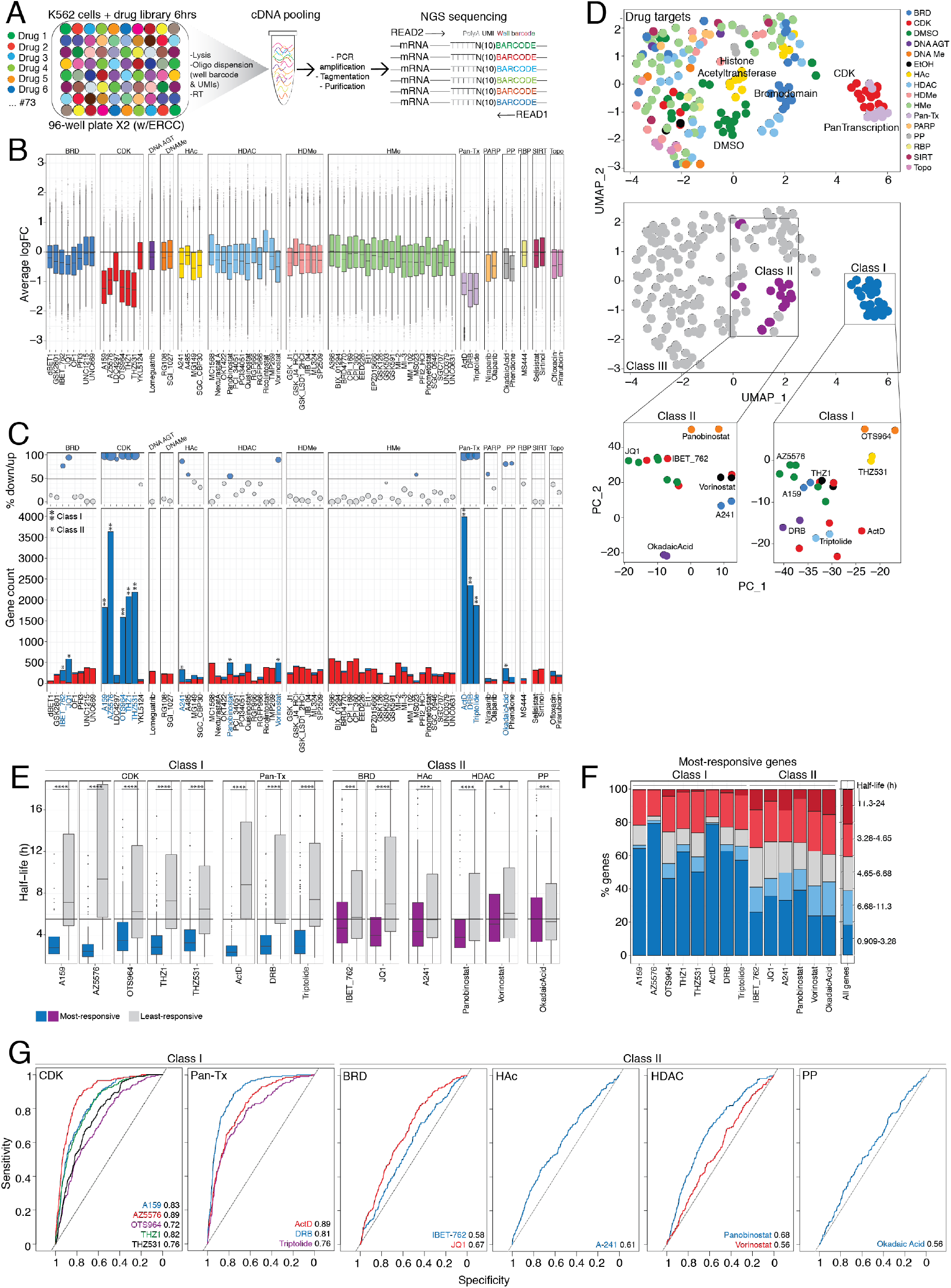
Gene intrinsic RNA decay properties shape the therapeutic response to several epigenetic and transcriptional inhibitors as defined by MAC-seq. **(A)** Schematic of MAC-seq experimental procedure. **(B)** Box plot of change in gene expression relative to DMSO/EtOH controls with inhibitors indicated. Inhibitors are grouped according to drug protein target family. **(C) (*Top*)** Percentage ratio or **(*bottom*)** absolute number of significantly up- and down-regulated genes. Inhibitors able to significantly down-regulate > 1000 genes are termed ‘Class I’ and by **. Inhibitors able to significantly down-regulate > 200 genes are termed ‘Class II’ and indicated by *. Remaining compounds as designated as class III. **(D)** Uniform Manifold Approximation and Projection (UMAP) of treatment conditions with **(*top*)** drug protein target family **(*middle*)** or pre-defined classes highlighted. **(*Bottom*)** PCA plot of drug classes **(right)** II and **(left)** I with drug names shown. **(E)** Boxplot of mRNA half-life of most- and least-responsive genes to class I and II inhibitors. **(F)** Bar chart representation of mRNA half-life pentiles in genes most-responsive to class I and II inhibitors. **(G)** Receiver Operator Characteristic (ROC) analysis of logFC with class I and II inhibitors and simulated total logFC after complete transcription shutdown for 6 hours. logFC values were binarized according to whether genes were most-responsive to treatments indicated. UMI: Unique Molecular Identifier. logFC: log2 fold change. BRD: Bromodomain. CDK: Cyclin Dependant Kinase. DNA AGT: DNA alkyl-transferase. DNAMe: DNA methyl-transferase. HAc: Histone acetylase. HDAC: Histone deacetylase. HDMe: Histone demethylase. HMe: Histone methyl-transferase. Pan-Tx: Pan-transcription. PP: Protein Phosphotase. RBP: RNA Binding Protein. SIRT: Sirtuin. Topo: Topoisomerase. Significantly up-regulated: P-Value < 0.05 and average logFC > 0.5. Significantly down-regulated: P-Value < 0.05 and average logFC < −0.5. logFC: log2 fold change relative to DMSO/EtOH. AUC: Area Under the Curve. Most-responsive: top 200 most significantly down-regulated genes (logFC < −0.5 and P Value < 0.05) using spike-in normalized reads. Least-responsive: 200 unaltered (−0.25 < logFC < 0.25 and P Value > 0.05) genes using spike-in normalized reads.

Uniform Manifold Approximation and Projection (UMAP) of drug treatments revealed that gene changes grouped according to drug treatment (Fig. S9. B, *middle*), drug protein target (Fig. S9. B, *bottom*) and family (Fig. 3D, *top*) and class (Fig. 3D, *middle*). Notably, despite all class I compounds forming an entirely distinct cluster (Fig. 3D, *middle*), more detailed analysis of this drug class in an isolated setting revealed separation between Pan-Tx, CDK9/7/12/13 and CDK11/12/13 inhibitors (Fig. 3D *bottom*, S9. A). This suggests that though class I compounds are able to broadly repress gene expression, they are mechanistically distinct and not equivalent. Inhibitors within the class II category however, with the exception of drugs targeting BET proteins, clustered separately (Fig. 3D *bottom*, S9. A), highlighting that they induce distinct and selective transcriptional responses in line with previously defined chromatin specificities (*12*).

Data from figure 2 demonstrated that mRNA decay is a critical determinant to changes in total gene expression following transcription inhibition with p300/CBPi, BRD4i, CDK9i and ACTD, so it was determined if these findings could be extended to other compounds able to globally (class I) or selectively (class II) repress gene expression as defined by MAC-seq (Fig. 3C). Genes most responsive (top 200 genes most significantly down-regulated) to class I and II inhibitors were significantly shorter lived in comparison to genes least responsive (P-Value > 0.05, −0.25 < logFC < 0.25; 200 genes closest to the mean logFC) to transcriptional targeting (Fig. 3E, S9. C). The difference in half-life between most and least responsive gene sets was most striking with class I inhibitors, where mRNA stability varied up to three-fold between the two categories (Fig. 3E, S9. C). Consistently, the most short-lived genes, as defined previously in Fig. 2, were enriched as genes most responsive to transcriptional targeting by class I and II inhibitors (Fig. 3F). This was more evident with class I compounds, which had > 40% of most-responsive genes within the most short-lived group (Fig. 3F). In agreement with these data, short-lived mRNAs were more significantly depleted amongst highly abundant transcripts in response to class I compounds in comparison to class II and III mRNAs (Fig. S9. E). In contrast, the most long-lived genes were enriched in genes least-responsive to targeting by most inhibitors within the two classes (Fig. S9. D) and as a consequence were found to be significantly over-represented across the most abundant mRNAs in responses to class I and II compounds compared to class III (Fig. S9. F). Furthermore, genes with short half-lives were highly predictive as genes amenable to t-CDK and general RNAPII inhibition using models of complete transcription shutdown (Fig. 3G), consistent with previous findings (Fig. 2E). Due to selective targeting observed on the nascent mRNA level and longer drug incubation times, predictive accuracy was modest with BRD4, p300/CBP, pan-HDAC and PP targeting (Fig. 3G). Taken together, these data demonstrate that the role of mRNA decay can be extended to several transcriptional and epigenetic inhibitors, where both selective and global down-regulation of gene expression is strongly dictated by gene intrinsic mRNA t_1/2_ parameters.

### c-MYC mRNA stabilization does not rescue biological and transcriptional effects of transcriptional targeting

A putative common effector mechanism of small molecules that inhibit mRNA production in leukaemia is repression of *c-MYC* (*13*). In K562 cells, *c-MYC* is necessary for driving the expression of a conserved set of gene targets encoding proteins necessary for nucleotide biosynthesis (*11*). Having determined that *c-MYC* itself and its target genes were short lived, we sought to assess whether altered *c-MYC* decay kinetics could modulate responses to transcriptional inhibition. As *c-MYC* stability is regulated by AU-rich elements (AREs), AUUUA and miRNA recognition motifs within its 3’UTR (*31*–*35*), CRISPR/Cas9 -mediated homology directed repair (HDR) in K562 cells was used to substitute the *c-MYC* endogenous 3’UTR for the 3’ region of the longer-lived gene *HPRT1* with a destabilized GFP (dsGFP) reporter, termed *‘c-MYC-HPRT1* 3’UTR’ (Fig. 4A). Designated as *‘c-MYC-*control 3’UTR’, dsGFP was knocked-in 5’ of the endogenous *c-MYC* 3’UTR sequence as a control cell line (Fig. 4A). Both cell models were validated by PCR (Fig. S10. A-F) and sequencing (Fig. S10. G-H) using primers complementary to various regions of the *c-MYC* locus. As our system genetically engineered the endogenous 3’UTR of *c-MYC*, its genomic location, in addition to *cis*- and *trans*-factors that impact its regulation remained unaffected. This therefore enables the direct assessment of altered mRNA stability to transcription targeting within the endogenous chromatin context.

**Figure 4.**
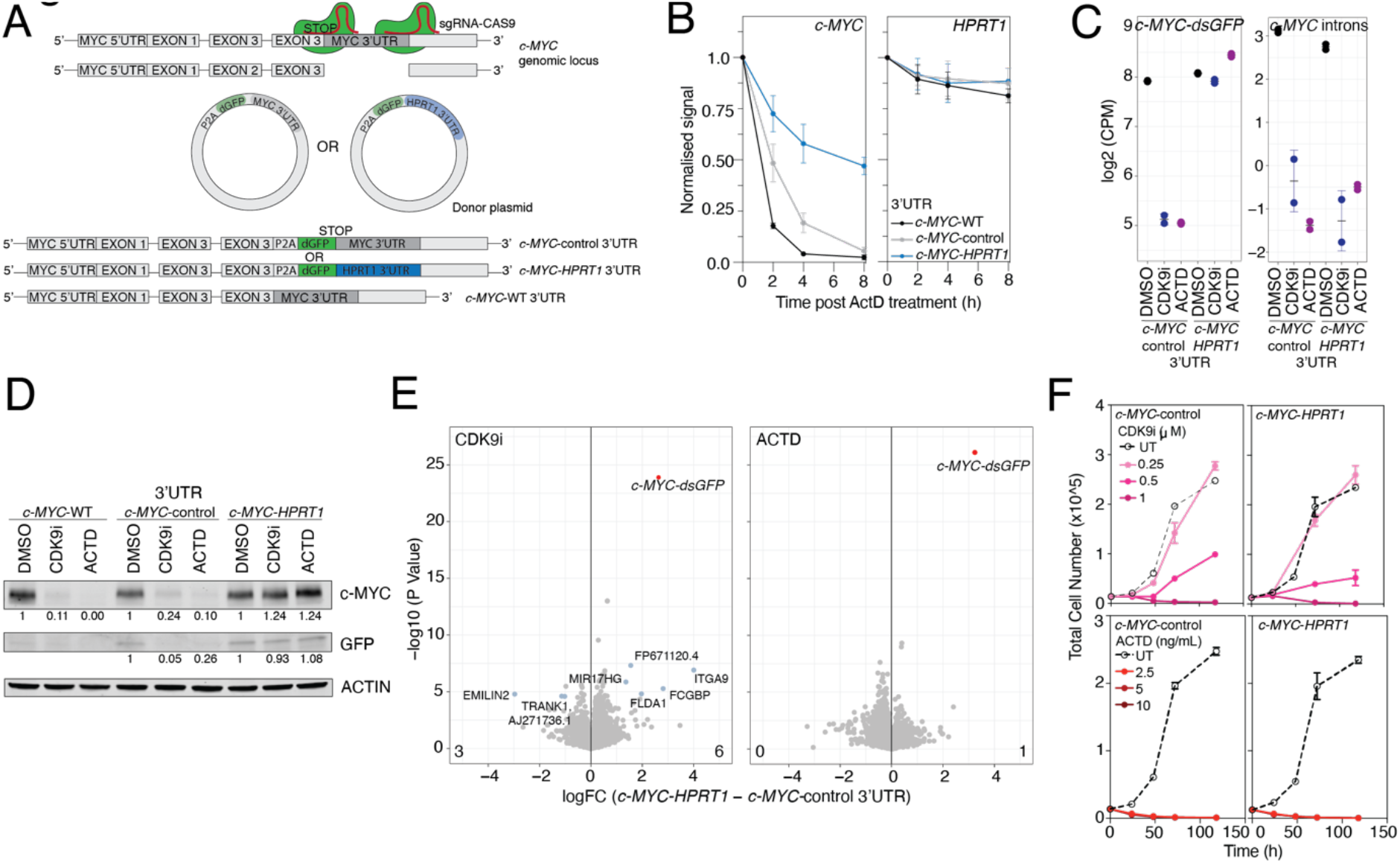
Increased *c-MYC* RNA stability does not rescue biological and transcriptional effects of transcriptional inhibitors. **(A)** Schematic of CRISPR/Cas9 -homology directed repair (HDR) used to endogenously swap the *c-MYC* 3’ untranslated region (3’UTR) for the *HPRT1* 3’UTR in the K562 cell line. **(B)** Normalized expression of **(left)***c-MYC* and **(right)***HPRT1* transcripts following the addition of ACTD at indicated time points as measured by quantitative real time PCR (qRT-PCR). Values are mean with error bars representing standard deviation (sd) of three biological replicates from three separate single cell clones. **(C) (Left)** Total gene expression of reads mapping across *c-MYC* and *dsGFP* sequences with indicated treatments and cell lines after 6 hours. **(Right)***c-MYC* intron expression with indicated treatments and cell lines after 6 hours. **(D)** Western blot of **(top)** c-MYC, **(middle)** GFP and **(bottom)** ACTIN protein with indicated treatments and cell lines after six hours. Values indicated are protein levels normalized to ACTIN and DMSO controls. **(E)** Scatter plot of significance and difference in total gene expression with **(left)** CDK9i and **(right)** ACTD treatment between *c-MYC*-control and *-HPRT1* 3’UTR cell lines. **(F)** Total cell number over time of indicated cell lines treated with increasing concentrations of **(top)** CDK9i and **(bottom)** ACTD. logFC: log2 fold change.

Assessment of mRNA stabilities demonstrated that knock-in of the *HPRT1* 3’UTR increased the half-life of the chimeric *c-MYC* transcript by approximately eight hours following treatment with ACTD, representing a four-fold increase over the half-life of the endogenous and entopic *c-MYC* transcripts (Fig. 4B, S11. A-B). Moreover, *c-MYC* transcript stabilisation did not alter baseline *c-MYC* total mRNA abundance (Fig. S11. C), in contrast to nascent mRNA levels, where it was significantly lower in comparison to the control (Fig. S11. D).

To further investigate the response of stabilized *c-MYC* to RNAPII targeting, c*-MYC*-control and *-HPRT1* 3’UTR cell lines were treated with the class I compounds AZ-5576 (CDK9i) or ACTD for six hours and whole transcriptome RNA-Seq was performed. Consistent with previous findings (Fig. 1, 3), DGEA of exon and intron reads revealed a global decrease in total and nascent gene expression, respectively (Fig. S11. E-G). Whilst d*e novo c-MYC* synthesis was significantly down-regulated irrespective of *c-MYC* mRNA half-life (Fig. 4C *right), c-MYC-dsGFP* total mRNA (Fig. 4C *left*) and protein abundance (Fig. 4D, S11. H-K) revealed that *c-MYC* stabilization rendered it less sensitive to targeting by either compound. The resistance to RNAPII targeting on the total mRNA level was *c-MYC* specific, as *c-MYC* was the most significantly altered gene with each treatment (Fig. 4E) and clustering of total mRNA changes highlighted that conditions grouped according to inhibitor and not cell line (Fig. S11. L). Consistently, correlation analysis of all DEGs in c*-MYC-*control and *-HPRT1* 3’UTR cell lines with CDK9i and ACTD revealed that changes were highly similar and significant (Pearson’s correlation coefficient 0.96-0.97, P-Value < 2.2e-16) (Fig. S11. M).

Remarkably, increasing *c-MYC* mRNA half-life through swapping of the endogenous *c-MYC* 3’UTR with that of *HPRT1* did not rescue the expression of previously defined *c-MYC* target genes (Fig. S11. N-O) (*11*). Although nascent *c-MYC* target gene expression was significantly less down-regulated with CDK9i treatment in the *c-MYC-HPRT1* 3’UTR cell line (Fig. S10. N), it was not sufficient to prevent reductions in total mRNA (Fig. S10. N). In line with these data, the proliferation and survival between K562 c*-MYC*-control and -*HPRT1* 3’UTR cell lines was not altered upon CDK9i and ACTD treatment *in vitro* (Fig. 4F).

Taken together, these data suggest that class I inhibitors can mediate anti-leukemic effects even when expression of a key target gene, *c-MYC*, is maintained due to prolonged mRNA half-life. This challenges the notion that loss of *c-MYC* expression is necessary for the biological and molecular consequences of perturbing nascent mRNA synthesis using agents such as AZ-5576 or ACTD. Therefore, the observation that *c-MYC* repression is the effector mechanism upon inhibition of transcriptional proteins in leukaemia is not due to specific targeting of *c-MYC* SEs and CR-TFs, but rather a consequence of short transcript half-lives. Gene-intrinsic mRNA decay properties are thus a key determinant in establishing gene responsiveness, such as that of *c-MYC*, to compounds that target RNAPII driven gene expression, and without which, selective responses would not be found with most transcriptional compounds.

### Selective modulation of mRNA decay through ELAVL1 targeting

Having established that the therapeutic response to transcriptional inhibition is shaped by mRNA decay kinetics, we next investigated whether mRNAs with long half-lives and consequently, not amenable to perturbation, could be sensitised to RNAPII targeting in leukaemia. We therefore re-analyzed our previously published genome-wide CRISPR-Cas9 knockout screening datasets performed in AML cell lines treated with CDK9i, with a focus on RBPs due to their ability to bind mRNAs at specific sequence motifs to affect mRNA post-transcriptional processing (*36*).

The analysis revealed that knockout of 16 RBPs sensitized both THP-1 and MV4;11 AML cell lines to CDK9i (Fig. 5A, S12. A). This included the RBP ELAVL1 (Fig. 5A, S12. A-B), which has been reported to stabilize oncogenic transcripts in AML and promote Leukemic Stem Cell (LSC) self-renewal and survival (*37*, *38*). THP-1 cells with knockout of *ELAVL1* (Fig. 5B) were sensitized to the anti-leukemic effects of AZ-5576 (Fig. 5C-D, S12. C), thereby independently validating the results from the genome-wide screens. Given the proposed oncogenic role of *ELAVL1* (*37*, *38*) it was unsurprising to observe that *ELAVL1* knockout also resulted in a proliferative disadvantage in the absence of any CDK9i treatment (Fig. 5D). This was concomitant with a modest decrease in global total mRNA levels (Fig. S12. D-E) and an enrichment of interferon signalling pathways (Fig. S12. F), in line with previous reports in THP-1 cells and validating our experimental model (*39*).

**Figure 5.**
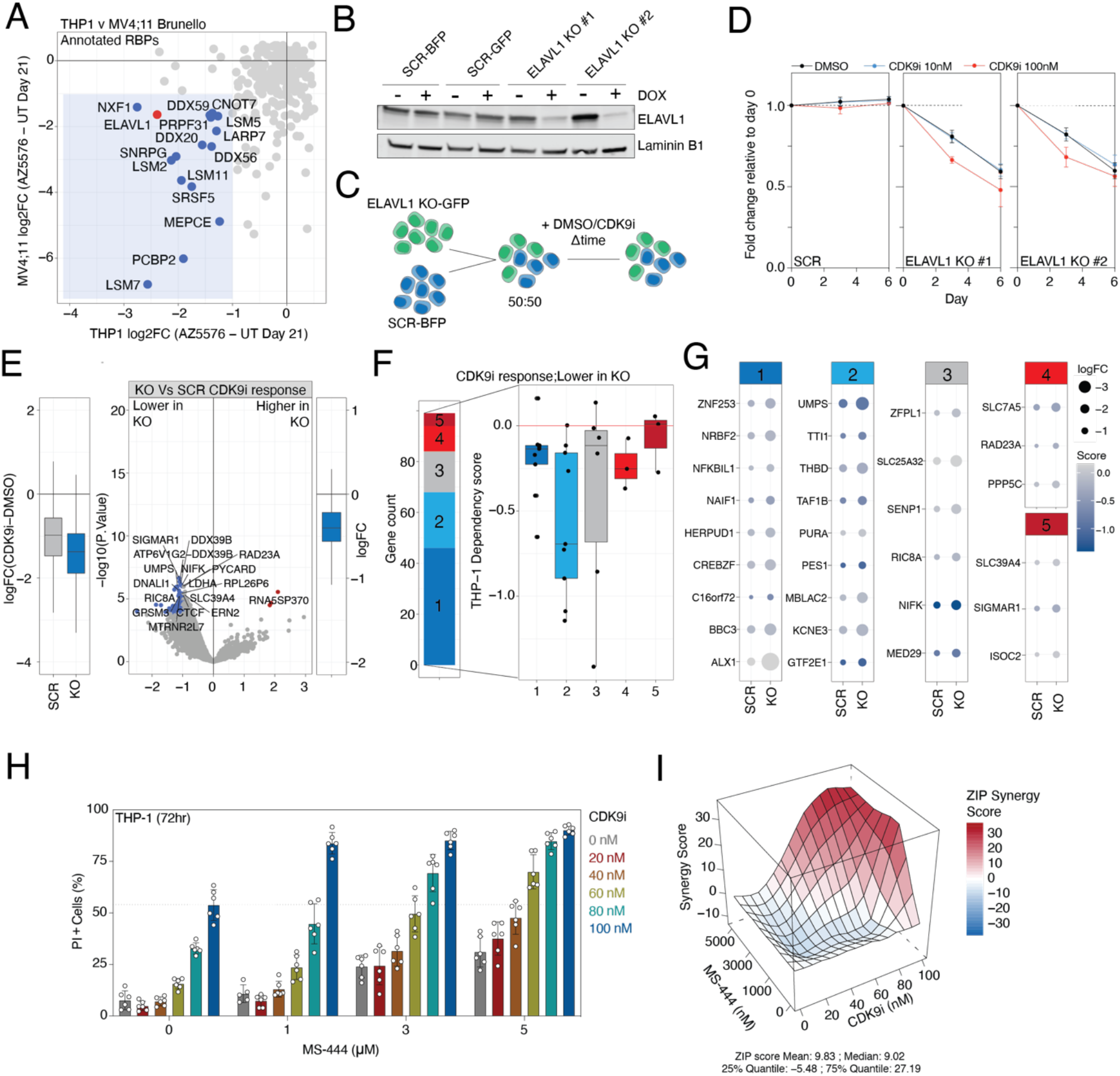
Dual targeting of CDK9 and ELAVL1 is synergistic in AML. **(A)** Scatter plot of change in sgRNA read counts in CDK9i compared to untreated (UT) CRISPR-Cas9 genome-wide screening groups after 21 days of passaging in THP-1 and MV4;11 AML cell lines. Annotated RBPs that are down-regulated (logFC < −1) in both THP-1 and MV;11 cell lines indicated in blue and red (ELAVL1). Data is obtained from (*72*). **(B)** Western blot of **(top)** ELAVL1 and **(bottom)** Laminin B1 protein with indicated treatments and THP-1 cell lines after four days. **(C)** Simplified schematic of competition assay experimental design. THP-1 cells expressing Cas9 and non-targeting control sgRNA (SCR) and sgRNAs targeting ELAVL1 (ELAVL1 knockout) are mixed in a 1:1 ratio and passaged in the presence of DMSO or CDK9i. **(D)** Fold change relative to day 0 of THP1 ELAVL1 knockout cells in competition with SCR cells as described in (C) at time points and treatments indicated. **(E) (Left)** Boxplot of spike-in normalized total mRNA reads with CDK9i relative to DMSO in THP-1 SCR and ELAVL1 KO cells. **(Right)** Scatter plot of significance and difference in spike-in normalized total gene expression of CDK9i response in ELAVL1 knockout relative to SCR cells. Significantly down- and up-regulated genes are highlighted in blue and red, respectively. **(F) (Left)** Stacked bar chart of previously determined transcript half-life pentiles in THP-1 cells across mRNAs sensitized to CDK9i upon ELAVL1 knockout (lower in KO; logFC < −1 and adjusted P Value < 0.01) and **(Right)** associated boxplot and **(G)** dot plot of THP-1 gene dependency scores from the Broad Institute Cancer Dependancy Map. LogFC values in each cell line are indicated. **(H)** Bar plot of THP-1 cell death (propidium iodide (PI) positive) in response to CDK9i and MS-444 at concentrations indicated following 72 hours. **(I)** ZIP synergy scores derived from (H). Representative of first replicate. logFC: log2 fold change. SCR: non-targeting sgRNA control. KO: knockout. Significantly down-regulated: logFC < −1 and adjusted P Value < 0.01. Significantly up-regulated: logFC > 1 and adjusted P Value < 0.01.

To interrogate the mechanism of CDK9i sensitization mediated by ELAVL1 knockout, THP1 ELAVL1 wild-type and knockout cells were treated with CDK9i for six hours and 3’UTR RNA-seq was performed. Consistent with our findings (Fig. 1, 3), DGEA of total read counts revealed a global decrease in total gene expression with CDK9i across both *ELAVL1* wild-type and knockout cell lines, and the magnitude of global transcript down-regulation was modestly greater in ELAVL1 knockout cells (Fig. 5E, S12. D, G). Whilst the CDK9i response between cell lines was similar as demonstrated by co-clustering of samples (Fig. S12. H) and correlation analysis (Pearson’s correlation coefficient 0.917, P-Value < 2.2e-16) (Fig. S12. I), further investigation revealed 57 genes that were significantly more down-regulated (logFC < −1 and adjusted P Value < 0.01) upon ELAVL1 genetic depletion (Fig. 5E, S12. J). As ELAVL1 has been described to regulate mRNA stabilities, the half-lives of the 57 transcripts sensitised to CDK9i following *ELAVL1* knockout were assessed. Using previous transcript decay measurements in THP-1 cells (Fig. S7. G), it was evident that whilst the 20% most short-lived mRNAs were enriched in transcripts sensitised in *ELAVL1* knockout upon CDK9i, there was a subset of transcripts with longer half-lives (Fig. 5F *left*). Moreover, incorporation of THP-1 dependency scores obtained from the Broad Institute Depmap Portal demonstrated that sensitised transcripts with longer half-lives were highly essential, such as *NIFK* and *MED29* (Fig. 5F *right*, G). Indeed, both *NIFK* and *MED29* have been associated with oncogenic characteristics in a variety of cancer types (*40*, *41*). Collectively, our findings highlight that leukaemia cell line sensation to CDK9i following *ELAVL1* knockout may be associated with the perturbation of essential mRNAs with long half-lives, which would otherwise be less amenable to transcriptional perturbation in *ELAVL1* wild-type cells.

To interrogate the therapeutic potential of ELAVL1 targeting in the context of CDK9i in leukaemia, THP-1 cells were treated with CDK9i in combination with the small molecule MS-444, a potent inhibitor of ELAVL1 (*42*). Importantly, MS-444 has been demonstrated to exhibit single agent efficacy in models of leukameia *in vivo* (*38*). Assessment of cell death revealed that the drug combination resulted in greater cytotoxicity than each inhibitor alone and was highly synergistic (Fig. 5H-I, S12. K). These data are in line with our genetic ELAVL1 knockout experiments and suggest that the therapeutic perturbation of the mRNA decay and transcriptional machinery is an actionable and novel combination therapy for the treatment of AML.

## Discussion

RNAPII-driven transcription has been therapeutically targeted across solid and haematological malignancies with biological and therapeutic effects proposed to occur through selective perturbation of oncogenic gene networks (*1*, *26*). The current literature suggests that the discrete targeting of selected genes following transcriptional perturbation is mechanistically linked to association with enhancers, SEs and disproportionate occupancy of critical chromatin co-factors, such as BRD4 and TFs, at promotors; however, the role of mRNA stability has not been extensively assessed in this context.

Here, we demonstrate that selective and broad targeting of RNAPII-driven transcription by class I and II therapeutic inhibitors, respectively, results in discrete alterations in total mRNA abundance that are defined by transcript decay rates (Fig. 6). Despite our analyses being completely agnostic to the chromatin landscape or the molecular consequences of the core-transcriptional component targeted, they were able to effectively isolate a group of responsive genes to all forms of transcriptional perturbation. Indeed, both class I and II inhibitors reduce the total mRNA levels of genes with short transcript half-lives, several of which are CR TFs, associated with oncogenic signalling pathways or promotor proximal to SEs. Although some studies indicate that these oncogenic networks are particularly sensitive due to reduction of the CR TFs within SE and promotor regions, whether down-regulation of TFs and SE-driven transcription were causally linked remained unclear (*26*, *43*, *44*). Our data indicates that targeting of SE-driven transcription is only possible when the associated gene has short-lived mRNA, and therefore implicates gene-intrinsic mRNA decay parameters in the selective perturbation of SE oncogenic programs. This further highlights that the selectivity of these therapies is very likely over-estimated and suggests that oncogenic networks driven by SEs and short-lived and highly produced CR TFs are intrinsically sensitized to any form of abrogation of *de novo* mRNA synthesis, thus challenging the notion that selective targeting is required to specifically disrupt oncogenic transcription. In contrast, genes with stable mRNAs are largely refractory to transcriptional targeting when assessing total mRNA levels and are functionally related to cellular housekeeping roles (*18*, *45*). This may prove a challenge for the use of transcriptional inhibitors as a therapeutic intervention for leukaemias that exhibit dysregulated metabolic activity or a heightened dependence on metabolic pathways.

**Figure 6.**
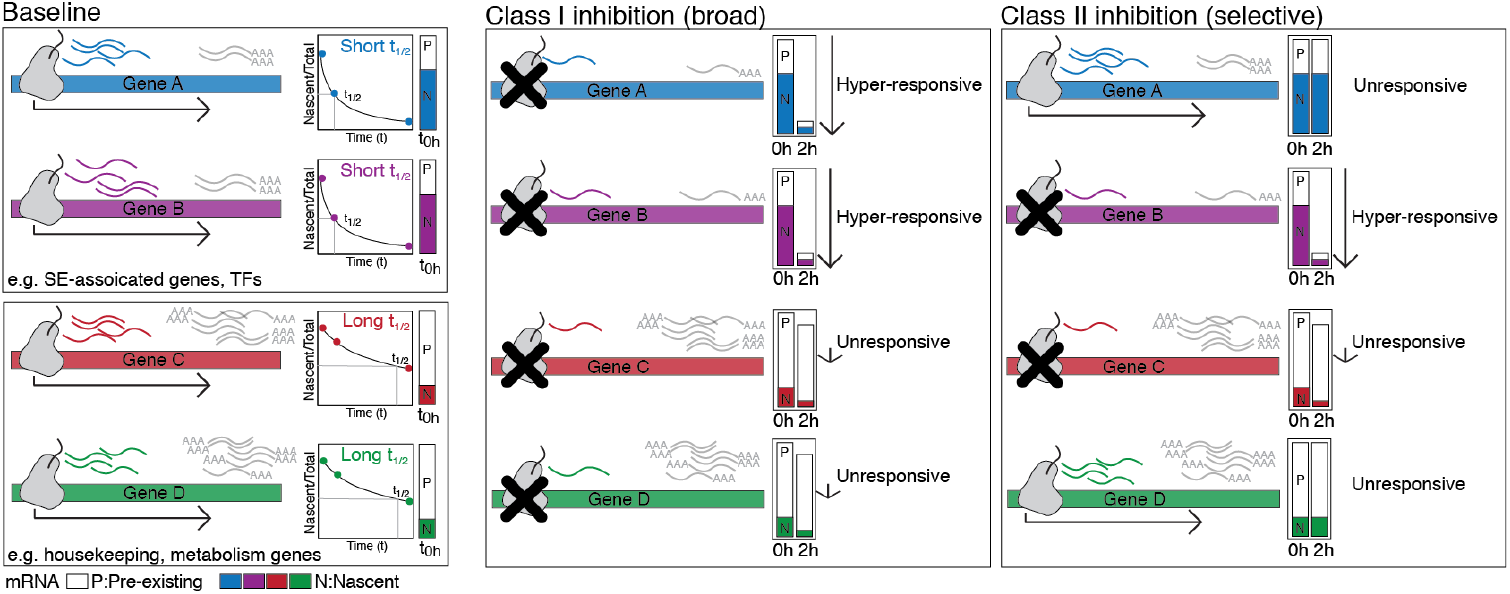
Simplified schematic of transcriptional responses of genes with short and long RNA half-lives with class I (selective) and class II (broad) inhibitors.

Interestingly, alterations in mRNA decay frequently occur in cancer, including leukaemias, in either a gene-specific manner or through mutations in mRNA complexes to thus affect global mRNA turnover (*35*, *46*–*49*). For example, stabilization and the resultant increase in steady state expression of the *PD-L1* transcript frequently occurs in haematological and stomach cancers from mutations that disrupt the *PD-L1* 3’UTR, and in *in vivo* models of lymphoma, leads to immune evasion and decreased survival (*35*). Similarly, c-MYC 3’UTR truncations caused by chromosomal translocations in human T-cell leukaemias (TCL) have been described to increase c-MYC expression via transcript stabilisation (*50*, *51*). In addition to gene-specific truncations of 3’UTR sequences in cancer, mRNA stability can be broadly affected via genetic alterations that impact the mRNA decay machinery and RBPs. This includes loss of function mutations in the Carbon Catabolite Repression—Negative On TATA-less (CCR4-NOT) Transcription subunit 3 (CNOT3) subunit of the CCR4-NOT deadenylase complex in T-cell Acute Lymphoblastic Leukaemia (T-ALL) (*47*) and the 3’-5’ exoribonuclease of the exosome complex, DIS3 in MM and RUNX1 -mutated AML (*48*, *49*). Moreover, widespread alternative polyadenylation (APA) in a variety of cancer types has been demonstrated to shorten 3’UTRs and increase transcript stability and protein levels (*46*, *52*). Whilst the mechanism of increased APA in cancers is poorly understood, there is evidence that down-regulation APA factors such as CFIm25 and PCF11 may contribute increased APA in certain malignancies (*53*). Based on our observations that long-lived transcripts are resistant to transcriptional inhibition, targeting stabilized oncogenic mRNA in the context of deregulated decay machinery may prove unsuccessful with class I and II therapeutic compounds.

Due to the lack of on-target inhibitors, therapeutic inhibition of mRNA decay pathways in the oncology setting remains largely unexplored. The widely utilised chemotherapeutic nucleoside analog 5-flurouracil (5-FU), initially characterised to perturb the DNA synthesis regulator thymidylate synthase, has been also reported to perturb the exosome subunit EXOSC10 (*54*, *55*). Moreover, small molecule inhibition of the scavenger mRNA de-capping enzyme DCPS using RG3039 in pre-clinical AML models affected pre-mRNA splicing, rather than cytoplasmic decay pathways to elicit cytotoxic effects (*56*). An alternative approach to alter transcript decay in cancer for therapeutic benefit has been to target RBPs that promote oncogenic mRNA stability. In the leukaemia setting, recent work has demonstrated that the ELAVL1 inhibitor MS-444 destabilises the mitochondrial import protein TOMM34 and hampers LSC-driven leukaemic reconstitution *in vivo* (*38*). In line with these findings, we observe that genetic knockout and therapeutic inhibition of ELAVL1 limits THP-1 cell line growth. Targeting mRNA decay pathways directly or through RBPs is of high clinical significance, particularly because long-lived transcripts not amenable to transcriptional inhibition encode for proteins that regulate cellular homeostasis and metabolism. Indeed, aberrant metabolism is recognised as a cancer hallmark and is central to leukaemia cell survival and proliferation (*57*, *58*). This is supported by pre-clinical studies in leukaemia models showing that small molecule inhibition of various metabolic pathways as a highly promising avenue for leukaemia therapy (*58*). Efforts to destabilise long-lived mRNAs may therefore be an alternative approach to perturb oncogenic metabolism.

Having observed that a large number of mRNAs are not sensitive to transcriptional perturbation due to their long half-lives, we demonstrated that combination therapy with a transcript destabilizing agent such as MS-444, provides a novel opportunity to target genes that were otherwise impervious to RNAPII targeting. Whilst we demonstrated that 57 transcripts were significantly sensitised with ELAVL1 knockout upon CDK9i treatment, only a minority possessed long half-lives. As knockout of ELAVL1 has been demonstrated to alter mRNA decay rates, our half-life measurements, derived from THP-1 cells with wild-type ELAVL1, may under-estimate the number of transcripts with short mRNA decay kinetics (*59*).

Overall, this study highlights the importance of mRNA decay parameters in governing total gene expression levels in response to selective and global inhibitors of RNAPII -driven transcription, and provides novel mechanistic insight that can be leveraged for therapeutic benefit. Moreover, our work shifts the paradigm that selective responses observed in these contexts are largely driven by selective abrogation of mRNA synthesis as a result of particular genomic determinants but rather demonstrates that these largely result from gene-intrinsic mRNA properties.

## Methods

### Cell lines and culture

K562 CML parental and HDR-edited and THP-1 AML parental and inducible knockout cells were cultured in Roswell Park Memorial Institute (RPMI) medium (Thermo Fisher Scientific, Waltham, MA, USA, 11875093) containing 10% (v/v) heat-inactivated fetal bovine serum (HI-FBS; Thermo Fisher Scientific, 10099), penicillin (100 U/ml), streptomycin (100 μg/ml) (Thermo Fisher Scientific, 15140122) and 2 mM GlutaMAX (Thermo Fisher Scientific, 35050061) at 37°C and 5% carbon dioxide. HEK293T cells were cultured in Dulbecco’s Modified Eagle’s Medium (DMEM; Thermo Fisher Scientific, 11995073) containing 10% (v/v) heat-inactivated fetal bovine serum (HI-FBS; Thermo Fisher Scientific, 10099), penicillin (100 U/ml) and streptomycin (100 μg/ml) (Thermo Fisher Scientific, 15140122) at 37°C and 10% carbon dioxide. Drosophila melanogaster S2 cells were cultured Schneider’s *Drosophila* medium (Thermo Fisher Scientific, 21720) supplemented with 10% HI-FBS, penicillin (100 U/ml), streptomycin (100 μg/ml), and 2 mM GlutaMAX at room temperature and atmospheric carbon dioxide. Cell lines for all assays were seeded at 50-70% confluency the day prior unless otherwise indicated.

### Compounds

See supplementary tables 1 and 2.

### Generation of ELAVL1 inducible knockout cell lines

Independent sgRNAs targeting ELAVL1 (*69*) and non-targeting sgRNA controls (SCR; supplementary table 3) flanked with BsmBI overhangs were ligated into the vector FgH1tUTG-GFP (Addgene Plasmid #70183) using 1uL BsmBI (New England Biolabs, R0739) in NEBuffer 3.1 (New England Biolabs, B7203S). This was similarly performed for SCR sgRNA cloned into the vector FgH1tUTG-BFP (*47*). Cloned FgH1tUTG vectors, pMDLg/pRRE (Addgene Plasmid #12251), pRSV-Rev (Addgene Plasmid #12253) and pVSV.G (Addgene Plasmid #12259) third generation lentiviral vectors were transiently transfected into HEK293T cells using polyethylenimine (PEI; 1μg/mL; 25kD linear; BioScientific Pty Ltd, 23966). Following 48 hours, viral supernatant was harvested and filtered using a 0.45μM filter. Viral supernatant was added THP-1 cells constitutively expressing S. *pyogenes* Cas9 (FUCas9-Cherry vector; Addgene Plasmid #70812) prior to transduction using sequa-brene (4μg/mL; Sigma Aldrich, S2667-1VL) and centrifugation at 500g for 4 minutes at room temperature (*47*). Expanded cells were sorted for populations expressing cherry and GFP or BFP using Becton Dickinson (BD) Fussion 3 or 5 sorters. sgRNA expression was induced using doxycycline (1μg/mL; Sigma Aldrich, D9891). All experiments were performed four days following doxycycline induction unless otherwise indicated.

### CRISPR HDR generation of MYC transcript stable clones

Parental K562 cells (5e+05) were washed in Phosphate Buffered Saline (PBS) twice and resuspended in Cell Line Nucleofector solution SF (16.4uL) with Supplement (3.6uL) (SF Cell Line 4D-nucleofector X Kit, Lonza, V4XC-2032). Alt-R SpCas9 nuclease (100pmol, Integrated DNA Technologies, 1074182), single guide RNAs (sgRNAs) targeting the c-MYC stop codon and 3’ end of its 3’UTR (300pmol, supplementary table 3) and pUC57 or pUC57-Mini donor plasmid (1000ng; GenScript Gene Sythesis) containing recombinant sequences for dGFP, P2A cleavable peptide and HPRT1 or c-MYC 3’UTRs, respectively (supplementary table 4), were incubated together for 20 minutes at room temperature, prior to being placed on ice. Ribonucleoprotein (RNP) complex (5uL) was added to the cell suspension (20uL), 20uL of which was subsequently transferred to 16-well Nucleocuvette Strip and electroporated using the 4D-Nuceleofector X unit (program FF120, Lonza, AAF-1002X). Warmed culture medium (100uL) was added to cell suspension and incubated at 37°C and 5% carbon dioxide for 10 minutes. Cell suspension was then transferred to a 24-well cell culture plate containing culture medium (1mL) and IDT Alt-R HDR electroporation enhancer (20uM; Integrated DNA Technologies, 1081073) and incubated at 37°C and 5% carbon dioxide for 24 hours, after which cells were washed twice with fresh culture medium and cultured at 37°C and 5% carbon dioxide. After expansion, cells were sorted and enriched for GFP positivity three successive times, followed by isolation of single cells into 96-well culture plates using Becton Dickinson (BD) FACSAria Fusion 3 or 5 Cell Sorters. Three clones with successful knock in of each donor vector were identified with KAPA HiFi (Roche, 7958935001) using isolated genomic DNA (DNeasy Blood & Tissue Kits (Qiagen, 69506) and primers designed outside or within plasmid homology arms (supplementary table 5). PCR fragments were subsequently separated using agarose gel electrophoresis (see below). Knock-in sequence was validated using Sanger sequencing of PCR fragments detailed above at the Australian Genome Research Facility (AGRF). Total-RNA sequencing was performed using a single representative clone of each knock-in.

### Propidium Iodide, 4’,6-diamidino-2-phenylindole and Cell Trace Violet staining

For cell viability and proliferation studies using propidium iodide (PI) and/or Cell Trace Violet staining, respectively, cells (1 x10^7^) were centrifuged (500g at 4°C for 4 minutes), resuspended in PBS supplemented with 0.1% (w/v) BSA and stained with 5μM CTV dye (Thermo Fisher Scientific, C34557) in a 37°C water bath for 10 or 20 minutes. Five volumes of ice-cold culture medium was added to cell suspension to quench unbound dye. Cells were then centrifuged (500g at 4°C for 4 minutes), resuspended in PBS supplemented with 2% (v/v) HI-FBS, sorted for a narrow peak of CTV-positivity on BD FACSAria Fusion 3 or 5 Cell Sorters. Following treatment, CTV-stained or unsustained cells were resuspended in PBS containing 1μg/mL PI (Sigma Aldrich, P4170).

For cell viability studies using 4’,6-diamidino-2-phenylindole (DAPI), cells were treated with AZ-5576 (CDK9i) for 72 hours, centrifuged (500g at 4°C for 4 minutes) and resuspended in PBS containing 1 μg/mL DAPI (Sigma Aldrich, D9542-100MG). Analysis of flow cytometric data was preformed using the BD Fortessa X20 and FlowJo v10 software (Ashland). ZIP synergy scores were determined and visualised using synergyfinder (v3.0.1) on Rstudio (v4.0.2).

### Total cell number quantification

Absolute cell number was determined with the addition of 1×10^4^ calibration beads directly to cells prior to analysis. 0.2 μM Propidium iodide (PI) was also added with the beads to identify dead cells by exclusion. Ratio of live cells to beads was measured by flow cytometry to determine the absolute live cell number in cell culture.

### Intracellular staining of c-MYC

Intracellular staining of Myc was performed as previously described (*60*). Briefly, cells were harvested at the timepoints indicated and were immediately resuspended in fixation buffer (0.5% paraformaldehyde, 0.2% Tween-20 and 0.1% bovine serum albumin in PBS) at room temperature, for at least 24 hours until staining was performed. Fixed cells were stained with either anti-Myc (clone D84C12, Cell Signalling Technology) or a rabbit IgG isotype-matched control antibody (clone D1AE, Cell Signalling Technology) before staining with an anti-rabbit IgG conjugated to Alexa Fluor 647. Staining of all fixed samples within one experiment was performed at the same time.

### Cell competition assays

Four days after induction of sgRNA using doxycycline (1μg/mL; Sigma Aldrich, D9891), THP-1 Cas9 cells expressing FgH1tUTG-GFP constructs were mixed at a 1:1 ratio with THP-1 Cas9 cells expressing a SCR sgRNA in the FgH1tUTG-BFP backbone. Cells were cultured in the presence of DMSO or AZ’5576 (CDK9i; 10nM and 100nM). The BD LSR Fortessa Flow Cytometer was used to assess the relative proportions of GFP- and BFP-positive cells following mixing (time point 0) and at time points indicated.

### Agarose gel electrophoresis and gel imaging

Blue/orange loading dye 6X (Promega, G1881) was added to PCR fragments, which were subsequently separated using 1% agarose gels prepared with molecular grade agarose (Bioline, BIO-41025), Tris base-acetic acid-EDTA (TAE) 1X solution and SYBR Safe DNA Gel Stain (Life Technologies, S33102). Agarose gels were imaged on the GelDoc XR+ Imager (BioRad) using ImageLab Software (BioRad).

### Quantitative PCR and analysis

Cells (1e+06 per time point) were incubated with 1ug/mL ACTD at 37°C and 5% carbon dioxide, harvested 0-, 2-, 4- and 8-hours post-treatment, centrifuged (1400rpm at 4°C for 4 minutes), washed in ice-cold PBS and resuspended in 300uL TRIzol (Thermo Fisher Scientific, 15596026). RNA was extracted from lysates using the Direct-zol RNA MiniPrep Kit (Zymo Research, R2052) and complimentary DNA (cDNA) was synthesised (from 1ug RNA) using the Applied Biosystems High Capacity cDNA Reverse Transcription Kit (Thermo Fisher Scientific, 4368814). Quantitative PCR (qPCR) was performed using cDNA, 0.25uM forward and reverse oligo primers (see supplementary table 6) and SensiFAST SYBR Hi-ROX Kit (Bioline, BIO-92005) in 384-well plates with the LightCycler 480 Instrument II (Roche, 05015243001). Threshold cycles for each reaction were analysed using the ΔΔ*C*_t_ method normalising to GAPDH as the housekeeping gene.

### SDS-polyacrylamide gel electrophoresis and western blotting

Cells (1e+06) were washed in PBS, lysed in Laemelli Buffer (60 mM tris-HCl (pH 6.8), 10% (v/v) glycerol, and 2% (w/v) SDS) and incubated at 95°C for 10 minutes. Lysate protein concentration was determined using the Pierce BCA Protein Assay Kit (Thermo Fisher Scientific, 23225) and absorbance at 562nm wavelength was measured on the iMark Microplate Absorbance reader (BioRad) using MicroPlate Manager Software (BioRad). 20 X sample buffer (100 %β-mercaptoethanol, and bromophenol blue) was added to lysates and were subsequently incubated for an additional 5 minutes. Lysates were separated using Mini-PROTEAN TGX 4 to 15% gradient gels (25 mM tris, 192 mM glycine, and 0.1% (w/v) SDS; Bio-Rad, 4561086) and transferred at 4°C to either Immobilon-FL or Immunoblon-P (IPVH00010) polyvinylidene fluoride membranes (Merck, IPFL00010) (1.5 hours; 250 mA, 25 mM tris, 192 mM glycine, and 5% (v/v) methanol).

Immobilon-FL membranes were dried for 1 hour at room temperature, washed in ultrapure water, 100% methanol, Tris buffered saline (TBS) in the listed order and blocked using Odyssey blocking buffer (Li-COR, 927-40000). They were then incubated in primary antibodies (supplementary table 7) diluted in Odyssey blocking buffer supplemented with 0.2% (v/v) Tween 20 (Sigma-Aldrich, P9416) overnight at 4°C, washed three times with TBS containing 0.1% (v/v) Tween 20 and incubated with IRDye-conjugated secondary antibodies (supplementary table 7) diluted in Odyssey blocking buffer supplemented with 0.2% (v/v) Tween 20 and 10% (v/v) sodium dodecyl sulfate (SDS) for 1 hour at room temperature. Immobilon-FL membranes were washed in PBS and protein was visualised and quantified using the Odyssey CLx and Image Studio Lite software 2 (Li-COR).

Immunoblon-P membranes were blocked in TBS supplemented with 5% (w/v) skim milk powder and Tween 20, incubated with primary antibodies (supplementary table 7) diluted in the same solution at 4°C overnight, washed three times with TBS containing 0.1% (v/v) Tween 20 and incubated with horseradish peroxidase-conjugated secondary antibodies (supplementary table 7) at room temperature for 1 hour. Protein was subsequently visualised using Amersham ECL Plus (GE Healthcare, RPN2132) and Super RX film (Fujifilm, 03G01).

### Total RNA-sequencing

Cells (1e+06^6^ per treatment condition) were centrifuged (1400rpm at 4°C for 4 minutes), washed in ice-cold PBS and resuspended in 350μl of Buffer RLT Plus supplemented with 1% (v/v) β-mercaptoethanol from the RNeasy Plus Mini Kit (Qiagen, 74134). RNA was extracted using the same kit according to manufacturer’s instructions and quality was assessed using the Agilent 2200 TapeStation System (Agilent, G2964AA) with RNA ScreenTape (Agilent, 5067-5576) and Sample Buffer (Agilent, 5067-5577). S2 spike-in material (5%) was added to RNA. Sequencing libraries were prepared using NEBNext Ultra II Directional RNA Library Prep Kit for Illumina, where ribodepletion was performed using the NEBNext rRNA Depletion Kit (New England BioLabs, E6310). Paired-end 75 base pair (bp) reads were sequenced using the NextSeq 500 (Illumina).

### 3’UTR-sequencing

Cells were treated with DMSO or AZ-5576 (CDK9i; 100nM) for 6 hours at 37°C and 5% carbon dioxide. Following incubation, cells were harvested, centrifuged (500g at 4°C for 4 minutes), washed in ice-cold PBS and resuspended in 600μL TRIzol. RNA was extracted using the Directzol RNA Miniprep Kit, where an additional DNase I digestion step was performed (Zymogen; R2052). Sequencing libraries were prepared using the QuantSeq 3’mRNA-seq Library Prep Kit FWD for Illumina (Lexogen, Vienna, Austria) and sequenced as single-end 75 base pair reads using the NextSeq 500 (Illumina).

### SLAM-sequencing

Protocol was adapted from (*18*). K562 and THP-1 cells (1e+06 per treatment) were first pre-treated with small molecule inhibitors for a total time of 2 hours to pre-establish protein-target inhibition. Newly synthesized RNA in K562 cells was then labelled using 100 μM 4-sU in the final 1 hour of treatment at 37°C and 5% carbon dioxide. Cells were washed in ice-cold PBS and resuspended in 300μL TRIzol. For direct measurement of RNA half-lives, newly synthesized RNA was labelled by incubating cells in 30μM 4-sU for 12 hours at 37°C and 5% carbon dioxide, whereby culture medium was exchanged every 3 hours for the duration of the pulse. For the uridine chase, cells were centrifuged (1400rpm at 4°C for 4 minutes), washed in sterile ice-cold PBS twice and resuspended in pre-warmed (37°C) culture medium containing 3mM uridine (Sigma Aldrich, U6381). At 0-,1-, 2-, 4-, 8-, 12-hours for K562 cells and 0-, 0.3-, 0.6-, 1-, 2-. 4-, 8-hours for THP-1 cells post the uridine chase, cells were harvested, centrifuged (1400rpm at 4°C for 4 minutes) and resuspended in 300μL TRIzol. For treatment-specific RNA decay rates, cells were centrifuged (1400rpm at 4°C for 4 minutes), washed in sterile ice-cold PBS twice and resuspended in pre-warmed (37°C) culture medium containing 3mM uridine (Sigma Aldrich, U6381) and either 1μM DMSO, JQ1 or AZ-5576. At 0-, 2-, 4- and 8-hours post uridine chase/drug addition, cells for each treatment condition were harvested, centrifuged (1400rpm at 4°C for 4 minutes) and resuspended in 300μL TRIzol. All SLAM-seq experiments included a non-4-sU labelled control unless otherwise stated. To extract RNA, one-fifth volume of chloroform was added to TRIzol lysates, followed by shaking, incubation at room temperature for 2 minutes and centrifugation (16000g at 4°C for 15 minutes). The aqueous phase was isolated and RNA was precipitated using DTT (10mM), 100% isopropanol (1 volume) and GlycolBlue co-precipitant (15ug, Ambion, AM9515), incubated at room temperature for 10 minutes and centrifuged (16000g at 4°C for 20 minutes). Supernatant was removed, RNA pellets were washed in 75% (v/v) ethanol and DTT (100μM) and centrifuged (7500g at room temperature for 5 minutes). Supernatant was removed and RNA pellets dried for 10 minutes prior to reconstitution in Ultrapure DNAse/RNAse-free distilled water (Thermo Fisher Scientific, 10977023) supplemented with DTT (1mM) and incubation at 55°C for 10 minutes. Thiol-containing bases were reduced by incubating RNA (10μg) with IAA (10mM, 50 mM NaPO4 pH 8.05 and 50% (v/v) DMSO) in a final volume of 50μL for 15 minutes at 55°C. Reaction was stopped by quenching IAA with DTT (20μM). RNA was precipitated using 3M NaOAc pH 5.2 (5μL), 100% ethanol (125μL) and GlycolBlue co-precipitant, incubated at −80°C for 30 minutes and centrifuged (16000g at 4°C for 30 minutes). Supernatant was removed and RNA pellet was washed in 75% (v/v) ethanol and centrifuged (16000g at 4°C for 10 minutes) twice, dried at room temperature for 10 minutes and reconstituted in Ultrapure DNAse/RNAse-free distilled water. RNA clean-up was performed by incubating RNA solution in 2 volumes of AMPure XP Beads (Beckman Coulter, A63881) for 2 minutes at room temperature. Beads were washed in 80% (v/v) ethanol twice, dried at room temperature for 3 minutes and resuspended in Ultrapure DNAse/RNAse-free distilled water. Eluate was collected and RNA quality and concentration was assessed using the Agilent 2200 TapeStation System (Agilent, G2964AA) with RNA ScreenTape (Agilent, 5067-5576) and Sample Buffer (Agilent, 5067-5577). S2 spike-in material (5%) was added to RNA. Sequencing libraries were prepared using the QuantSeq 3’mRNA-seq Library Prep Kit FWD for Illumina (Lexogen, Vienna, Austria) and sequenced as single-end 75 base pair reads using the NextSeq 500 (Illumina).

### TT-sequencing

Protocol is adapted from (*21*). Cells (5e+07 per treatment condition) were incubated with 1mM 4sU for either 5 or 15 minutes at 37°C and 5% carbon dioxide, centrifuged (1400rpm at 4°C for 5 minutes) and resuspended in TRIzol (5mL). One-fifth volume of chloroform was added to lysates, followed by shaking, incubation at room temperature for 2 minutes and centrifugation (13000g at 4°C for 10 minutes). The aqueous phase was isolated and RNA was precipitated using 100% isopropanol (1 volume), incubated at room temperature for 10 minutes and centrifuged (13000rpm at 4°C for 10 minutes). Supernatant was removed and RNA pellet was washed in 70% (v/v) ethanol, reconstituted in Ultrapure DNAse/RNAse-free distilled water (100μL) and denatured at 65°C for 10 minutes. S2 spike-in (15μg) was added to RNA (150μg), material was adjusted to a final volume 100μL and sonicated in microTUBE AFA Fiber screw cap tubes (6 mm × 16 mm, Covaris, 520096) using the Covaris S220 Focused-ultrasonicator at a maximum power for 15 seconds. Thiol-specific biotinylation of RNA was performed in a final volume of 1.5mL by incubation with 10 mM tris (pH 7.4), 1 mM EDTA, 20% (v/v) dimethylformamide (200 μg/ml; Sigma-Aldrich, 227056), and 300 μg of EZ-Link HPDP-Biotin (Thermo Fisher Scientific, 21341) for 1.5 hours at room temperature. An equal volume of chloroform was added to reaction, followed by shaking, incubation at room temperature for 2 minutes and centrifugation (1400rpm at 4°C for 5 minutes). Aqueous phase was isolated and an equal volume of chloroform was added, followed by shaking, incubation at room temperature for 2 minutes and centrifugation (1400rpm at 4°C for 5 minutes). This step was repeated an additional one time. Aqueous phase was isolated and RNA was precipitated using 5M NaCl (10% volume) and 100% isopropanol (1 volume) and centrifugation (20000g at 4°C for 20 minutes). RNA pellets were washed in 75% (v/v) ethanol, reconstituted in Ultrapure DNAse/RNAse-free distilled water (100μL) and denatured at 65°C for 10 minutes. Biotinylated RNA was separated from the total RNA pool by incubation with streptavidin beads (μMACs Streptavidin Kit, Miltenyi Biotec, Bergisch Gladbach, Germany, 130-074-101) at room temperature for 15 min with constant rotation. μMACs columns pre-equilibrated with room temperature wash buffer (100 mM tris-HCl (pH 7.4), 10 mM EDTA, 1 M NaCl, and 0.1% (v/v) Tween 20) were used to bind streptavidin beads, which were then washed using 900μL wash buffer that was heated to 65 °C or at room temperature, 5 times each. Biotinylated RNA was eluted into 700μL Buffer RLT from the RNeasy MinElute Cleanup Kit (Qiagen, 74204) through two additions of 100 mM DTT (100μl) 3 min apart, and then isolated using the same kit according to manufacturer’s instructions. RNA concentration was quantified using the Agilent 2200 TapeStation System (Agilent, G2964AA) with High Sensitivity RNA ScreenTape (Agilent, 5067-5579) and Sample Buffer (Agilent, 5067-5580).

Sequencing libraries were prepared with the NEBNext Ultra II Directional RNA Library Prep Kit for Illumina (without additional fragmentation), where ribodepletion was performed using the NEBNext rRNA Depletion Kit (New England BioLabs, E6310). Single-end 75 base pair reads were sequenced using the NextSeq 500 (Illumina).

### MAC-seq

Cells (5e+04 per well) in a final volume of 100μL in a 96-well plate format were incubated with transcriptional and epigenetic inhibitors in technical duplicate for 6 hours (supplementary table 8) at 37°C and 5% carbon dioxide. 5e+03 cells from each well were aliquoted into a separate 96-well plate, washed in ice-cold PBS twice and centrifuged (1400rpm at 4°C for 4minutes). Supernatant was removed and cell pellets were frozen at −80°C. Library preparation is adapted from (*61*). In detail, 17 μl lysis buffer were added into each well of a 96-well plate containing cell pellets and incubated at room temperature for 15 min under agitation (900 rpm). 12.5 μl of cell lysate were transferred into each well of a new 96-well plate previously prepared with 1 μl of 10 nM well-specific RT MAC-seq primer and 7.5 μl RT mix; the RT mix contains a TSO primer and external ERCC RNAs. The mixture was incubated for 2 hours at 42 C to create well-barcoded full length cDNA and then all the wells of a plate were combined into a single tube. Concentration and clean-up was done via column purification (DNA Clean and Concentrator^™^-100, Zymo Research) and RNAClean XP beads (Beckman Coulter) and each plate eluted in 22 μl nuclease free water. The purified cDNA was pre-amplified with KAPA HiFi HotStart ReadyMix (Roche) and MAC-seq PreAmp PCR primer and the quality checked on a D5000 Screentape (TapeStation, Agilent). One barcoded library was prepared per plate using TD buffer and TDE1 enzyme (Illumina) for tagmentation and KAPA HiFi HotStart Ready Mix (Roche) and custom primers (MAC-seq P5 PCR and MAC-seq Indexing Mix) for amplification. Libraries were purified with DNA Ampure XP beads (Beckman Coulter), quality checked on a DNA1000 tape (TapeStation, Agilent) and quantity verified by qPCR. Two indexed libraries were sequenced on a NextSeq 500 instrument (Illumina) using a custom sequencing primer (MAC-seq Read primer) and a High Output Kit v2.5 75 Cycles (Illumina) with paired-end configuration (25 base pairs for read 1 and 50 base pairs for read 2).

### SLAM-seq analysis

Single-end reads were demultiplexed using bcl2fastq (v2.17.1.14) and resulting FASTQ files were trimmed for adapter sequences using Trim_galore (v0.6.5) with a stringency overlap of 3bp. Trimmed FASTQ files were processed with SLAM-dunk (v0.4.3), enabling multi-mapper reconciliation and using a threshold of at least 2 T > C conversions to mark a read as converted. The no 4sU control sample was processed first to find single nucleotide polymorphisms (SNPs), which was then used to filter the subsequent samples. ‘bedtools merge’ was then used to merge the 3’ UTRs of each Ensembl transcript by gene (v77 identifiers for hg38) for use in the SLAM-dunk counting step. The Broad Institute GSEA software was used to perform GSEA (*62*). Differential gene testing was performed on counts normalized to library sizes scaled to external S2 spike-ins and filtered for lowly expressed genes using edgeR (v3.32.1) on Rstudio (v4.0.2). SLAM sequencing tcount files from (*30*) (GEO accession GSE138210) and (*11*) (GEO accession GSE111463) were downloaded and differential gene testing was performed as described above.

### TT-seq analysis

Single-end reads were demultiplexed using bcl2fastq (v2.17.1.14) and resulting FASTQ files were aligned to the genome using STAR (v2.7), which were then summarized using featureCounts in Subread (v2.0.1): counting reads with a minimum Mapping Quality Score (MQS) of 7 in the union of all transcript isoforms of each Ensembl gene. Only genes with at least 10 reads per million across at least two samples within an experiment were retained for further analysis. In order to compare expression levels between samples, raw read counts for each gene were converted to reads per million using adjusted library sizes calculated with edgeR’s TMM implementation on the spike-in read counts.

### Total RNA-seq analysis

Paired-end reads were demultiplexed using bcl2fastq (v2.17.1.14) and resulting FASTQ files were quality checked using fastqc (v0.11.6), trimmed 15 bp from the 5’ end to remove primer bias and filtered for quality and length using cutadapt (v2.1; -u 15 -U 15 -q 15 --pair-filter any --minimum-length 50). Trimmed reads were mapped to GRCh38/hg38 and BDGP6/dm6 genomes using hisat2 (v2.1.0) with paired read settings and resulting SAM files were converted to BAM files using samtools (v1.9; view), which were then sorted and indexed. Reads mapping to either exonic or intronic genomic intervals were counted using a combined hg38/dm6/GFP GTF file with FeatureCounts from the subread package (v2.0.0; featureCounts -O -M -T 32 -p -s 2). Differential gene testing was performed on counts normalized to library sizes scaled to external S2 spike-ins and filtered for lowly expressed genes using edgeR (v3.32.1) on Rstudio (v4.0.2). Gene Ontology analysis was performed using the ToppGene suite (*63*–*65*).

### 3’UTR-seq analysis

Single-end reads were demultiplexed using bcl2fastq (v2.17.1.14) and resulting FASTQ files were quality checked using fastqc (v0.11.6), trimmed 12 bp from the 5’ end to remove primer bias, trimmed for polyA stretches, and filtered for quality and length using cutadapt (v2.1; -u 12 -q 10 -m 20 -a AAAAAAAAAA). Trimmed reads were mapped to GRCh37/hg19 and BDGP R5/dm3 genomes using hisat2 (v2.1.0) using standard single-end settings and resulting SAM files were converted to BAM files using samtools (v1.9; view), which were then sorted (sort) and indexed (index). Reads mapping to exonic genomic intervals were counted using a hg19 GTF file and dm3 SAF file with FeatureCounts from the subread package (v2.0.0; featureCounts -O -M -F GTF -T 16 -s 0). Differential gene testing was performed on counts normalized to library sizes scaled to external S2 spike-ins and filtered for lowly expressed genes using edgeR (v3.32.1) on Rstudio (v4.0.2). The Broad Institute GSEA software was used to perform GSEA (*62*).

### MAC-seq analysis

Paired-end reads were demultiplexed using bcl2fastq (v2.17.1.14) and resulting FASTQ files were quality checked using fastqc (v0.11.6) and read 2 (R2) was trimmed 15 bp from the 5’ end to remove primer bias using cutadapt (v2.1; -u 15). R2 FASTQ files of paired-end reads were demultiplexed according to well barcodes (supplementary table 9) and filtered for PCR duplicates using Unique Molecular Identifiers (UMIs), both present in read 1 (R1) using the scruff (*66*) R (v4.0.2) package dumultiplex function (bcStart = 1, bcStop = 10, bcEdit = 0, umiStart = 11, umiStop = 20, keep = 35, minQual = 20, yieldReads = 1e+06). R2 FASTQ files were then mapped to the GRCh37/hg19 genome and ERCC sequences using alignRsubread (unique = FALSE, nBestLocations = 1, format = “BAM”) and resulting BAM files were used to count unique R2 reads mapping to exonic genomic intervals and ERCC sequences using a combined hg19/ERCC GTF file with countUMI (umiEdit = 0, format = “BAM”, cellPerWell = 1. Both functions are from the scruff R package. Gene expression counts were normalized to library size and reads mapping to ERCC spike-ins using the RUVseq R package (v1.24.1). Subsequent count processing was performed using the Seurat R package (v3.2.1) (*67*), where lowly expressed genes were filtered and counts were normalized for latent variables including plate, well row and column, using the SCTransform function. SCTransformed scaled gene RNA expression values were then used for PCA, where shared-nearest-neighbours (SNN) network was calculated using the top 10 Principal Components with the FindNeighbours function using default parameters. Drug-treatment clusters were subsequently identified with the Louvain algorithm using a resolution parameter of 2. Uniform Manifold Approximation and Projection (UMAP) values were also calculated using the top 10 Principal Components with the RunUMAP function using default parameters. Differential gene testing relative to treatment controls (DMSO or EtOH) was performed using a hurdle model (MAST) with un-scaled gene RNA expression counts, plate and column numbers as latent variables and a logFC threshold of 0 with the FindMarkers Function. Area Under the Curve (AUC) scores for each drug treatment and gene lists indicated was calculated using all expressed genes with the R AUCell Package (v0.10.0).

### ChIP-seq analysis and super-enhancer identification

H3K27ac ChIP-seq FASTQ files (SRR1957037, SRR1957038, GEO Accession GSM1652918) were downloaded using sratoolkit (v2.9.0, fastq-dump --gzip --split-files), quality checked using fastqc (v0.11.6) and mapped to the GRCh37/hg19 genome using bowtie2 (v2.3.4.1) with paired end read settings. Resulting SAM files were converted to BAM files using samtools (v1.9; view), which were then sorted and indexed. Potential PCR duplicates were filtered using the MarkDuplicates function from picard (v2.6.0; REMOVE_DUPLICATES= true) and BAM files were converted to TDF files using igvtools (v2.3.95; count -z 5 -w 10). H3K27ac peaks were called relative to input using macs (v2.1.1; callpeak -f BAMPE -g hs -q 0.01 -call-summits –cutoff-analysis) and peaks within hg19 ENCODE blacklist regions (https://www.encodeproject.org/files/ENCFF356LFX/) were removed using bedtools (v2.27.1) intersect function. Super-enhancer calling was performed using Ranking Ordering of Super-Enhancer (ROSE2) (v1.0.5; -c INPUT -g hg19 -t 2000 -s 12500) (*68*) and were annotated to genes according to “TOP GENE.”

### Genome-wide CRISPR Cas9 screening analysis

Trimmed 20 base pair reads from survival screens performed in THP-1 and MV4;11 Cas9 cells transduced with the Brunello genome-wide sgRNA library and cultured in either DMSO or AZ-5576 (CDK9i; 150nM) for 21 days from (*69*), were used for sgRNA counting with the MAGeCK algorithm, where read counts from treatment replicates were pooled (v0.5.6; mageck count). The MAGeCK algorithm test subcommand (mageck test) was then used to determine genes negatively selected in CDK9i relative to DMSO treatment groups. Visualization of screen data was performed in R (v4.0.2) using ggplot2 (v3.1.1) and ggrepel (v0.8.1) packages. Annotated RBPs were identified using data from (*70*).

### Coltron analysis

The coltron algorithm (v1.0.2) (*27*) was used to identify SE-associated TFs and regulatory networks using ROSE2-determined super-enhancer peaks and H3K27ac BAM signal with default parameters.

### TCGA survival analysis

SEM (RNA-Seq by Expectation Maximization) (*71*) scaled expression values (transcripts per million) for TCGA tumour samples were downloaded from the GDAC Firehose website. Entrez gene IDs were mapped to HGNC gene symbols using the biomaRt R package (version 2.42.0) and collapsed to unique values per gene symbol by selecting the most variable entrez ID among all samples for each gene symbol. Primary samples from the TCGA LAML cohort were selected using the TCGA biolinks R package (version 2.14.0) and were matched with overall survival (OS) endpoints from the TCGA Pan-Cancer Clinical Data Resource. Gene signature scores were calculated using the singscore R package,then used to fit Kaplan-Meier and Cox regression models to the OS endpoints using the survival R package (version 3.1-8).

### Determination of RNA synthesis and decay rates

Solving the first order differential equation in Fig. 2F yields an exponential function with parameters for the synthesis rate (k_1_), decay rate (k_2_) and initial quantity of RNA (M_0_):

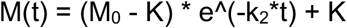

where K = k_1_/k_2_ is the quantity at equilibrium.

To determine decay rates (k_2_) for each gene, precision-weighted nonlinear regression was used to fit an exponential curve to the decreasing quantity of SLAM-seq labelled reads measured post-washout of 4sU.

Assuming no synthesis of labelled reads post-washout (k_1_ = 0) leaves one parameter for the starting RNA concentration, which was set to the initial data point at T = 0, and one for the decay rate, which was fit using MINPACK’s (v1.2-1) Levenberg-Marquardt implementation in R.

Precision weights for the fit were estimated using local regression. The standard deviation between technical replicates for a given mean expression level in the baseline SLAM-seq dataset was modelled with R’s ‘loess’ implementation, and then applied to calculate weights for the other samples during fitting.

Synthesis rates were determined by dividing the TT-seq adjusted reads per million for each gene by the length and sampling time, producing a scaled FPKM h^-1^ averaged across replicates.

### Predictions

Predictions were made with the model using the fitted rates and differing initial conditions, then compared against a separate dataset of total mRNA measurements at T = 2, 6h.

Parameters set for simulated results:

| **Simulated Result** | **Decay rate ($k_1$)** | **Synth. rate ($k_2$)** | **Initial qty ($M_0$)** |
| --- | --- | --- | --- |
| Simulated total transcriptional shutdown | Fitted decay rate under drug | 0 | Baseline equilibrium |

## Acknowledgments

We thank members of the Victorian Centre for Functional Genomics, Peter MacCallum Centre Molecular Genomics Core, Daniel Ho from Novartis and the other authors for their help during DRUG-seq protocol implementation and Compounds Australia at Griffith University for their provision of specialized compound management and logistics research services to the project.

## Author contributions

I.T. performed experiments, next generation sequencing analysis, data interpretation and wrote the manuscript. B.F. and A.T.P performed next generation sequencing analysis and computational modelling. Z.F. designed sgRNA and plasmid templates for CRISPR-HDR. S.G., D.Y., G.M.A. and K.J.S. optimized and performed DRUG-seq. I.Y.K. performed Myc intracellular staining and total cell number assays and analysis and was supervised by E.D.H. S.B. performed combination therapy experiments. M.Z. performed TCGA analysis. S.J.V. and R.W.J. performed data interpretation, study supervision and co-wrote the manuscript. All co-authors proof-read and edited the manuscript.

## References

1. J. E. Bradner, D. Hnisz, R. A. Young, Transcriptional Addiction in Cancer. Cell. 168, 629–643 (2017).

2. M. J. Bywater, R. B. Pearson, G. A. Mcarthur, R. D. Hannan, Dysregulation of the basal RNA polymerase transcription apparatus in cancer. Nat. Rev. Cancer. 13 (2013), doi:10.1038/nrc3496.

3. E. Smith, C. Lin, A. Shilatifard, The super elongation complex (SEC) and MLL in development and disease. Genes Dev. 25, 661–672 (2011).

4. C. Villicaña, G. Cruz, M. Zurita, The basal transcription machinery as a target for cancer therapy (2014).

5. N. Laham-karam, G. P. Pinto, A. Poso, P. Kokkonen, Transcription and Translation Inhibitors in Cancer Treatment. 8, 1–24 (2020).

6. S. M. Abedin, C. S. Boddy, H. G. Munshi, BET inhibitors in the treatment of hematologic malignancies: Current insights and future prospects. Onco. Targets. Ther. (2016), doi:10.2147/OTT.S100515.

7. G. E. Winter, D. L. Buckley, J. Paulk, J. M. Roberts, A. Souza, S. Dhe-paganon, J. E. Bradner, Phthalimide conjugation as a strategy for in vivo target protein degradation. Science (80-.). 348, 1376–1382 (2015).

8. L. M. Lasko, C. G. Jakob, P. Rohinton, W. Qiu, D. Montgomery, E. L. Digiammarino, T. M. Hansen, R. M. Risi, R. Frey, V. Manaves, B. Shaw, M. Algire, P. Hessler, L. T. Lam, T. Uziel, E. Faivre, D. Ferguson, F. G. Buchanan, R. L. Martin, M. Torrent, G. G. Chiang, K. Karukurichi, J. W. Langston, B. T. Weinert, C. Choudhary, P. De Vries, A. F. Kluge, M. A. Patane, J. H. Van Drie, C. Wang, D. Mcelligott, E. A. Kesicki, R. Marmorstein, C. Sun, P. A. Cole, S. H. Rosenberg, M. R. Michaelides, A. Lai, K. D. Bromberg, Discovery of a selective catalytic p300/CBP inhibitor that targets lineage-specific tumours. Nature. 550, 128–132 (2017).

9. O. Bensaude, Inhibiting eukaryotic transcription: Which compound to choose? How to evaluate its activity? Transcription. 2, 103–108 (2011).

10. R. D. Martin, T. E. Hébert, J. C. Tanny, Therapeutic targeting of the general RNA polymerase II transcription machinery. Int. J. Mol. Sci. 21 (2020), doi:10.3390/ijms21093354.

11. M. Muhar, A. Ebert, T. Neumann, C. Umkehrer, J. Jude, C. Wieshofer, P. Rescheneder, J. J. Lipp, V. A. Herzog, B. Reichholf, D. A. Cisneros, T. Hoffmann, M. F. Schlapansky, P. Bhat, A. Von Haeseler, T. Köcher, A. C. Obenauf, J. Popow, S. L. Ameres, J. Zuber, SLAM-seq defines direct gene-regulatory functions of the BRD4-MYC axis. Science (80-.). 360, 800–805 (2018).

12. C. Neumayr, V. Haberle, L. Serebreni, K. Karner, O. Hendy, A. Boija, J. E. Henninger, C. H. Li, K. Stejskal, G. Lin, K. Bergauer, M. Pagani, M. Rath, K. Mechtler, C. D. Arnold, A. Stark, Differential cofactor dependencies define distinct types of human enhancers. Nature. 606, 406–413 (2022).

13. J. Zuber, J. Shi, E. Wang, A. R. Rappaport, H. Herrmann, E. A. Sison, D. Magoon, J. Qi, K. Blatt, M. Wunderlich, M. J. Taylor, C. Johns, A. Chicas, J. C. Mulloy, S. C. Kogan, P. Brown, P. Valent, J. E. Bradner, S. W. Lowe, C. R. Vakoc, RNAi screen identifies Brd4 as a therapeutic target in acute myeloid leukaemia. Nature. 478, 524–528 (2011).

14. J. E. Delmore, G. C. Issa, M. E. Lemieux, P. B. Rahl, J. Shi, H. M. Jacobs, E. Kastritis, T. Gilpatrick, R. M. Paranal, J. Qi, M. Chesi, A. C. Schinzel, M. R. McKeown, T. P. Heffernan, C. R. Vakoc, P. L. Bergsagel, I. M. Ghobrial, P. G. Richardson, R. A. Young, W. C. Hahn, K. C. Anderson, A. L. Kung, J. E. Bradner, C. S. Mitsiades, BET bromodomain inhibition as a therapeutic strategy to target c-Myc. Cell. 146, 904–917 (2011).

15. M. A. Dawson, R. K. Prinjha, A. Dittmann, G. Giotopoulos, M. Bantscheff, W. Chan, S. C. Robson, C. Chung, C. Hopf, M. M. Savitski, C. Huthmacher, E. Gudgin, D. Lugo, S. Beinke, T. D. Chapman, E. J. Roberts, P. E. Soden, K. R. Auger, O. Mirguet, K. Doehner, R. Delwel, A. K. Burnett, P. Jeffrey, G. Drewes, K. Lee, B. J. P. Huntly, T. Kouzarides, Inhibition of BET recruitment to chromatin as an effective treatment for MLL-fusion leukaemia. Nature. 478, 529–533 (2011).

16. J. Shi, W. A. Whyte, C. J. Zepeda-Mendoza, J. P. Milazzo, C. Shen, J. S. Roe, J. L. Minder, F. Mercan, E. Wang, M. A. Eckersley-Maslin, A. E. Campbell, S. Kawaoka, S. Shareef, Z. Zhu, J. Kendall, M. Muhar, C. Haslinger, M. Yu, R. G. Roeder, M. H. Wigler, G. A. Blobel, J. Zuber, D. L. Spector, R. A. Young, C. R. Vakoc, Role of SWI/SNF in acute leukemia maintenance and enhancer-mediated Myc regulation. Genes Dev. 27, 2648–2662 (2013).

17. J. S. Roe, F. Mercan, K. Rivera, D. J. Pappin, C. R. Vakoc, BET Bromodomain Inhibition Suppresses the Function of Hematopoietic Transcription Factors in Acute Myeloid Leukemia. Mol. Cell. 58, 1028–1039 (2015).

18. V. A. Herzog, B. Reichholf, T. Neumann, P. Rescheneder, P. Bhat, T. R. Burkard, W. Wlotzka, A. Von Haeseler, J. Zuber, S. L. Ameres, Thiol-linked alkylation of RNA to assess expression dynamics. Nat. Methods. 14, 1198–1204 (2017).

19. A. Lusser, C. Gasser, L. Trixl, P. Piatti, I. Delazer, D. Rieder, J. Bashin, C. Riml, T. Amort, R. Micura, Thiouridine-to-Cytidine Conversion Sequencing (TUC-Seq) to Measure mRNA Transcription and Degradation Rates. Methods Mol. Biol. (2020), doi:10.1007/978-1-4939-9822-7_10.

20. J. A. Schofield, E. E. Duffy, L. Kiefer, M. C. Sullivan, M. D. Simon, TimeLapse-seq: Adding a temporal dimension to RNA sequencing through nucleoside recoding. Nat. Methods. 15, 221–225 (2018).

21. M. Michel, B. Zacher, C. Demel, A. Tresch, J. Gagneur, TT-seq maps the human transient transcriptome. Science (80-.). 352 (2016).

22. M. Rabani, J. Z. Levin, L. Fan, X. Adiconis, R. Raychowdhury, M. Garber, A. Gnirke, C. Nusbaum, N. Hacohen, N. Friedman, I. Amit, A. Regev, Metabolic labeling of RNA uncovers principles of RNA production and degradation dynamics in mammalian cells. Nat. Biotechnol. 29 (2011), doi:10.1038/nbt.1861.

23. E. E. Duffy, D. Catherine, R. R. Kitchen, B. Mark, M. D. Simon, E. E. Duffy, M. Rutenberg-schoenberg, C. D. Stark, R. R. Kitchen, M. B. Gerstein, M. D. Simon, Tracking Distinct RNA Populations Using Efficient and Reversible Covalent Chemistry Technology Tracking Distinct RNA Populations Using Efficient and Reversible Covalent Chemistry. Mol. Cell. 59, 858–866 (2015).

24. J. A. Mertz, A. R. Conery, B. M. Bryant, P. Sandy, S. Balasubramanian, D. A. Mele, L. Bergeron, R. J. Sims, Targeting MYC dependence in cancer by inhibiting BET bromodomains. Proc. Natl. Acad. Sci. U. S. A. 108, 16669–16674 (2011).

25. P. K. Parua, G. T. Booth, M. Sansó, B. Benjamin, J. C. Tanny, J. T. Lis, R. P. Fisher, A Cdk9-PP1 switch regulates the elongation-termination transition of RNA polymerase II. Nature. 558, 460–464 (2018).

26. Q. Jia, S. Chen, Y. Tan, Y. Li, F. Tang, Oncogenic super-enhancer formation in tumorigenesis and its molecular mechanisms. Exp. Mol. Med. 52, 713–723 (2020).

27. C. J. O. Alexander J. Federation, Donald R. Polaski, A. Fan, C. Y. Lin, J. E., Bradner, Identification of candidate master transcription factors within enhancer-centric transcriptional regulatory networks. bioRxiv (2018).

28. T. Yamada, N. Akimitsu, Contributions of regulated transcription and mRNA decay to the dynamics of gene expression. Wiley, 1–18 (2019).

29. Q. Wu, S. G. Medina, G. Kushawah, M. L. Devore, L. A. Castellano, J. M. Hand, M. Wright, A. A. Bazzini, Translation affects mRNA stability in a codon-dependent manner in human cells. Elife, 1–22 (2019).

30. O. Gilan, I. Rioja, K. Knezevic, M. J. Bell, M. M. Yeung, N. R. Harker, E. Y. N. Lam, C. Chung, P. Bamborough, M. Petretich, M. Urh, S. J. Atkinson, A. K. Bassil, E. J. Roberts, D. Vassiliadis, M. L. Burr, A. G. S. Preston, C. Wellaway, T. Werner, J. R. Gray, A. M. Michon, T. Gobbetti, V. Kumar, P. E. Soden, A. Haynes, J. Vappiani, D. F. Tough, S. Taylor, S. J. Dawson, M. Bantscheff, M. Lindon, G. Drewes, E. H. Demont, D. L. Daniels, P. Grandi, R. K. Prinjha, M. A. Dawson, Selective targeting of BD1 and BD2 of the BET proteins in cancer and immunoinflammation. Science (80-.). 368, 387–394 (2020).

31. A. Lal, F. Navarro, C. A. Maher, L. E. Maliszewski, N. Yan, E. O’Day, D. Chowdhury, D. M. Dykxhoorn, P. Tsai, O. Hofmann, K. G. Becker, M. Gorospe, W. Hide, J. Lieberman, miR-24 Inhibits Cell Proliferation by Targeting E2F2, MYC, and Other Cell-Cycle Genes via Binding to “Seedless” 3’UTR MicroRNA Recognition Elements. Mol. Cell. 35, 610–625 (2009).

32. M. Sachdeva, S. Zhu, F. Wu, H. Wu, V. Walia, S. Kumar, R. Elble, K. Watabe, Y. Y. Mo, p53 represses c-Myc through induction of the tumor suppressor miR-145. Proc. Natl. Acad. Sci. U. S. A. 106, 3207–3212 (2009).

33. N. M. Yeilding, M. T. Rehman, W. M. Lee, Identification of sequences in c-myc mRNA that regulate its steady-state levels. Mol. Cell. Biol. 16, 3511–3522 (1996).

34. D. Caput, B. Beutler, K. Hartog, R. Thayer, S. Brown-Shimer, A. Cerami, Identification of a common nucleotide sequence in the 3’-untranslated region of mRNA molecules specifying inflammatory mediators. Proc. Natl. Acad. Sci. U. S. A. (1986), doi:10.1073/pnas.83.6.1670.

35. K. Kataoka, Y. Shiraishi, Y. Takeda, S. Sakata, M. Matsumoto, S. Nagano, T. Maeda, Y. Nagata, A. Kitanaka, S. Mizuno, H. Tanaka, K. Chiba, S. Ito, Y. Watatani, N. Kakiuchi, H. Suzuki, T. Yoshizato, K. Yoshida, M. Sanada, H. Itonaga, Y. Imaizumi, Y. Totoki, W. Munakata, H. Nakamura, N. Hama, K. Shide, Y. Kubuki, T. Hidaka, T. Kameda, K. Masuda, N. Minato, K. Kashiwase, K. Izutsu, A. Takaori-Kondo, Y. Miyazaki, S. Takahashi, T. Shibata, H. Kawamoto, Y. Akatsuka, K. Shimoda, K. Takeuchi, T. Seya, S. Miyano, S. Ogawa, Aberrant PD-L1 expression through 3’-UTR disruption in multiple cancers. Nature. 534, 402–406 (2016).

36. M. W. Hentze, A. Castello, T. Schwarzl, T. Preiss, A brave new world of RNA-binding proteins. Nat. Rev. Mol. Cell Biol. 19, 327–341 (2018).

37. C. Ko, W. Wang, C. Li, Y. Jeng, Y. Chu, H. Wang, J. T. Tseng, J. Wang, IL-18-induced interaction between IMP3 and HuR contributes to COX-2 mRNA stabilization in acute myeloid leukemia. J. Leukoc. Biol. 99, 131–141 (2016).

38. A. Vujovic, L. de Rooij, A. Keyvani Chahi, H. T. Chen, B. A. Yee, S. K. Loganathan, L. Liu, D. C. H. Chan, A. Tajik, E. Tsao, S. Moreira, P. Joshi, J. Xu, N. Wong, Z. Balde, S. Jahangiri, S. Zandi, S. Aigner, J. E. Dick, M. D. Minden, D. Schramek, G. W. Yeo, K. J. Hope, In vivo screening unveils pervasive RNA-binding protein dependencies in leukemic stem cells and identifies ELAVL1 as a therapeutic target. Blood Cancer Discov. (2023), doi:10.1158/2643-3230.BCD-22-0086.

39. K. Rothamel, ELAVL1 Exclusively Couples mRNA Stability with the 3’UTRs of Interferon Stimulated Genes. bioRxiv (2020).

40. R. Kuuselo, K. Savinainen, S. Sandström, R. Autio, A. Kallioniemi, MED29, a component of the mediator complex, possesses both oncogenic and tumor suppressive characteristics in pancreatic cancer. Int. J. Cancer. 129, 2553–2565 (2011).

41. T. C. Lin, C. Y. Su, P. Y. Wu, T. C. Lai, W. A. Pan, Y. H. Jan, Y. C. Chang, C. T. Yeh, C. L. Chen, L. P. Ger, H. T. Chang, C. J. Yang, M. S. Huang, Y. P. Liu, Y. F. Lin, J. Y. J. Shyy, M. D. Tsai, M. Hsiao, The nucleolar protein NIFK promotes cancer progression via ck1α/β-catenin in metastasis and ki-67-dependent cell proliferation. Elife. 5, 1–21 (2016).

42. F. F. Blanco, R. Preet, A. Aguado, V. Vishwakarma, L. E. Stevens, A. Vyas, S. Padhye, L. Xu, S. J. Weir, S. Anant, N. Meisner-Kober, J. R. Brody, D. A. Dixon, Impact of HuR inhibition by the small molecule MS-444 on colorectal cancer cell tumorigenesis. Oncotarget (2016), doi:10.18632/oncotarget.12189.

43. D. Hnisz, B. J. Abraham, T. I. Lee, A. Lau, V. Saint-André, A. A. Sigova, H. A. Hoke, R. A. Young, Super-enhancers in the control of cell identity and disease. Cell (2013), doi:10.1016/j.cell.2013.09.053.

44. B. E. Gryder, L. Wu, G. M. Woldemichael, S. Pomella, T. R. Quinn, P. M. C. Park, A. Cleveland, B. Z. Stanton, Y. Song, R. Rota, O. Wiest, M. E. Yohe, J. F. Shern, J. Qi, J. Khan, Chemical genomics reveals histone deacetylases are required for core regulatory transcription. Nat. Commun. 10, 1–12 (2019).

45. B. Schwanhüusser, D. Busse, N. Li, G. Dittmar, J. Schuchhardt, J. Wolf, W. Chen, M. Selbach, Global quantification of mammalian gene expression control. Nature. 473, 337–342 (2011).

46. C. Mayr, D. P. Bartel, Widespread Shortening of 3’UTRs by Alternative Cleavage and Polyadenylation Activates Oncogenes in Cancer Cells. Cell. 138, 673–684 (2009).

47. C. Vicente, R. Stirparo, S. Demeyer, C. E. De Bock, O. Gielen, M. Atkins, J. Yan, G. Halder, B. A. Hassan, J. Cools, The CCR4-NOT complex is a tumor suppressor in Drosophila melanogaster eye cancer models. J. Hematol. Oncol. (2018), doi:10.1186/s13045-018-0650-0.

48. S. Weißbach, C. Langer, B. Puppe, T. Nedeva, E. Bach, M. Kull, R. Bargou, H. Einsele, A. Rosenwald, S. Knop, E. Leich, The molecular spectrum and clinical impact of DIS3 mutations in multiple myeloma. Br. J. Haematol. (2015), doi:10.1111/bjh.13256.

49. C. Desterke, A. Bennaceur-Griscelli, A. G. Turhan, DIS3 Mutation in RUNX1-Mutated AML1 Confers a Highly Dismal Prognosis in AML By Repressing Sister Chromatid Cohesion. Blood (2019), doi:10.1182/blood-2019-123352.

50. D. F. Aghib, J. M. Bishop, A 3’ truncation of myc caused by chromosomal translocation in a human T-cell leukemia is tumorigenic when tested in established rat fibroblasts. Oncogene. 6, 2371–5 (1991).

51. D. F. Aghib, J. M. Bishop, S. Ottolenghi, A. Guerrasio, A. Serra, G. Saglio, A 3’ truncation of MYC caused by chromosomal translocation in a human T-cell leukemia increases mRNA stability. Oncogene. 5, 707–11 (1990).

52. Z. Xia, L. A. Donehower, T. A. Cooper, J. R. Neilson, D. A. Wheeler, E. J. Wagner, W. Li, Dynamic analyses of alternative polyadenylation from RNA-seq reveal a 3’2-UTR landscape across seven tumour types. Nat. Commun. 5 (2014), doi:10.1038/ncomms6274.

53. F. Yuan, W. Hankey, E. J. Wagner, W. Li, Q. Wang, Alternative polyadenylation of mRNA and its role in cancer. Genes Dis. 8, 61–72 (2021).

54. S. Kammler, S. Lykke-Andersen, T. H. Jensen, The RNA exosome component hRrp6 is a target for 5-fluorouracil in human cells. Mol. Cancer Res. 6, 990–995 (2008).

55. R. A. Silverstein, E. G. De Valdivia, N. Visa, The incorporation of 5-fluorouracil into RNA affects the ribonucleolytic activity of the exosome subunit Rrp6. Mol. Cancer Res. 9, 332–340 (2011).

56. T. Yamauchi, T. Masuda, M. C. Canver, M. Seiler, Y. Semba, M. Shboul, M. Al-Raqad, M. Maeda, V. A. C. Schoonenberg, M. A. Cole, C. Macias-Trevino, Y. Ishikawa, Q. Yao, M. Nakano, F. Arai, S. H. Orkin, B. Reversade, S. Buonamici, L. Pinello, K. Akashi, D. E. Bauer, T. Maeda, Genome-wide CRISPR-Cas9 Screen Identifies Leukemia-Specific Dependence on a Pre-mRNA Metabolic Pathway Regulated by DCPS. Cancer Cell. 33, 386–400.e5 (2018).

57. D. Hanahan, R. A. Weinberg, Hallmarks of cancer: The next generation. Cell. 144, 646–674 (2011).

58. M. Rashkovan, A. Ferrando, Metabolic dependencies and vulnerabilities in leukemia. Genes Dev. 33, 1460–1474 (2019).

59. K. Rothamel, S. Arcos, B. Kim, C. Reasoner, S. Lisy, N. Mukherjee, K. Rothamel, S. Arcos, B. Kim, C. Reasoner, S. Lisy, N. Mukherjee, ELAVL1 primarily couples mRNA stability with the 3 0 UTRs of interferon-stimulated genes ll ll ELAVL1 primarily couples mRNA stability with the 3 0 UTRs of interferon-stimulated genes. Cell Rep. 35, 109178 (2021).

60. S. Heinzel, T. Binh Giang, A. Kan, J. M. Marchingo, B. K. Lye, L. M. Corcoran, P. D. Hodgkin, A Myc-dependent division timer complements a cell-death timer to regulate T cell and B cell responses. Nat. Immunol. (2017), doi:10.1038/ni.3598.

61. C. Ye, D. J. Ho, M. Neri, C. Yang, T. Kulkarni, R. Randhawa, M. Henault, N. Mostacci, P. Farmer, S. Renner, R. Ihry, L. Mansur, C. G. Keller, G. Mcallister, M. Hild, J. Jenkins, A. Kaykas, DRUG-seq for miniaturized high-throughput transcriptome profiling in drug discovery. Nat. Commun., 1–9 (2018).

62. A. Subramanian, P. Tamayo, V. K. Mootha, S. Mukherjee, B. L. Ebert, M. A. Gillette, A. Paulovich, S. L. Pomeroy, T. R. Golub, E. S. Lander, J. P. Mesirov, Gene set enrichment analysis: A knowledge-based approach for interpreting genome-wide expression profiles. Proc. Natl. Acad. Sci. U. S. A. (2005), doi:10.1073/pnas.0506580102.

63. J. Chen, H. Xu, B. J. Aronow, A. G. Jegga, Improved human disease candidate gene prioritization using mouse phenotype. BMC Bioinformatics. 8, 1–13 (2007).

64. J. Chen, E. E. Bardes, B. J. Aronow, A. G. Jegga, ToppGene Suite for gene list enrichment analysis and candidate gene prioritization. Nucleic Acids Res. 37, 305–311 (2009).

65. J. Chen, B. J. Aronow, A. G. Jegga, Disease candidate gene identification and prioritization using protein interaction networks. BMC Bioinformatics. 10, 1–14 (2009).

66. Z. Wang, J. Hu, E. W. Johnson, J. D. Campbell, Scruff: An R/Bioconductor package for preprocessing single-cell RNA-sequencing data. bioRxiv, 1–9 (2019).

67. T. Stuart, A. Butler, P. Hoffman, C. Hafemeister, E. Papalexi, W. M. Mauck, Y. Hao, M. Stoeckius, P. Smibert, R. Satija, Comprehensive Integration of Single-Cell Data. Cell (2019), doi:10.1016/j.cell.2019.05.031.

68. H. A. Hoke, C. Y. Lin, A. Lau, D. A. Orlando, C. R. Vakoc, J. E. Bradner, T. I. Lee, R. A. Young, Selective Inhibition of Tumor Oncogenes by Disruption of Super-Enhancers (2013), doi:10.1016/j.cell.2013.03.036.

69. S. J. Vervoort, S. A. Welsh, J. R. Devlin, E. Barbieri, D. A. Knight, S. Offley, S. Bjelosevic, M. Costacurta, I. Todorovski, C. J. Kearney, J. J. Sandow, Z. Fan, B. Blyth, V. McLeod, J. H. A. Vissers, K. Pavic, B. P. Martin, G. Gregory, E. Demosthenous, M. Zethoven, I. Y. Kong, E. D. Hawkins, S. J. Hogg, M. J. Kelly, A. Newbold, K. J. Simpson, O. Kauko, K. F. Harvey, M. Ohlmeyer, J. Westermarck, N. Gray, A. Gardini, R. W. Johnstone, The PP2A-Integrator-CDK9 axis fine-tunes transcription and can be targeted therapeutically in cancer. Cell. 184, 3143–3162.e32 (2021).

70. B. Sundararaman, L. Zhan, S. M. Blue, R. Stanton, K. Elkins, S. Olson, X. Wei, E. L. Van Nostrand, G. A. Pratt, S. C. Huelga, B. M. Smalec, X. Wang, E. L. Hong, J. M. Davidson, E. Lécuyer, B. R. Graveley, G. W. Yeo, Resources for the Comprehensive Discovery of Functional RNA Elements. Mol. Cell. 61, 903–913 (2016).

71. B. Li, C. N. Dewey, RSEM: accurate transcript quantification from RNA-Seq data with or without a reference genome. BMC Bioinformatics. 12, 323 (2011).

72. S. J. Vervoort, S. A. Welsh, J. R. Devlin, N. Gray, A. Gardini, R. W. Johnstone, The PP2A-Integrator-CDK9 axis fine-tunes transcription and can be targeted therapeutically in cancer. Cell, 1–20 (2021).

