## Supplementary_data for "RNA decay defines the response to transcriptional perturbation in leukaemia"

**Supplementary figures**

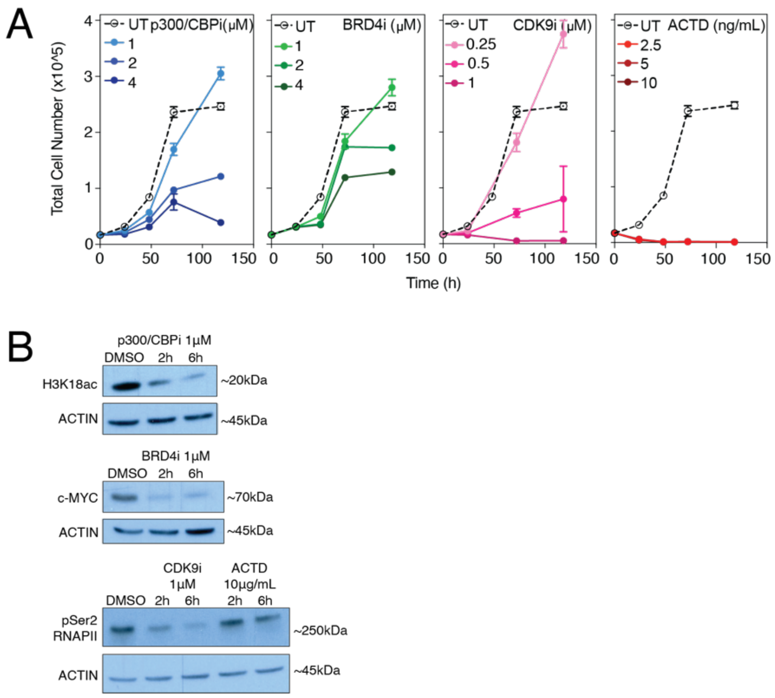


**Supplementary 1**

**(A)** Total cell numbers of K562 cells treated with increasing concentrations of p300/CBPi, BRD4i, CDK9i and ACTD at indicated time points. Experiments were performed in biological triplicate. **(B)** **(Top)** Western blot of H3K18ac and ACTIN protein following treatment with 1µM of p300/CBPi. **(Middle)** Western blot of c-MYC and ACTIN protein following treatment with 1µM of BRD4i. **(Bottom)** Western blot of phospho-Serine 2 (pSer2) residue of the RNAPII carboxy-terminal domain (CTD) and ACTIN protein following treatment with 1µM of CDK9i or 10µg/mL ACTD. Each western blot is a representative image from three biological replicates performed at time points indicated. kDa: kiloDalton. H: hour

16

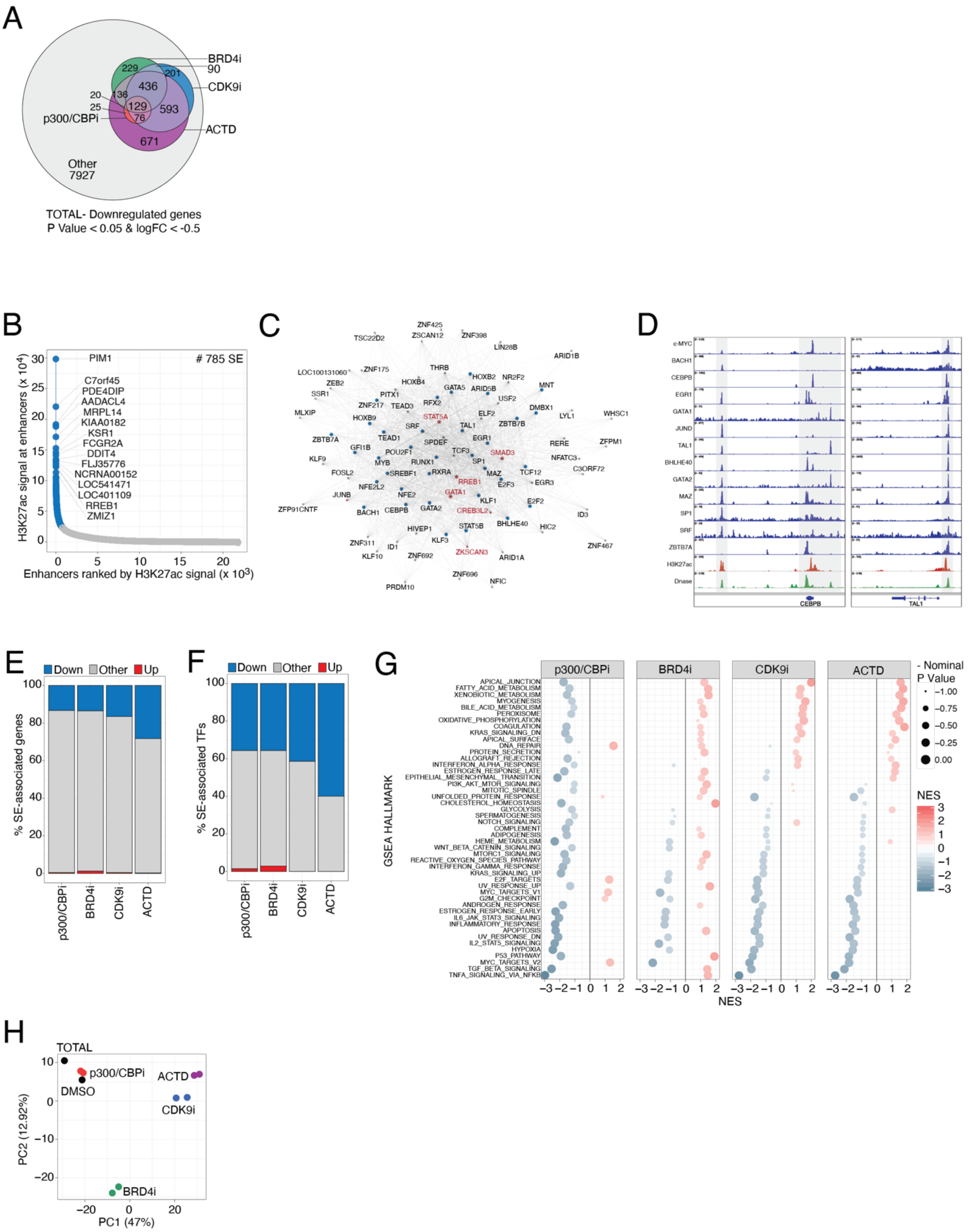
A
BRD4i
90
CDK9i
ACTD
p300/CBPi
229
201
436
20
136
25
129
76
593
671
Other
7927
TOTAL- Downregulated genes
P Value < 0.05 & logFC < -0.5
B
# 785 SE
PIM1
C7orf45
PDE4DIP
AADACL4
MRPL14
KIAA0182
KSR1
FCGR2A
DDIT4
FLJ35776
NCRNA00152
LOC541471
LOC401109
RREB1
ZMIZ1
H3K27ac signal at enhancers (x 10^4)
Enhancers ranked by H3K27ac signal (x 10^3)
C
D
c-MYC
BACH1
CEBPB
EGR1
GATA1
JUND
TAL1
BHLHE40
GATA2
MAZ
SP1
SRF
ZBTB7A
H3K27ac
Dnase
CEBPB
TAL1
E
Down
Other
Up
% SE-associated genes
p300/CBPi
BRD4i
CDK9i
ACTD
F
Down
Other
Up
% SE-associated TFs
p300/CBPi
BRD4i
CDK9i
ACTD
G
p300/CBPi
BRD4i
CDK9i
ACTD
GSEA HALLMARK
- Nominal P Value
NES
H
TOTAL
p300/CBPi
DMSO
ACTD
CDK9i
BRD4i
PC2 (12.92%)
PC1 (47%)

17

**Supplementary 2**

**(A)** Venn diagram of significantly down-regulated genes for each treatment condition using spike in normalized total mRNA reads. Remaining genes indicated as 'other.' **(B)** Super-enhancer (SEs; blue) and enhancers (grey) ranked by H3K27ac signal. A total of 785 SEs were identified and genes associated with the top 15 are highlighted. **(C)** Network plot of SE-associated core-regulatory (CR) transcription factors (TFs) identified using the Coltron algorithm (*1*). TFs previously identified as part of K562 cell core regulatory networks highlighted in red (*2*). **(D)** Integrated Genomics Viewer (IGV) screen shots of *CEBPB* and *TAL1* genomic loci with CR TF occupancy obtained from ENCODE. **(E)** Bar chart of significantly up- (red) or down-regulated (blue) SE-associated genes and **(F)** SE-associated TFs upon two hours of transcription inhibition. **(G)** Gene Set Enrichment Analysis (GSEA) Normalized Enrichment Scores (NES) and significance of cancer hallmark pathways using ranked total mRNA changes upon two hours of transcription inhibition. **(H)** Principal Component Analysis (PCA) plot of spike-in normalized total mRNA reads.

logFC: log2 fold change. PC: principal component. Significantly up-regulated: P Value < 0.05 & logFC > 0.5. Significantly down-regulated: P Value < 0.05 & logFC < -0.5. ****, **, *: P Value < 0.0001, < 0.01 and <0.05, respectively, using an unpaired Wilcoxon test.

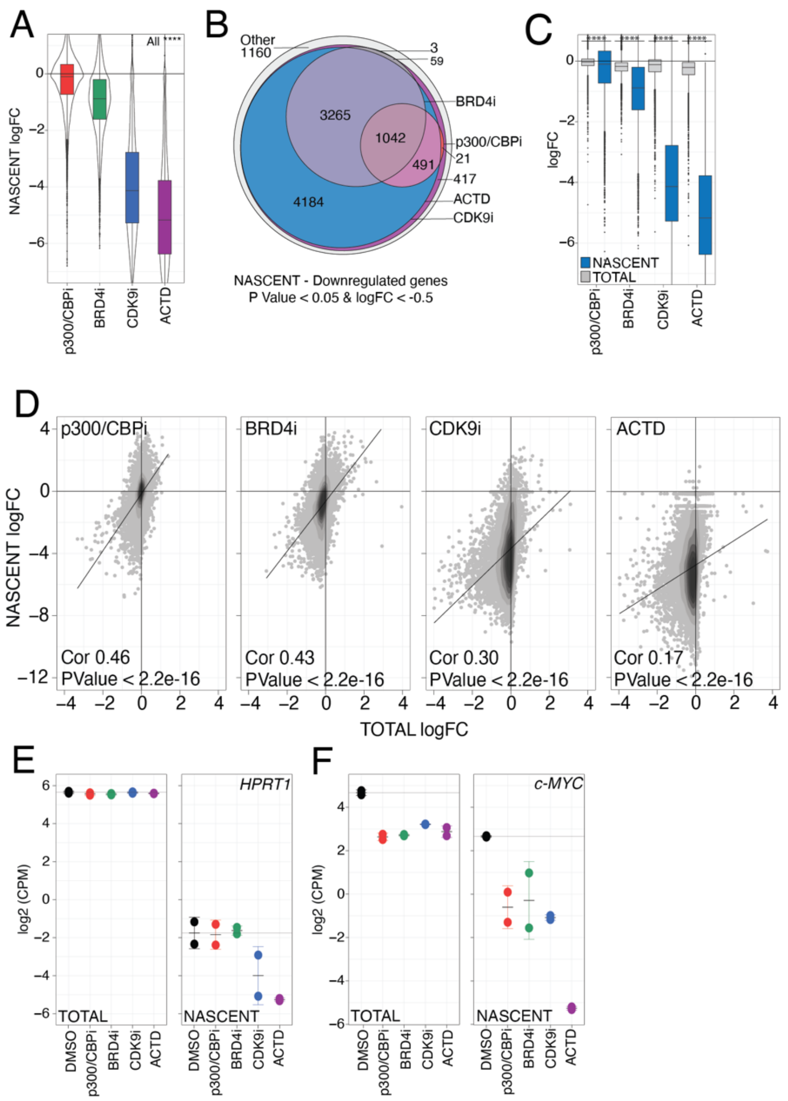


**Supplementary 3**

**(A)** Boxplot of spike-in normalized nascent gene expression upon two hours of transcription inhibition. **(B)** Venn diagram of significantly down-regulated genes for each treatment condition using spike in normalized nascent mRNA reads. Remaining genes indicated as ‘other.’ **(C)** Boxplot and **(D)** scatterplot of spike-in normalized total and nascent gene expression upon two hours of transcription inhibition. **(E)** Spike-in normalized **(left)** total and **(right)** nascent mRNA expression of *HPRT1* or **(F)** *c-MYC* upon two hours of transcription inhibition.

logFC: log2 fold change. Cor: Pearson’s correlation co-efficient. CPM: Counts Per Million. ****: P Value < 0.0001 using an unpaired Wilcoxon test.

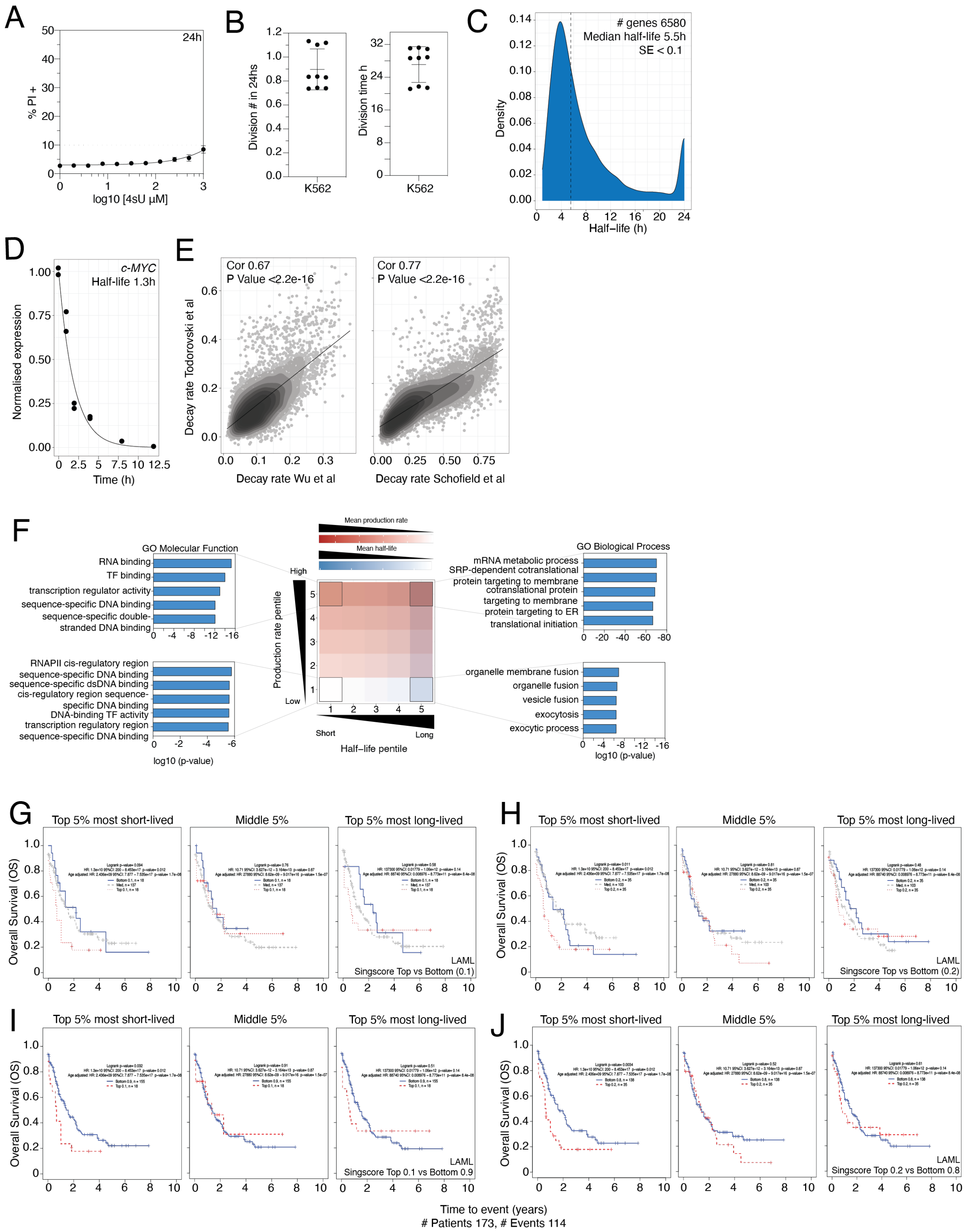
A
24h
% PI +
log10 [4sU µM]
B
Division # in 24hs
Division time h
K562
C
### genes 6580
Median half-life 5.5h
SE < 0.1
Density
Half−life (h)
D
c-MYC
Half-life 1.3h
Normalised expression
Time (h)
E
Cor 0.67
P Value <2.2e-16
Cor 0.77
P Value <2.2e-16
Decay rate Todorovski et al
Decay rate Wu et al
Decay rate Schofield et al
F
GO Molecular Function
RNA binding
TF binding
transcription regulator activity
sequence-specific DNA binding
sequence-specific double-stranded DNA binding
RNAPII cis-regulatory region sequence-specific DNA binding
sequence-specific dsDNA binding
cis-regulatory region sequence-specific DNA binding
DNA-binding TF activity
transcription regulatory region sequence-specific DNA binding
log10 (p-value)
Mean production rate
Mean half-life
High
Low
Production rate pentile
Short
Long
Half−life pentile
GO Biological Process
mRNA metabolic process
SRP-dependent cotranslational protein targeting to membrane
cotranslational protein targeting to membrane
protein targeting to ER
translational initiation
organelle membrane fusion
organelle fusion
vesicle fusion
exocytosis
exocytic process
log10 (p-value)
G
Top 5% most short-lived
Middle 5%
Top 5% most long-lived
Overall Survival (OS)
LAML
Singscore Top vs Bottom (0.1)
H
Top 5% most short-lived
Middle 5%
Top 5% most long-lived
Overall Survival (OS)
LAML
Singscore Top vs Bottom (0.2)
I
Top 5% most short-lived
Middle 5%
Top 5% most long-lived
Overall Survival (OS)
LAML
Singscore Top 0.1 vs Bottom 0.9
J
Top 5% most short-lived
Middle 5%
Top 5% most long-lived
Overall Survival (OS)
LAML
Singscore Top 0.2 vs Bottom 0.8
Time to event (years)
### Patients 173, # Events 114

**Supplementary 4**

**(A)** Cell death (Propidium Iodide (PI) positive) assessment of K562 cells following treatment with increasing concentration of 4sU following 24 hours. Values represented are mean with error bars indicating standard deviation (sd) from three technical and biological replicates. **(B) (Left)** Division number and **(right)** time of K562 cells as assessed by Cell Trace Violet (CTV) labelling. Mean division time of 27.1 hours was calculated using values from three technical and biological replicates. Error bars indicating sd shown. **(C)** Distribution of mRNA stability of 6580 genes in the K562 cell line. Half-life values represented are from technical duplicates and normalized for K562 cell division time. **(D)** *c-MYC* transcript half-life estimation. Half-life values represented are from technical duplicates and normalized for K562 cell division time. **(E) (Left)** Scatter plot of K562 decay measurements obtained in this study and (*3*) or **(right)** (*4*). **(F)** Heatmap of mean mRNA half-life (blue) and production rate (red) for each mRNA decay and production pentile. Gene Ontology (GO) terms of genes within subsets highlighted in black are indicated in bar charts. **(G)** Kaplan-Meier curve of TCGA AML patients stratified according to top 10% versus bottom 10% gene expression, **(H)** top 20% versus bottom 20% gene expression, **(I)** top 10% versus bottom 90% gene expression and **(J)** top 20% versus bottom 80% gene expression of **(left)** short-, **(middle)** medium- and **(right)** long-lived transcripts.

H: hour. Cor: Pearson's correlation co-efficient.

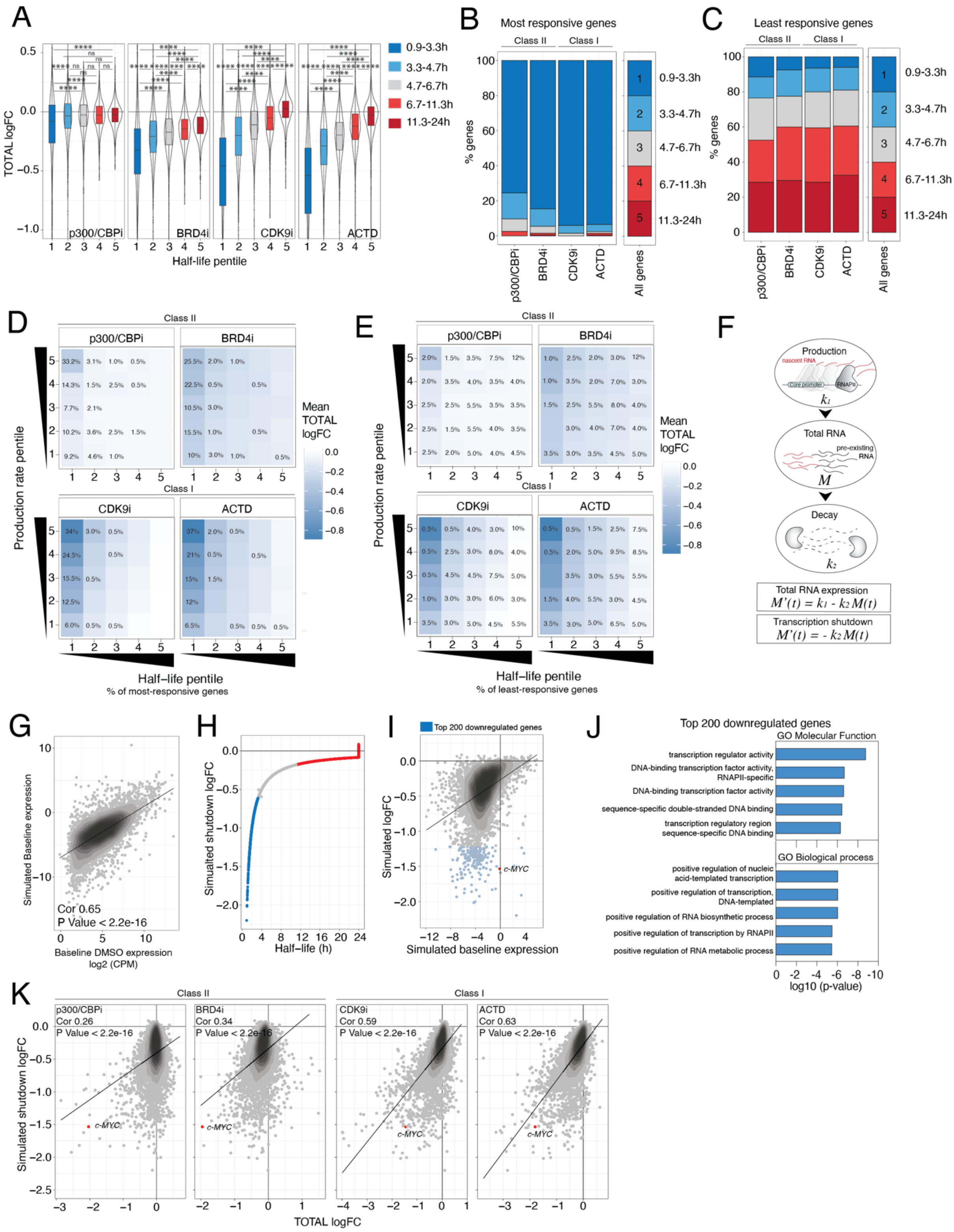
A
TOTAL logFC
p300/CBPi
BRD4i
CDK9i
ACTD
Half-life pentile
0.9-3.3h
3.3-4.7h
4.7-6.7h
6.7-11.3h
11.3-24h
B
Most responsive genes
Class II
Class I
% genes
p300/CBPi
BRD4i
CDK9i
ACTD
All genes
C
Least responsive genes
D
Class II
Class I
Production rate pentile
Mean TOTAL logFC
Half−life pentile
% of most-responsive genes
E
% of least-responsive genes
F
Production
nascent RNA
Core promoter
RNAPII
k1
Total RNA
pre-existing RNA
M
Decay
k2
Total RNA expression
M'(t) = k1 - k2 M(t)
Transcription shutdown
M'(t) = - k2 M(t)
G
Simulated Baseline expression
Cor 0.65
P Value < 2.2e-16
Baseline DMSO expression
log2 (CPM)
H
Simulated shutdown logFC
Half-life (h)
I
Top 200 downregulated genes
Simulated logFC
c-MYC
Simulated baseline expression
J
Top 200 downregulated genes
GO Molecular Function
transcription regulator activity
DNA-binding transcription factor activity, RNAPII-specific
DNA-binding transcription factor activity
sequence-specific double-stranded DNA binding
transcription regulatory region sequence-specific DNA binding
GO Biological process
positive regulation of nucleic acid-templated transcription
positive regulation of transcription, DNA-templated
positive regulation of RNA biosynthetic process
positive regulation of transcription by RNAPII
positive regulation of RNA metabolic process
log10 (p-value)
K
Class II
Class I
p300/CBPi
Cor 0.26
P Value < 2.2e-16
BRD4i
Cor 0.34
P Value < 2.2e-16
CDK9i
Cor 0.59
P Value < 2.2e-16
ACTD
Cor 0.63
P Value < 2.2e-16
Simulated shutdown logFC
c-MYC
TOTAL logFC

**Supplementary 5**

**(A)** Boxplot of total gene expression response with inhibitors indicated of gene groups (1-5) obtained by dividing expressed genes into pentiles according to mRNA half-life. **(B)** Stacked bar chart of mRNA half-life pentiles in genes most-responsive and **(C)** least-responsive to treatments indicated. **(D)** Heatmap of mean total mRNA logFC for each mRNA decay and production pentile. Numbers indicated are percentage of most-responsive and **(E)** least-responsive genes within each subset. **(F)** Mathematical model of mRNA production ($k_1$) and decay ($k_2$). **(*Top*)** Differential equation describing total mRNA. **(*Bottom*)** Simulation of total mRNA levels with complete transcription shutdown. **(G)** Scatter plot of measured total DMSO and simulated baseline expression. **(H)** Simulated total gene expression response following two hours of complete transcription shutdown ranked by RNA half-life. The 20% shortest- and longest- lived genes indicated in blue and red, respectively. **(I)** Scatter plot of simulated baseline expression versus simulated change in total gene expression with total transcription shutdown for two hours. Top 200 genes most down-regulated indicated in blue. **(J)** GO analysis of the top 200 most down-regulated genes with simulated total transcription shutdown for two hours indicated in blue in (I). **(K)** Scatter plot of simulated total gene expression response following two hours of complete transcription shutdown and measured total gene expression response to inhibitors indicated.
logFC: log2 fold change. H: hour. Cor: Pearson’s correlation co-efficient.

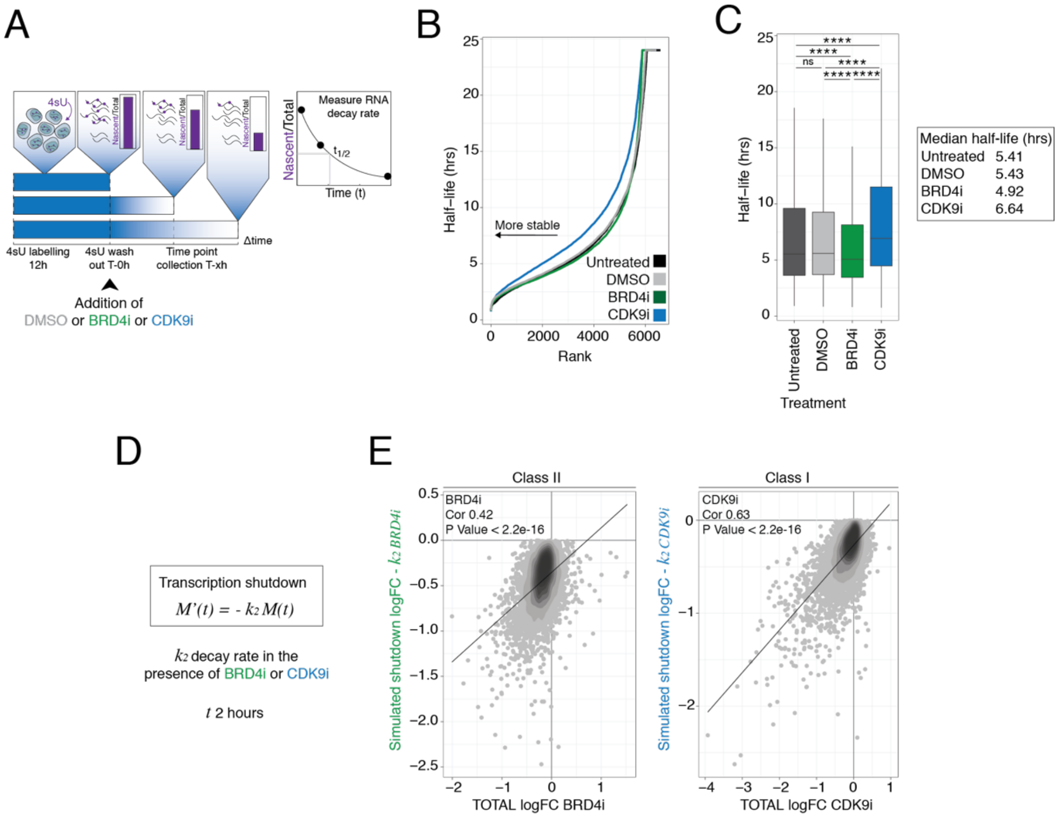


**Supplementary 6**

**(A)** Schematic of 4sU pulse-labelling and chase in the presence of therapeutic transcriptional inhibitors in the K562 cell line. T>C conversion rates were calculated for each time point and treatment condition, normalised to 0 hours and fit with exponential decay functions to derive half-life (t ½). **(B)** Ranked curve and **(C)** boxplot of measured RNA half-lives in K562 cells treated with DMSO, JQ1 (BRD4i) and AZ-5576 (CDK9i). Prior stability measurements are indicated as ‘untreated.’ Median half-life (hours) for each group are indicated. Experiment was performed as a single replicate. **(D)** Mathematical model used to simulate complete transcriptional shutdown (RNA production rate 0) using decay measurements (k2) obtained in K562 cells treated with BRD4i or CDK9i as shown in (A-C). M: total RNA. T: time. **(E)** Scatter plot of simulated total gene expression response with decay rates from (D) and measured total RNA changes to the inhibitors indicated. ****, ns: P Value < 0.0001 or not significant respectively, using an unpaired Wilcoxon test.

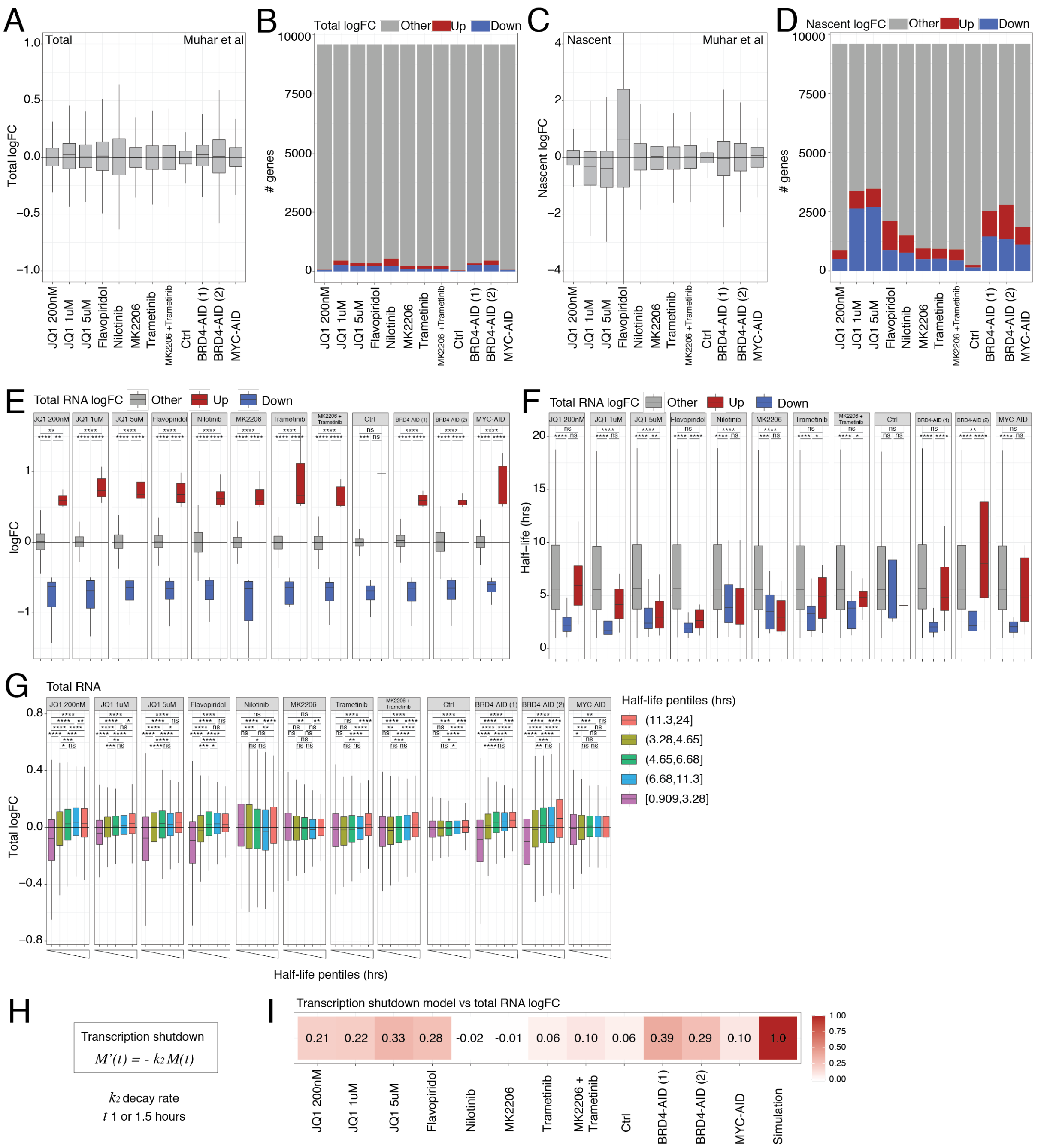


**Supplementary 7**

**(A)** Boxplot of total mRNA log2 fold change (logFC) in response to inhibitors and degradation of AID-tagged proteins indicated from (*5*). **(B)** Number of significantly up- (logFC > 0.5 and P Value < 0.05) and down-regulated (logFC < -0.5 and P Value < 0.05) genes on the total mRNA level using data shown in (A). Remaining genes indicated as 'other.' **(C)** Boxplot of nascent RNA logFC in response to inhibitors and degradation of AID-tagged proteins indicated from (*5*). **(D)** Number of significantly up- and down-regulated genes on the nascent mRNA level using data shown from (C). Remaining genes indicated as 'other.' **(E)** Boxplot of total RNA logFC and **(F)** **m**RNA half-life (in hours) of significantly up- and down-regulated genes. Remaining genes indicated as 'other.' **(G)** Total RNA logFC of mRNA half-life pentiles in response to inhibitors and degradation of proteins indicated from (*5*). **(H)** Mathematical model used to simulate complete transcriptional shutdown (RNA production rate 0) using decay measurements (k2) obtained in K562 cells. **(I)** Correlation matrix of simulated total gene expression response from (H) and measured total RNA changes to the inhibitors indicated from (A). Pearson's correlation co-efficients indicated.

M: total RNA. T: time. ****, ***,** ,* , ns: P Value < 0.0001, < 0.001, <0.01, < 0.1 or not significant respectively, using an unpaired Wilcoxon test.

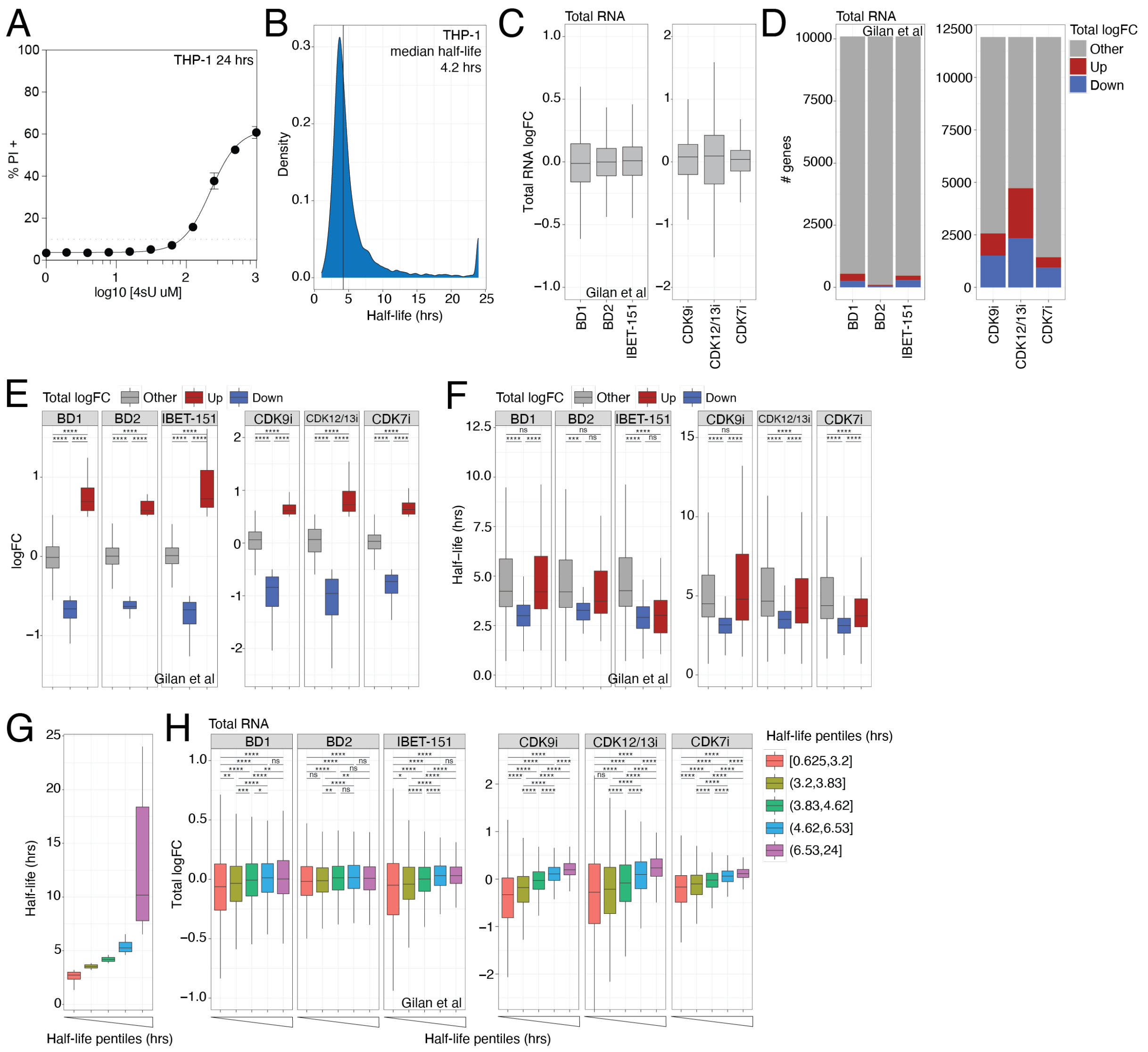


**Supplementary 8**

**(A)** Curve of THP-1 cell death (propidium iodide (PI) positive) with increasing concentrations of 4sU after 24 hours of treatment. Experiment was performed in biological duplicate. **(B)** Distribution of mRNA stability of 7001 genes in the THP-1 cell line. Measurements were obtained according to the methodology described as a single replicate. **(C)** **(Left)** Box plot of total RNA log2 fold change (logFC) of THP-1 cells treated with BD1, BD2 and IBET-151 relative to DMSO from (*5*) and **(right)** AZ-5576 (CDK9i), THZ531 (CDK12/13i) and YKL-5-124 (CDK7i) relative to DMSO. Experiment on (*right*) was performed in technical triplicate. **(D)** Bar chart of the number of significantly up- (logFC > 0.5 and P Value < 0.05) and down-regulated (logFC < -0.5 and P Value < 0.05) genes on the total RNA level using data from (A). Remaining genes indicated as 'other.' **(E)** Boxplot of total RNA logFC and **(F)** RNA half -life (hours) of significantly up- and down-regulated genes on the total RNA level from (B). **(G)** Boxplot of RNA half-life (hours) across THP-1 RNA half-life pentiles determined using measurements from (D). **(H)** Total RNA logFC of RNA half-life pentile groups from (G).
****, ***, ** ,* , ns: P Value < 0.0001, < 0.001, <0.01, < 0.1 or not significant respectively, using an unpaired Wilcoxon test.

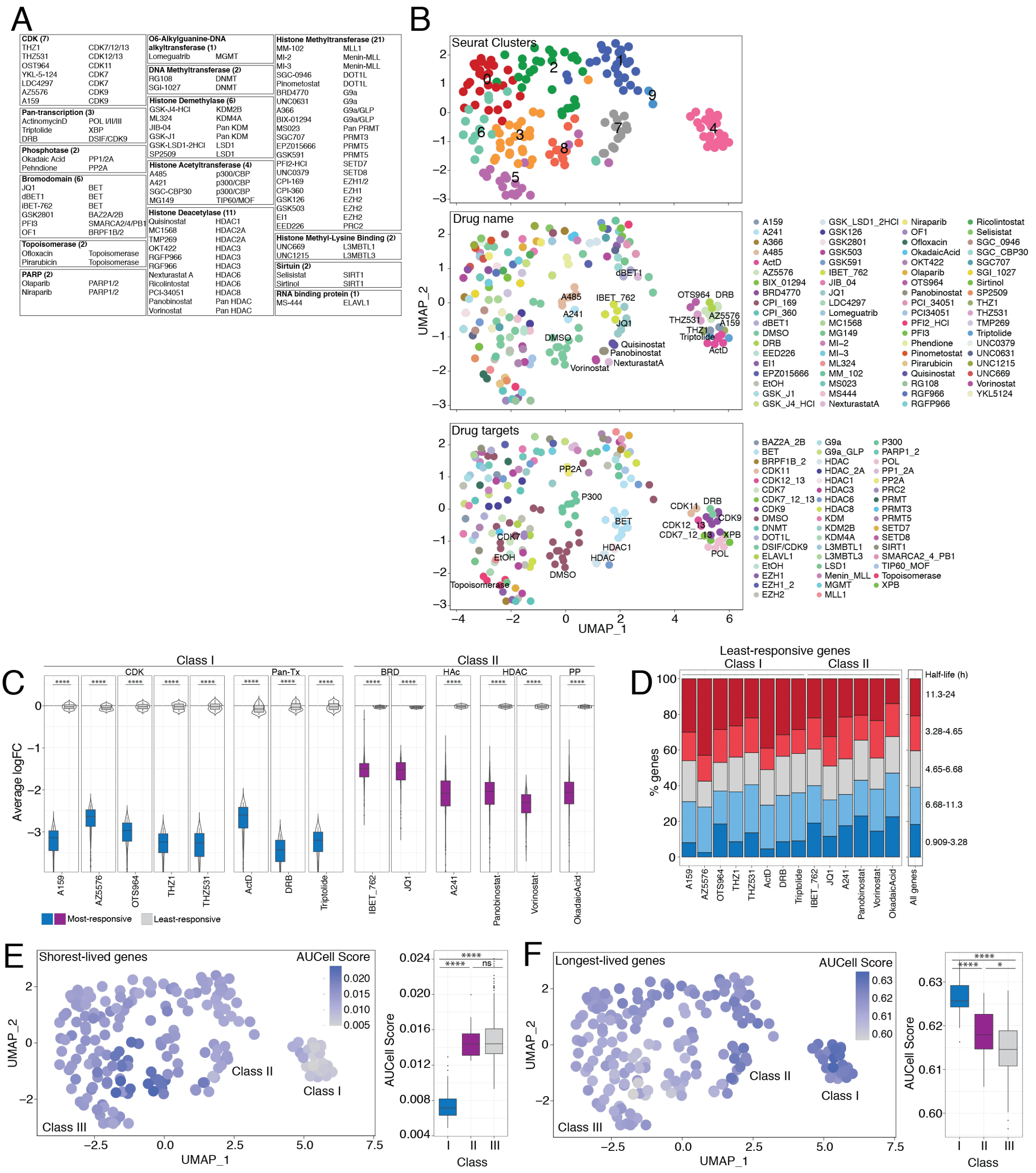
A
CDK (7)
THZ1 CDK7/12/13
THZ531 CDK12/13
OST964 CDK11
YKL-5-124 CDK7
LDC4297 CDK7
AZ5576 CDK9
A159 CDK9
Pan-transcription (3)
ActinomycinD POL I/II/III
Triptolide XBP
DRB DSIF/CDK9
Phosphotase (2)
Okadaic Acid PP1/2A
Pehndione PP2A
Bromodomain (6)
JQ1 BET
dBET1 BET
iBET-762 BET
GSK2801 BAZ2A/2B
PFI3 SMARCA2/4/PB1
OF1 BRPF1B/2
Topoisomerase (2)
Ofloxacin Topoisomerase
Pirarubicin Topoisomerase
PARP (2)
Olaparib PARP1/2
Niraparib PARP1/2
O6-Alkylguanine-DNA alkyltransferase (1)
Lomeguatrib MGMT
DNA Methyltransferase (2)
RG108 DNMT
SGI-1027 DNMT
Histone Demethylase (6)
GSK-J4-HCl KDM2B
ML324 KDM4A
JIB-04 Pan KDM
GSK-J1 Pan KDM
GSK-LSD1-2HCl LSD1
SP2509 LSD1
Histone Acetyltransferase (4)
A485 p300/CBP
A421 p300/CBP
SGC-CBP30 p300/CBP
MG149 TIP60/MOF
Histone Deacetylase (11)
Quisinostat HDAC1
MC1568 HDAC2A
TMP269 HDAC2A
OKT422 HDAC3
RGFP966 HDAC3
RGF966 HDAC3
Nexturastat A HDAC6
Ricolintostat HDAC6
PCI-34051 HDAC8
Panobinostat Pan HDAC
Vorinostat Pan HDAC
Histone Methyltransferase (21)
MM-102 MLL1
MI-2 Menin-MLL
MI-3 Menin-MLL
SGC-0946 DOT1L
Pinometostat DOT1L
BRD4770 G9a
UNC0631 G9a
A366 G9a/GLP
BIX-01294 G9a/GLP
MS023 Pan PRMT
SGC707 PRMT3
EPZ015666 PRMT5
GSK591 PRMT5
PFI2-HCl SETD7
UNC0379 SETD8
CPI-169 EZH1/2
CPI-360 EZH1
GSK126 EZH2
GSK503 EZH2
EI1 EZH2
EED226 PRC2
Histone Methyl-Lysine Binding (2)
UNC669 L3MBTL1
UNC1215 L3MBTL3
Sirtuin (2)
Selisistat SIRT1
Sirtinol SIRT1
RNA binding protein (1)
MS-444 ELAVL1
B
Seurat Clusters
Drug name
Drug targets
UMAP_2
UMAP_1
C
Class I
Class II
CDK
Pan-Tx
BRD
HAc
HDAC
PP
Average logFC
A159
AZ5576
OTS964
THZ1
THZ531
ActD
DRB
Triptolide
IBET_762
JQ1
A241
Panobinostat
Vorinostat
OkadaicAcid
Most-responsive
Least-responsive
D
Least-responsive genes
Class I
Class II
% genes
All genes
Half-life (h)
11.3-24
3.28-4.65
4.65-6.68
6.68-11.3
0.909-3.28
E
Shortest-lived genes
AUCell Score
Class I
Class II
Class III
UMAP_1
UMAP_2
AUCell Score
Class
F
Longest-lived genes
AUCell Score
Class I
Class II
Class III
UMAP_1
UMAP_2
AUCell Score
Class

**Supplementary 9**

**(A)** Table of inhibitors used and associated protein target grouped by protein class. **(B)** UMAP plot of treatment conditions with (***top***) Seurat clusters, ***(middle)*** drug treatment and (***bottom***) protein target highlighted. **(C)** Boxplot of average logFC of genes most- and least-responsive to class I and II inhibitors. **(D)** Bar chart of mRNA half-life pentiles in genes least-responsive to class I and II inhibitors. (**E)** ***(Left)*** UMAP plot with AUCell signature score for the top 20% shortest-lived genes highlighted for each treatment condition. ***(Right)*** AUCell Score summarised for each drug class defined in Fig. 3C. **(F)** Same as S4. E except scores were generated for the top 20% longest-lived genes.

logFC: log2 fold change relative to DMSO/EtOH. Most-responsive: top 200 most significantly down-regulated genes (logFC < -0.5 and P Value < 0.05) using spike-in normalized reads. Least-responsive: 200 unaltered (-0.25 < logFC < 0.25 and P Value > 0.05) genes using spike-in normalized reads. ****, *, ns: P Value < 0.0001, < 0.05 or not significant respectively, using an unpaired Wilcoxon test.

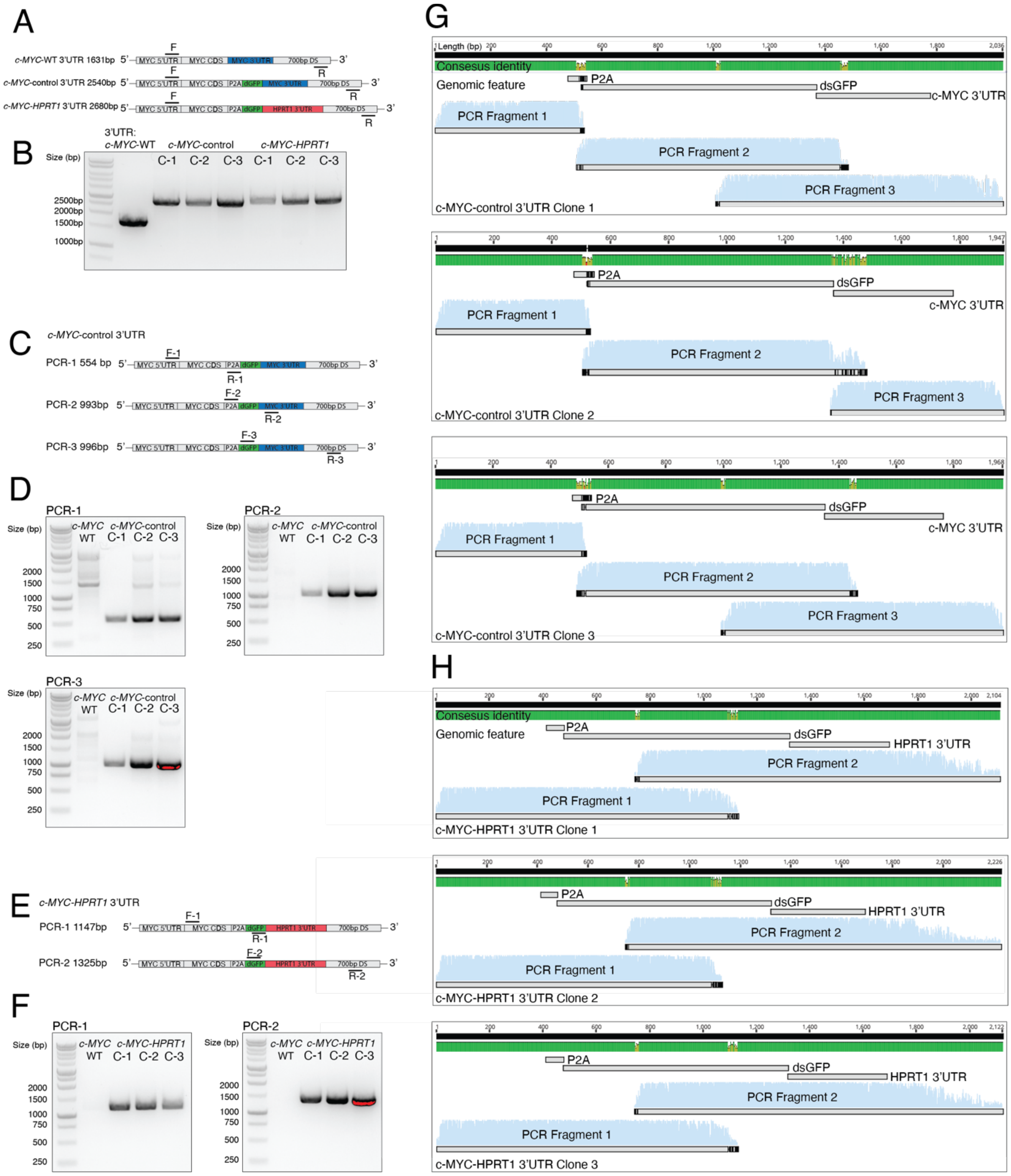


**Supplementary 10**

**(A)** Schematic and **(B)** agarose gel image of DNA fragments PCR amplified with primers designed outside CRISPR/Cas9-HDR donor plasmid in parental and three single cell clones of either *c-MYC*-control and *-HPRT1* 3'UTR cell lines. **(C)** Schematic and **(D)** agarose gel image of DNA fragments PCR amplified with primers designed to span knock-in locus of *c-MYC*-control or **(E-F)** *-HPRT1* 3'UTR single cell clones. **(G)** Screen shots from the Geneious Prime (v2020.0.4) program of Sanger sequenced PCR fragments from (C)-(D) aligning to P2A, dsGFP and *c-MYC* 3'UTR sequences. **(H)** Same as (G), except Sanger sequenced PCR fragments from (E)-(F) aligning to P2A, dsGFP and *HPRT1* 3'UTR sequences. Consensus identify indicated above each alignment, where 100% identity is represented as green.

C- 1-3: Single cell clones 1-3. F: forward primer. R: reverse primer.

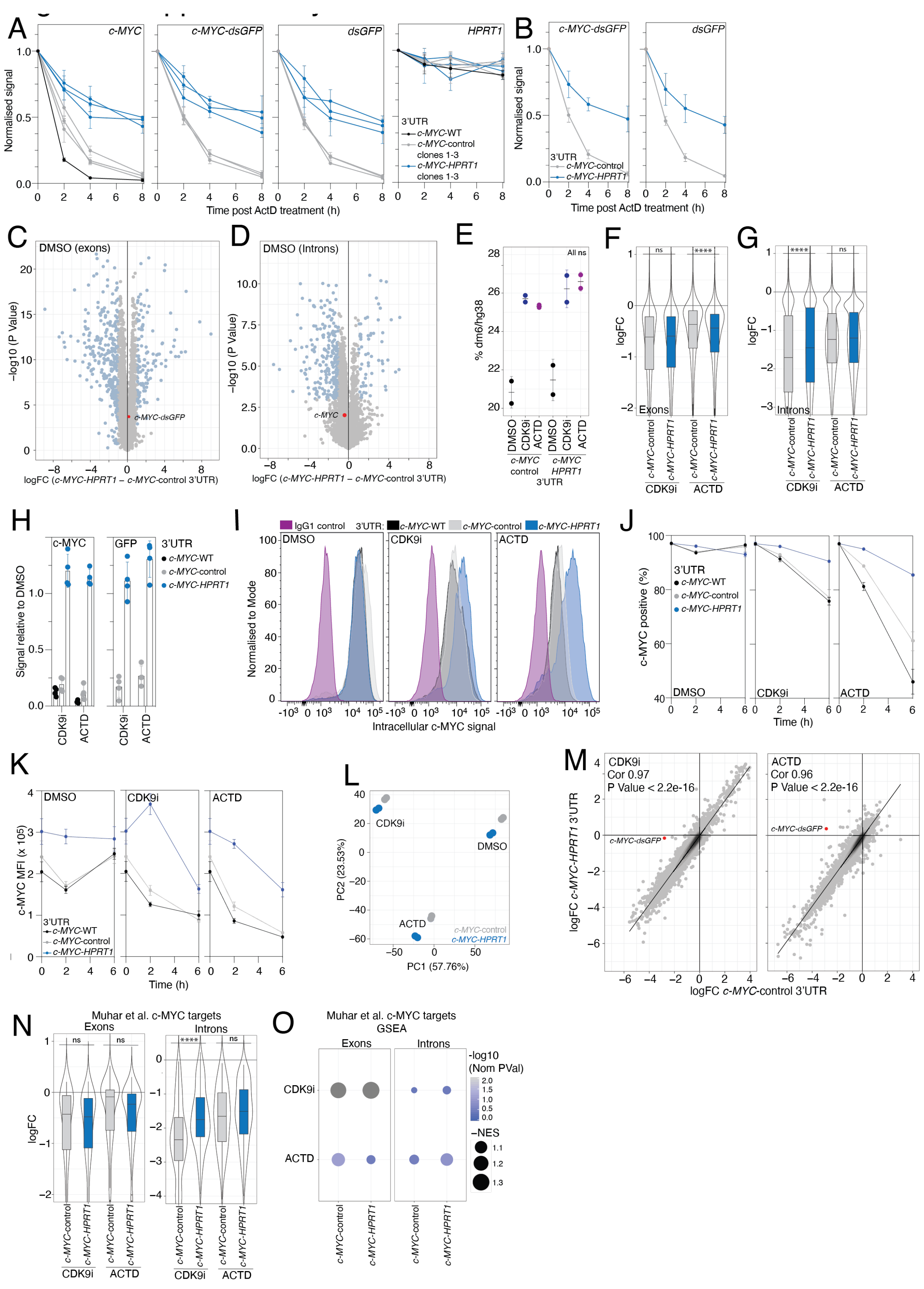
A
c-MYC
c-MYC-dsGFP
dsGFP
HPRT1
Normalised signal
3'UTR
c-MYC-WT
c-MYC-control clones 1-3
c-MYC-HPRT1 clones 1-3
Time post ActD treatment (h)
B
c-MYC-dsGFP
dsGFP
3'UTR
c-MYC-control
c-MYC-HPRT1
Time post ActD treatment (h)
C
DMSO (exons)
−log10 (P Value)
c-MYC-dsGFP
logFC (c-MYC-HPRT1 – c-MYC-control 3'UTR)
D
DMSO (Introns)
c-MYC
logFC (c-MYC-HPRT1 – c-MYC-control 3'UTR)
E
All ns
% dm6/hg38
DMSO
CDK9i
ACTD
c-MYC control
c-MYC HPRT1
3'UTR
F
ns
****
logFC
Exons
c-MYC-control
c-MYC-HPRT1
CDK9i
ACTD
G
****
ns
Introns
H
c-MYC
GFP
3'UTR
c-MYC-WT
c-MYC-control
c-MYC-HPRT1
Signal relative to DMSO
I
IgG1 control
3'UTR: c-MYC-WT c-MYC-control c-MYC-HPRT1
DMSO
CDK9i
ACTD
Normalised to Mode
Intracellular c-MYC signal
J
3'UTR
c-MYC-WT
c-MYC-control
c-MYC-HPRT1
c-MYC positive (%)
Time (h)
K
DMSO
CDK9i
ACTD
c-MYC MFI (x 105)
3'UTR
c-MYC-WT
c-MYC-control
c-MYC-HPRT1
Time (h)
L
CDK9i
DMSO
ACTD
c-MYC-control
c-MYC-HPRT1
PC2 (23.53%)
PC1 (57.76%)
M
CDK9i
Cor 0.97
P Value < 2.2e-16
ACTD
Cor 0.96
P Value < 2.2e-16
c-MYC-dsGFP
logFC c-MYC-HPRT1 3'UTR
logFC c-MYC-control 3'UTR
N
Muhar et al. c-MYC targets
Exons
Introns
ns
****
logFC
c-MYC-control
c-MYC-HPRT1
CDK9i
ACTD
O
Muhar et al. c-MYC targets
GSEA
Exons
Introns
CDK9i
ACTD
-log10 (Nom PVal)
−NES
c-MYC-control
c-MYC-HPRT1

**Supplementary 11**

**(A)** Normalized expression of **(left)** *c-MYC*, **(middle-left)** *c-MYC-dsGFP*, **(middle-right)** *dsGFP* and **(right)** *HPRT1* transcripts following the addition of ACTD and collection of RNA at indicated time points as measured by quantitative real time PCR (qRT-PCR). Values are mean with error bars representing standard deviation (sd) of three biological replicates for each single cell clone. **(B)** Same as (A) *left* and *middle-left*, except values are mean with error bars representing standard deviation (sd) of three biological replicates of single cell clones *combined* for each cell line. **(C)** Scatter plot of significance and difference in exon and **(D)** intron gene expression in DMSO controls between representative *c-MYC*-control and *-HPRT1* 3'UTR single cell clones. Significantly (adjusted P Value < 0.05) altered genes are highlighted in blue. **(E)** Percentage ratio of reads mapping to BDGP6/dm6 and GRCh38/hg38 genomes. **(F)** Boxplot of spike-in normalized exon and **(G)** intron gene expression relative to DMSO. **(H)** Western blot protein signal of **(left)** c-MYC and **(right)** GFP relative to ACTIN and DMSO controls after 6 hours of treatment. Values are mean with error bars representing sd of three biological replicates. **(I)** Representative histogram of intracellular c-MYC protein staining following 6 hours of indicated treatments. **(J)** Proportion of cells positive and **(K)** Mean Fluorescent Intensity (MFI) of intracellular c-MYC protein staining following 0, 2 and 6 hours with indicated treatments. Values are mean with error bars representing sd of two biological replicates. **(L)** PCA plot of spike-in normalized exon gene expression. **(M)** Scatter plot of change spike-in normalised exon expression in *c-MYC*-control and *-HPRT1* 3'UTR cell lines with **(left)** CDK9i and **(right)** ACTD treatments relative to DMSO. **(N)** **(Left)** Boxplot of change in exon or **(Right)** intron expression of *c-MYC* target genes (*5*) with indicated treatments and cell lines. **(O)** GSEA Normalized Enrichment Scores (NES) and significance of MYC targets using ranked total mRNA changes.

H: hour. logFC: log2 fold change relative to DMSO. ****, ns; P Value < 0.0001 and not significant, respectively, using an unpaired Wilcoxon test. PC: Principal component. Cor: Pearson's correlation coefficient. Nom P Val: Nominal P Value.

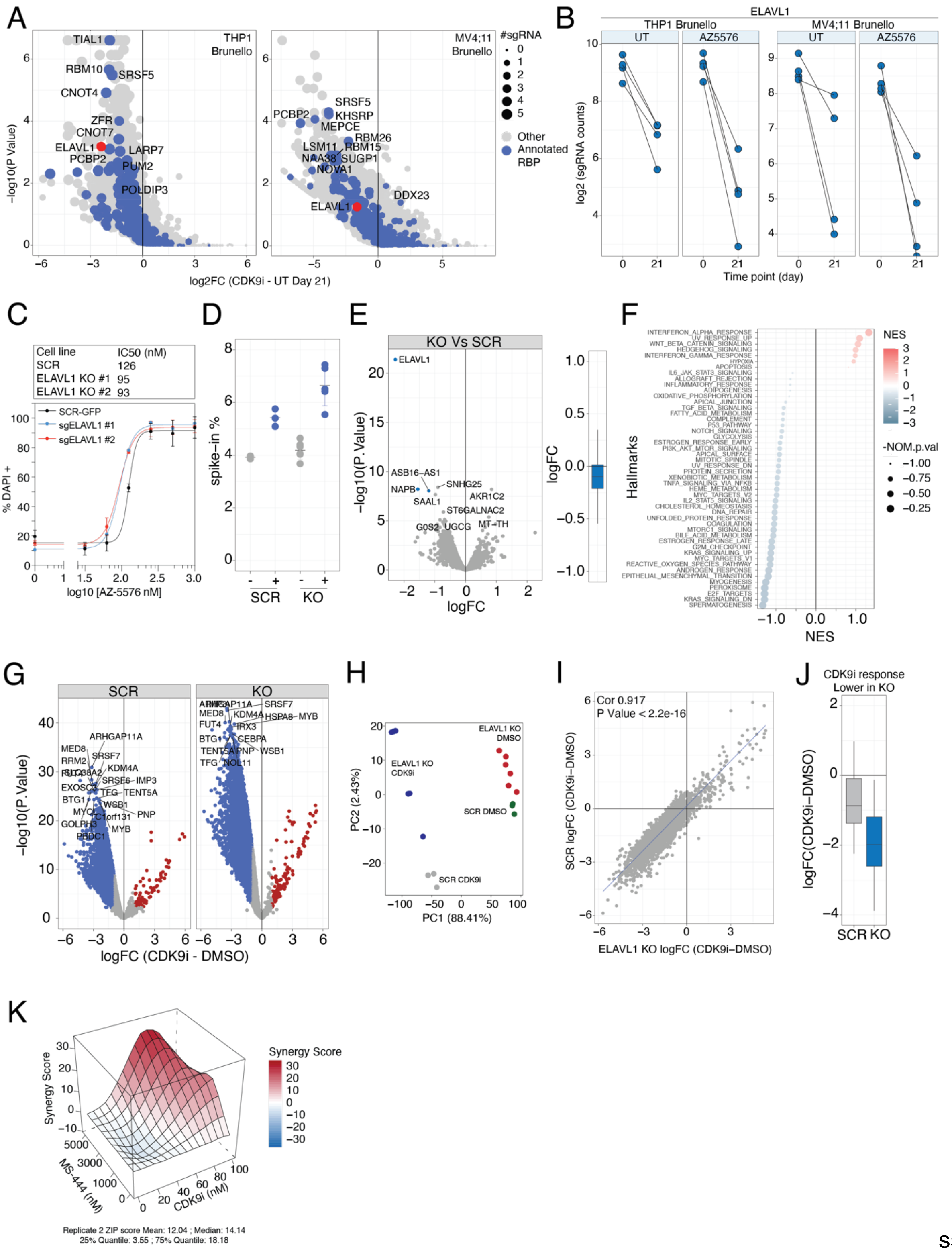
A
THP1 Brunello
MV4;11 Brunello
#sgRNA
0
1
2
3
4
5
Other
Annotated RBP
TIAL1
RBM10
SRSF5
CNOT4
ZFR
CNOT7
ELAVL1
LARP7
PCBP2
PUM2
POLDIP3
PCBP2
SRSF5
KHSRP
MEPCE
RBM26
LSM11
RBM15
NAA38
SUGP1
NOVA1
DDX23
ELAVL1
−log10(P Value)
log2FC (CDK9i - UT Day 21)
B
ELAVL1
THP1 Brunello
MV4;11 Brunello
UT
AZ5576
UT
AZ5576
log2 (sgRNA counts)
Time point (day)
C
Cell line IC50 (nM)
SCR 126
ELAVL1 KO #1 95
ELAVL1 KO #2 93
SCR-GFP
sgELAVL1 #1
sgELAVL1 #2
% DAPI +
log10 [AZ-5576 nM]
D
spike−in %
SCR
KO
E
KO Vs SCR
ELAVL1
ASB16−AS1
NAPB
SNHG25
SAAL1
AKR1C2
ST6GALNAC2
G0S2
UGCG
MT−TH
−log10(P. Value)
logFC
F
Hallmarks
NES
-NOM.p.val
INTERFERON_ALPHA_RESPONSE
UV_RESPONSE_UP
WNT_BETA_CATENIN_SIGNALING
HEDGEHOG_SIGNALING
INTERFERON_GAMMA_RESPONSE
HYPOXIA
APOPTOSIS
IL6_JAK_STAT3_SIGNALING
ALLOGRAFT_REJECTION
INFLAMMATORY_RESPONSE
ADIPOGENESIS
OXIDATIVE_PHOSPHORYLATION
APICAL_JUNCTION
TGF_BETA_SIGNALING
FATTY_ACID_METABOLISM
COMPLEMENT
P53_PATHWAY
NOTCH_SIGNALING
GLYCOLYSIS
ESTROGEN_RESPONSE_EARLY
PI3K_AKT_MTOR_SIGNALING
APICAL_SURFACE
MITOTIC_SPINDLE
UV_RESPONSE_DN
PROTEIN_SECRETION
XENOBIOTIC_METABOLISM
TNFA_SIGNALING_VIA_NFKB
HEME_METABOLISM
MYC_TARGETS_V2
IL2_STAT5_SIGNALING
CHOLESTEROL_HOMEOSTASIS
DNA_REPAIR
UNFOLDED_PROTEIN_RESPONSE
COAGULATION
MTORC1_SIGNALING
BILE_ACID_METABOLISM
ESTROGEN_RESPONSE_LATE
G2M_CHECKPOINT
KRAS_SIGNALING_UP
MYC_TARGETS_V1
REACTIVE_OXYGEN_SPECIES_PATHWAY
ANDROGEN_RESPONSE
EPITHELIAL_MESENCHYMAL_TRANSITION
MYOGENESIS
PEROXISOME
E2F_TARGETS
KRAS_SIGNALING_DN
SPERMATOGENESIS
G
SCR
KO
−log10(P. Value)
logFC (CDK9i - DMSO)
H
ELAVL1 KO DMSO
ELAVL1 KO CDK9i
SCR DMSO
SCR CDK9i
PC2 (2.43%)
PC1 (88.41%)
I
Cor 0.917
P Value < 2.2e-16
SCR logFC (CDK9i−DMSO)
ELAVL1 KO logFC (CDK9i−DMSO)
J
CDK9i response Lower in KO
logFC(CDK9i−DMSO)
SCR KO
K
Synergy Score
MS-444 (nM)
CDK9i (nM)
Replicate 2 ZIP score Mean: 12.04 ; Median: 14.14
25% Quantile: 3.55 ; 75% Quantile: 18.18

**Supplementary 12**

**(A)** Scatter plot of significance and change in sgRNA read counts in CDK9i compared to untreated (UT) CRISPR-Cas9 genome-wide screening groups after 21 days of passaging in **(left)** THP-1 and **(right)** MV4;11 AML cell lines. Annotated RBPs indicated in blue and number of sgRNAs indicated by point size. Data is obtained from (*6*). **(B)** sgRNA read counts targeting ELAVL1 in CRISPR-Cas9 genome-wide screening groups and treatments indicated in **(left)** THP-1 and **(right)** MV4;11 cell lines. **(C)** Dose-response curve of THP-1 cell death (DAPI positive) expressing Cas9 and non-targeting sgRNA (SCR) or sgRNA targeting ELAVL1 (ELAVL1 KO) following four days of doxycycline induction treated with CDK9i for 72 hours. Half maximal inhibitory concentrations (IC50) for each cell line are indicated. **(D)** Percentage ratio of reads mapping to BDGP6/dm6 and GRCh38/hg38 genomes. **(E)** **(Left)** Scatter plot of significance and difference in spike-in normalized total gene expression in DMSO treated ELAVL1 knockout relative to SCR cells. Significantly down-regulated genes (logFC < -1 and adjusted P Value < 0.05) are highlighted in blue. **(Right)** Boxplot of logFC derived from data in (left). **(F)** GSEA Normalized Enrichment Scores (NES) and significance of cancer hallmark pathways using ranked spike-in normalized total mRNA changes from comparisons between DMSO treated ELAVL1 knockout and SCR cells. **(G)** Scatter plot of significance and difference in spike-in normalized total gene expression with CDK9i relative to DMSO in **(left)** SCR and **(right)** ELAVL1 knockout cells. Significantly up- and down-regulated genes indicated in red and blue, respectively. **(H)** PCA of spike-in normalized total mRNA reads. **(I)** Scatter plot of spike-in normalized total gene expression with CDK9i relative to DMSO in SCR and ELAVL1 knockout cells. **(J)** Boxplot of spike-in normalized total mRNA reads with CDK9i relative to DMSO across cell lines indicated of genes sensitized to CDK9i upon ELAVL1 knockout (lower in KO; logFC < -1 and adjusted P Value < 0.01). **(K)** ZIP synergy scores of THP-1 cells treated with CDK9i and MS-444 at concentrations indicated for 72 hours. Representative of second replicate.

#### Supplementary tables

**Table. S1.**

| Compound | Provider | Cat # | Reconstitution |
|---|---|---|---|
| 4-thiouridine | Sigma Aldrich | T4509 | Ultrapure DNAse/RNAse-free distilled water (Thermo Fisher Scientific, 10977023) |
| Uridine | Sigma Aldrich | U6381 | Ultrapure DNAse/RNAse-free distilled water |
| A-159 (A-1592668) | Sigma-Aldrich | 472301 | DMSO |
| A-1641241241 | Provided by Abbvie | | DMSO |
| A-1619485 | Provided by Abbvie | | DMSO |
| ACTD | Sigma Aldrich | A1410 | DMSO |
| AZ-5576 | Provided by AstraZeneca | | DMSO |
| dBET1 | Cayman Chemical Company | 18044 | DMSO |
| DRB | Sigma-Aldrich | D1916 | DMSO |
| EED226 | Selleckchem | S8496 | DMSO |
| iBET-151 | Provided by Roche | | DMSO |
| GSK126 | Provided by GSK | | DMSO |
| GSK503 | Provided by GSK | | Ethanol |
| JQ1 | Provided by Roche | | DMSO |
| Okadaic Acid | Abcam | ab120375 | DMSO |
| OKT422 | Obtained from OnKure Therapeutics | | DMSO |
| Panobinostat | Provided by Novartis | | DMSO |
| PCI34061 | Xcessbio | M60178 | DMSO |
| Phendione | Sigma Aldrich | 496383 | DMSO |
| RGF966 | Sigma Aldrich | SML1652 | DMSO |
| Ricolinostat | Xcessbio | M60076 | DMSO |
| THZ1 | Provided by Nathaneal Gray | | DMSO |
| THZ531 | AOBIOUS, Gloucester, MA, USA | AOB8107 | DMSO |
| Triptolide | Abcam | ab120720 | DMSO |
| Vorinostat | Provided by Merek, Boston, MA USA | | DMSO |
| YKL-5-124 | Selleckchem | S8863 | DMSO |

**Table. S2.**

| Compounds Australia (CA) | | |
|---|---|---|
| Name | Catalog Number | CA Sample Number |
| GSK J4 HCl | S7070 | SN01730839 |
| ML324 | S7296 | SN01730861 |
| JIB-04 | S7281 | SN01730857 |
| GSK J1 | S7581 | SN01730878 |
| GSK-LSD1 2HCl | S7574 | SN01730877 |
| SP2509 | S7680 | SN01730891 |
| BRD4770 | S7591 | SN01730880 |
| UNC0631 | S7610 | SN01730882 |

| | | |
|---|---|---|
| BIX 01294 | S8006 | SN01730904 |
| A-366 | S7572 | SN01730876 |
| CPI-360 | S7656 | SN01730890 |
| GSK503 | S7804 | SN01730897 |
| EI1 | S7611 | SN01730883 |
| CPI-169 | S7616 | SN01730884 |
| MI-2 (Menin-MLL Inhibitor) | S7618 | SN01730885 |
| MI-3 (Menin-MLL Inhibitor) | S7619 | SN01730886 |
| MM-102 | S7265 | SN01730855 |
| EPZ015666(GSK3235025) | S7748 | SN01730893 |
| GSK591 | S8111 | SN01730911 |
| SGC707 | S7832 | SN01730899 |
| MS023 | S8112 | SN01730912 |
| MC1568 | S1484 | SN01730788 |
| TMP269 | S7324 | SN01730866 |
| Quisinostat (JNJ-26481585) 2HCl | S1096 | SN01730759 |
| RGFP966 | S7229 | SN01730848 |
| PCI-34051 | S2012 | SN01730794 |
| Nexturastat A | S7473 | SN01730871 |
| Selisistat (EX 527) | S1541 | SN01730792 |
| Sirtinol | S2804 | SN01730820 |
| RG108 | S2821 | SN01730823 |
| SGI-1027 | S7276 | SN01730856 |
| Pinometostat (EPZ5676) | S7062 | SN01730838 |
| SGC 0946 | S7079 | SN01730840 |
| PFI-2 HCl | S7294 | SN01730859 |
| UNC0379 | S7570 | SN01730875 |
| Olaparib (AZD2281 or Ku-0059436) | S1060 | SN01730754 |
| Niraparib (MK-4827) tosylate | S7625 | SN01730888 |
| I-BET-762 | S7189 | SN01730847 |
| SGC-CBP30 | S7256 | SN01730854 |
| Pirarubicin | S1393 | SN01730782 |
| Ofloxacin | S1463 | SN01730787 |
| Lomeguatrib | S8056 | SN01730908 |
| UNC1215 | S7088 | SN01730841 |
| UNC669 | S7373 | SN01730869 |
| GSK2801 | S7231 | SN01730849 |
| PFI-3 | S7315 | SN01730865 |
| MG149 | S7476 | SN01730872 |
| OF-1 | S7681 | SN01730892 |

**Table. S3.**

| Singe Guide RNA sequences | |
|---|---|
| sgRNA | Sequence (5'-3') |
| Non-targeting control (SCR) | GCCTAGTCTCGGTAAGAGTC |

| | |
|---|---|
| ELAVL1 #1 (Brunello ID 5490) | TACCTGTAGTCTGATCCACG |
| ELAVL1 #2 (Brunello ID 5491) | TATCCGGTTTGACAAACGGT |
| c-MYC STOP codon | CTTGTGCGTAAGGAAAAGTA |
| c-MYC 3' of 3'UTR | AACATTCACAACTTAAGATT |

**Table. S4.**

| Vector insert sequences | |
|---|---|
| Vector | Insert sequence (5'-3') |
| PUC57 ECORV-MYC-GSG-P2A-D1EGFP-STOP-MYC-UTR- ECORV (c-MYC CONTROL 3’UTR) | ACCTTCTGCTGGAGGCCACAGCAAACCTCCTCACAGCCCACTGGTCCTCAA<br>GAGGTGCCACGTCTCCACACATCAGCACAACTACGCAGCGCCTCCCTCCAC<br>TCGGAAGGACTATCCTGCTGCCAAGAGGGTCAAGTTGGACAGTGTCAGAGT<br>CCTGAGACAGATCAGCAACAACCGAAAATGCACCAGCCCCAGGTCCTCGGA<br>CACCGAGGAGAATGTCAAGAGGCGAACACACAACGTCTTGGAGCGCCAGA<br>GGAGGAACGAGCTAAAACGGAGCTTTTTTGCCCTGCGTGACCAGATCCCGG<br>AGTTGGAAAACAATGAAAAGGCCCCCAAGGTAGTTATCCTTAAAAAAGCCAC<br>AGCATACATCCTGTCCGTCCAAGCAGAGGAGCAAAAGCTCATTTCTGAAGA<br>GGACTTGTTGCGGAAACGACGAGAACAGTTGAAACACAAACTTGAACAGCT<br>ACGGAACTCTTGTGCGGGCAGCGGCGCCACAAACTTCTCTCTGCTAAAGCA<br>AGCAGGTGATGTTGAAGAAAACCCCGGGCCTGTGAGCAAGGGCGAGGAGC<br>TGTTCACCGGGGTGGTGCCCATCCTGGTCGAGCTGGACGGCGACGTAAAC<br>GGCCACAAGTTCAGCGTGTCCGGCGAGGGCGAGGGCGATGCCACCTACGG<br>CAAGCTGACCCTGAAGTTCATCTGCACCACCGGCAAGCTGCCCGTGCCCTG<br>GCCCACCCTCGTGACCACCCTGACCTACGGCGTGCAGTGCTTCAGCCGCTA<br>CCCCGACCACATGAAGCAGCACGACTTCTTCAAGTCCGCCATGCCCGAAGG<br>CTACGTCCAGGAGCGCACCATCTTCTTCAAGGACGACGGCAACTACAAGAC<br>CCGCGCCGAGGTGAAGTTCGAGGGCGACACCCTGGTGAACCGCATCGAGC<br>TGAAGGGCATCGACTTCAAGGAGGACGGCAACATCCTGGGGCACAAGCTG<br>GAGTACAACTACAACAGCCACAACGTCTATATCATGGCCGACAAGCAGAAG<br>AACGGCATCAAGGTGAACTTCAAGATCCGCCACAACATCGAGGACGGCAGC<br>GTGCAGCTCGCCGACCACTACCAGCAGAACACCCCCATCGGCGACGGCCC<br>CGTGCTGCTGCCCGACAACCACTACCTGAGCACCCAGTCCGCCCTGAGCAA<br>AGACCCCAACGAGAAGCGCGATCACATGGTCCTGCTGGAGTTCGTGACCGC<br>CGCCGGGATCACTCTCGGCATGGACGAGCTGTACAAGAAGCTTAGCCATGG<br>CTTCCCGCCGGCGGTGGCGGCGCAGGATGATGGCACGCTGCCCATGTCTT<br>GTGCCCAGGAGAGCGGGATGGACCGTCACCCTGCAGCCTGTGCTTCTGCT<br>AGGATCAATGTGTAAGGAAAAGTAAGGAAAACGATTCCTTCTAACAGAAATG<br>TCCTGAGCAATCACCTATGAACTTGTTTCAAATGCATGATCAAATGCAACCTC<br>ACAACCTTGGCTGAGTCTTGAGACTGAAAGATTTAGCCATAATGTAAACTGC<br>CTCAAATTGGACTTTGGGCATAAAAGAACTTTTTTATGCTTACCATCTTTTTTT<br>TTTCTTTAACAGATTTGTATTTAAGAATTGTTTTTAAAAAATTTTAAGATTTACA<br>CAATGTTTCTCTGTAAATATTGCCATTAAATGTAAATAACTTTAATAAAACGTT<br>TATAGCAGTTACACAGAATTTCAATCCTAGTATATAGTACCTAGTATTATAGG<br>TACTATAAACCCTAATTTTTTTTATTTAAGTACATTTTGCTTTTTAAAGTT |
| ECORV-MYC-GSG-P2A-D1EGFP-STOP-HPRT1-UTR-MYC-POST-UTR-ECORV (c-MYC HPRT1 3’UTR) | ACCTTCTGCTGGAGGCCACAGCAAACCTCCTCACAGCCCACTGGTCCTCAA<br>GAGGTGCCACGTCTCCACACATCAGCACAACTACGCAGCGCCTCCCTCCAC<br>TCGGAAGGACTATCCTGCTGCCAAGAGGGTCAAGTTGGACAGTGTCAGAGT<br>CCTGAGACAGATCAGCAACAACCGAAAATGCACCAGCCCCAGGTCCTCGGA<br>CACCGAGGAGAATGTCAAGAGGCGAACACACAACGTCTTGGAGCGCCAGA<br>GGAGGAACGAGCTAAAACGGAGCTTTTTTGCCCTGCGTGACCAGATCCCGG<br>AGTTGGAAAACAATGAAAAGGCCCCCAAGGTAGTTATCCTTAAAAAAGCCAC<br>AGCATACATCCTGTCCGTCCAAGCAGAGGAGCAAAAGCTCATTTCTGAAGA<br>GGACTTGTTGCGGAAACGACGAGAACAGTTGAAACACAAACTTGAACAGCT<br>ACGGAACTCTTGTGCGGGCAGCGGCGCCACAAACTTCTCTCTGCTAAAGCA<br>AGCAGGTGATGTTGAAGAAAACCCCGGGCCTGTGAGCAAGGGCGAGGAGC |

| | |
|---|---|
| | TGTTCACCGGGGTGGTGCCCATCCTGGTCGAGCTGGACGGCGACGTAAAC<br>GGCCACAAGTTCAGCGTGTCCGGCGAGGGCGAGGGCGATGCCACCTACGG<br>CAAGCTGACCCTGAAGTTCATCTGCACCACCGGCAAGCTGCCCGTGCCCTG<br>GCCCACCCTCGTGACCACCCTGACCTACGGCGTGCAGTGCTTCAGCCGCTA<br>CCCCGACCACATGAAGCAGCACGACTTCTTCAAGTCCGCCATGCCCGAAGG<br>CTACGTCCAGGAGCGCACCATCTTCTTCAAGGACGACGGCAACTACAAGAC<br>CCGCGCCGAGGTGAAGTTCGAGGGCGACACCCTGGTGAACCGCATCGAGC<br>TGAAGGGCATCGACTTCAAGGAGGACGGCAACATCCTGGGGCACAAGCTG<br>GAGTACAACTACAACAGCCACAACGTCTATATCATGGCCGACAAGCAGAAG<br>AACGGCATCAAGGTGAACTTCAAGATCCGCCACAACATCGAGGACGGCAGC<br>GTGCAGCTCGCCGACCACTACCAGCAGAACACCCCCATCGGCGACGGCCC<br>CGTGCTGCTGCCCGACAACCACTACCTGAGCACCCAGTCCGCCCTGAGCAA<br>AGACCCCAACGAGAAGCGCGATCACATGGTCCTGCTGGAGTTCGTGACCGC<br>CGCCGGGATCACTCTCGGCATGGACGAGCTGTACAAGAAGCTTAGCCATGG<br>CTTCCCGCCGGCGGTGGCGGCGCAGGATGATGGCACGCTGCCCATGTCTT<br>GTGCCCAGGAGAGCGGGATGGACCGTCACCCTGCAGCCTGTGCTTCTGCT<br>AGGATCAATGTGTAAGATGAGAGTTCAAGTTGAGTTTGGAAACATCTGGAGT<br>CCTATTGACATCGCCAGTAAAATTATCAATGTTCTAGTTCTGTGGCCATCTGC<br>TTAGTAGAGCTTTTTGCATGTATCTTCTAAGAATTTTATCTGTTTTGTACTTTA<br>GAAATGTCAGTTGCTGCATTCCTAAACTGTTTATTTGCACTATGAGCCTATAG<br>ACTATCAGTTCCCTTTGGGCGGATTGTTGTTTAACTTGTAAATGAAAAAATTC<br>TCTTAAACCACAGCACTATTGAGTGAAACATTGAACTCATATCTGTAAGAAAT<br>AAAGAGAAGATATATTAGTTTTTTAATTGGTATTTTAATTTTTATATATGCAGG<br>AAAGAATAGAAGTGATTGAATATTGTTAATTATACCACCGTGTGTTAGAAAAG<br>TAAGAAGCAGTCAATTTTCACATCAAAGACAGCATCTAAGAAGTTTTGTTCTG<br>TCCTGGAATTATTTTAGTAGTGTTTCAGTAATGTTGACTGTATTTTCCAACTTG<br>TTCAAATTATTACCAGTGAATCTTTGTCAGCAGTTCCCTTTTAAATGCAAATC<br>AATAAATTCCCAAAAATTTAAGTTGTGAATGTTTTGTTTCGTTTCTTCCCCCTC<br>CCAACCACCACCATCCCTGTTTGTTTTCATCAATTGCCCCTTCAGAGGGTGG<br>TCTTAAGAAAGGCAAGAGTTTTCCTCTGTTGAAATGGGTCTGGGGGCCTTAA<br>GGTCTTTAAGTTCTTGGAGGTTCTAAGATGCTTCCTGGAGACTATGATAACA<br>GCCAGAGTTGACAGTTAGAAGGAATGGCAGAAGGCAGGTGAGAAGGTGAG<br>AGGTAGGCAAAGGAGATACAAGAGGTCAAAGGTAGCAGTTAAGTACACAAA<br>GAGGCATAAGGACTGGGGAGTTGGGAGGAAGGTGAGGAAGAAACTCCTGT<br>TACTTTAGTTAACCAGTGCCAGTCCCCTGCTCACTCCAAACCCAGGAATTCT<br>GCCCAGTTGATGGGGACACGGTGGGAACCAGCTTCTGCTGCCTTCACAACC<br>AGGCGCCAGTCCTGTCCATGGGTTATCTCGCAAACCCCAGAG |

**Table. S5.**

| Primer sequences | | | | |
|---|---|---|---|---|
| Name | Sequence (5'-3') | Direction | Location | PCR product size |
| MYC_F | CTCCTGGCAAAAGGTCAGAGTC | Forward | Upstream of homology arm | Variable |
| MYC_R | GAACCAAGGCATGATAGCGA | Reverse | Downstream of homology arm | |
| MYC_Control3UTR_F1 | TCCTGGCAAAAGGTCAGAGTC | Forward | MYC coding sequence | 554 bp |
| MYC_Control3UTR_R1 | CTTCAACATCACCTGCTTGCT | Reverse | P2A | |
| MYC_Control3UTR_F2 | TCTCTGCTAAAGCAAGCAGGT | Forward | P2A/dGFP | 993 bp |
| MYC_Control3UTR_R2 | CTCAGCCAAGGTTGTGAGGT | Reverse | MYC 3'UTR | |
| MYC_Control3UTR_F3 | GCCGACAAGCAGAAGAACG | Forward | GFP/MYC 3'UTR | 996 bp |

| | | | | |
|---|---|---|---|---|
| MYC_Control3UTR_R3 | TAAGGCCCCCAGACCCATTT | Reverse | Downstream of homology arm | |
| MYC_HPRT13UTR_F1 | CCCACTGGTCCTCAAGAGGTG | Forward | MYC CDS/P2A | 1147 bp |
| MYC_HPRT13UTR_R1 | CTTCTCGTTGGGGTCTTTGC | Reverse | dGFP | |
| MYC_HPRT13UTR_F2 | CCCACTGGTCCTCAAGAGGTG | Forward | dGFP/HPRT1 3'UTR | 1325 bp |
| MYC_HPRT13UTR_R2 | CTTCTCGTTGGGGTCTTTGC | Reverse | Downstream of homology arm | |

**Table. S6.**

| qPCR primers | | |
|---|---|---|
| Gene Target | Forward sequence (5'-3') | Reverse sequence (5'-3') |
| c-MYC | TGGAAAACCAGCCTCCCG | TTCTCCTCCTCGTCGCAGTA |
| c-MYC-dsGFP | GTTGCGGAAACGACGAGAAC | GCCCGGGGTTTTCTTCAACA |
| dsGFP | GAGCAAAGACCCCAACGAGA | ATCCTAGCAGAAGCACAGGC |
| HPRT1 | CCTGGCGTCGTGATTAGTGA | CGAGCAAGACGTTCAGTCCT |
| GAPDH | TGAAGGTCGGAGTCAACGG | TCCTGGAAGATGGTGATGGGA |

**Table. S7.**

| Antibodies | | | | |
|---|---|---|---|---|
| | Species | Company | Cat no | Dilution used |
| Primary | | | | |
| H3K18ac | Rabbit | Abcam | ab1191 | 1:1000 |
| c-MYC | Rabbit | Cell Signalling | 9420S | 1:1000 |
| RNAPII subunit B1 (clone 3E10) | Rat | Sigma Aldrich | 04-1571-I | 1:1000 |
| ACTIN (clone AC-74) | Mouse | Sigma Aldrich | A2228 | 1:6000 |
| GFP | Rabbit | Invitrogen | A6455 | 1:1000 |
| ELAVL1 | Mouse | Santa Cruz | sc-5261 | 1:1000 |
| Laminin B1 | Rabbit | Abcam | ab16048 | 1:6000 |
| Secondary | | | | |
| Polyclonal Rabbit Anti-Mouse Immunoglobulin/HRP | Rabbit | Dako | P0260 | 1:5000 |
| Polyclonal Rabbit Anti-Rat Immunoglobulin/HRP | Rabbit | Dako | P0450 | 1:5000 |
| Polyclonal Swine Anti-Rabbt Immunoglobulin/HRP | Swine | Dako | P0217 | 1:5000 |
| IRDy 800CW Donkey Anti-Rabbit | Donkey | LICOR | 926-32213 | 1:20000 |
| IRDye 680RD Donkey Anti-Mouse | Donkey | LICOR | 926-68072 | 1:20000 |

**Table. S8.**

| MAC-seq well treatment conditions | | | |
|---|---|---|---|
| Well | Treatment | Final Concentration | Treatment duration |
| A01 | DMSO | 1:5000, | 6h |
| A02 | GSK_J4_HCl | 1uM | 6h |
| A03 | BIX_01294 | 1uM | 6h |
| A04 | MM_102 | 1uM | 6h |
| A05 | RGFP966 | 1uM | 6h |
| A06 | SGC_0946 | 1uM | 6h |
| A07 | Ofloxacin | 1uM | 6h |

| | | | |
|---|---|---|---|
| A08 | THZ1 | 1uM | 6h |
| A09 | THZ531 | 1uM | 6h |
| A10 | Triptolide | 1uM | 6h |
| A11 | AZ5576 | 1uM | 6h |
| A12 | DMSO | 1:5000, | 6h |
| B01 | DMSO | 1:5000, | 6h |
| B02 | ML324 | 1uM | 6h |
| B03 | A366 | 1uM | 6h |
| B04 | EPZ015666 | 1uM | 6h |
| B05 | PCI_34051 | 1uM | 6h |
| B06 | PFI2_HCl | 1uM | 6h |
| B07 | Lomeguatrib | 1uM | 6h |
| B08 | OTS964 | 1uM | 6h |
| B09 | ActD | 10ug/mL | 6h |
| B10 | JQ1 | 1uM | 6h |
| B11 | dBET1 | 1uM | 6h |
| B12 | DMSO | 1:5000, | 6h |
| C01 | A485 | 1uM | 6h |
| C02 | JIB_04 | 1uM | 6h |
| C03 | CPI_360 | 1uM | 6h |
| C04 | GSK591 | 1uM | 6h |
| C05 | NexturastatA | 1uM | 6h |
| C06 | UNC0379 | 1uM | 6h |
| C07 | UNC1215 | 1uM | 6h |
| C08 | IBET_762 | 1uM | 6h |
| C09 | A485 | 1uM | 6h |
| C10 | YKL5124 | 1uM | 6h |
| C11 | Ricolintostat | 1uM | 6h |
| C12 | AZ5576 | 1uM | 6h |
| D01 | A485 | 1uM | 6h |
| D02 | GSK_J1 | 1uM | 6h |
| D03 | GSK503 | 1uM | 6h |
| D04 | SGC707 | 1uM | 6h |
| D05 | Selisistat | 1uM | 6h |
| D06 | Olaparib | 1uM | 6h |
| D07 | UNC669 | 1uM | 6h |
| D08 | PCI34051 | 1uM | 6h |
| D09 | Panobinostat | 1uM | 6h |
| D10 | Vorinostat | 1uM | 6h |
| D11 | GSK503 | 1uM | 6h |
| D12 | AZ5576 | 1uM | 6h |
| E01 | JQ1 | 1uM | 6h |
| E02 | GSK_LSD1_2HCl | 1uM | 6h |
| E03 | EI1 | 1uM | 6h |
| E04 | MS023 | 1uM | 6h |
| E05 | Sirtinol | 1uM | 6h |
| E06 | Niraparib | 1uM | 6h |
| E07 | GSK2801 | 1uM | 6h |

| | | | |
|---|---|---|---|
| E08 | EED226 | 1uM | 6h |
| E09 | OKT422 | 1uM | 6h |
| E10 | A241 | 1uM | 6h |
| E11 | RGF966 | 1uM | 6h |
| E12 | ActD | 10ug/mL | 6h |
| F01 | JQ1 | 1uM | 6h |
| F02 | SP2509 | 1uM | 6h |
| F03 | CPI_169 | 1uM | 6h |
| F04 | MC1568 | 1uM | 6h |
| F05 | RG108 | 1uM | 6h |
| F06 | IBET_762 | 1uM | 6h |
| F07 | PFI3 | 1uM | 6h |
| F08 | A159 | 1uM | 6h |
| F09 | DRB | 100uM | 6h |
| F10 | OkadaicAcid | 1uM | 6h |
| F11 | Phendione | 1uM | 6h |
| F12 | ActD | 10ug/mL | 6h |
| G01 | DMSO | 1:5000, | 6h |
| G02 | BRD4770 | 1uM | 6h |
| G03 | MI-2 | 1uM | 6h |
| G04 | TMP269 | 1uM | 6h |
| G05 | SGI_1027 | 1uM | 6h |
| G06 | SGC_CBP30 | 1uM | 6h |
| G07 | MG149 | 1uM | 6h |
| G08 | GSK126 | 1uM | 6h |
| G09 | LDC4297 | 1uM | 6h |
| G10 | MS444 | 10uM | 6h |
| G11 | DMSO | 1:5000, | 6h |
| G12 | DMSO | 1:5000, | 6h |
| H01 | DMSO | 1:5000, | 6h |
| H02 | UNC0631 | 1uM | 6h |
| H03 | MI-3 | 1uM | 6h |
| H04 | Quisinostat | 1uM | 6h |
| H05 | Pinometostat | 1uM | 6h |
| H06 | Pirarubicin | 1uM | 6h |
| H07 | OF1 | 1uM | 6h |
| H08 | DMSO | 1:5000, | 6h |
| H09 | DMSO | 1:5000, | 6h |
| H10 | DMSO | 1:5000, | 6h |
| H11 | EtOH | 1:10000 | 6h |
| H12 | DMSO | 1:5000, | 6h |

**Table. S9.**

| MAC-seq well barcodes | |
|---|---|
| Well | 10bp barcode |
| A01 | AACAAGGTAC |
| A02 | AAGACGGATT |
| A03 | AACACCTAGT |

| | |
|---|---|
| A04 | CCGGCCAATT |
| A05 | CGCCAACCAT |
| A06 | CGTCCTAGGA |
| A07 | GCAAGCGAAT |
| A08 | GGCGCTATAA |
| A09 | GTCGGTGACA |
| A10 | TCACAGATAC |
| A11 | TCGCGTAGCA |
| A12 | TGTGCACTAA |
| B01 | AACAATCAGG |
| B02 | AAGATCGGCG |
| B03 | AACAGGCAAT |
| B04 | CCGTCAGAAC |
| B05 | CGCGGATTCA |
| B06 | CGTCGGCAAT |
| B07 | GCAATGTAAG |
| B08 | GGCGTTAAGT |
| B09 | GTCTCGAGTG |
| B10 | TCACCGCCTA |
| B11 | TCGGCGTTAA |
| B12 | TGTTGTGACT |
| C01 | AACATGGAGA |
| C02 | AAGCGATGTT |
| C03 | AACCAGCCAG |
| C04 | CCTAGACACG |
| C05 | CGCTTAAGGC |
| C06 | CTAACTTCAG |
| C07 | GCACACTATA |
| C08 | GGTAATGTGT |
| C09 | GTCTCTTAAG |
| C10 | TCATTGTCCA |
| C11 | TCTCTCCTAT |
| C12 | TTAACGCTGA |
| D01 | AACATTACCG |
| D02 | AAGCGTTCAG |
| D03 | AACCAGTTGA |
| D04 | CCTCAACCGA |
| D05 | CGCTTACTAA |
| D06 | CTAATAGCGT |
| D07 | GCACTTAATC |
| D08 | GGTGGTTGGA |
| D09 | GTCTTCCGAG |
| D10 | TCCACACTAG |
| D11 | TCTTGCTCGG |
| D12 | TTATAGGAGG |
| E01 | AACCGCGACT |
| E02 | AAGGTCTGGA |
| E03 | AACCGGCGTA |

| | |
|---|---|
| E04 | CCTCCATAAG |
| E05 | CGGCAACTTA |
| E06 | CTATGAACGG |
| E07 | GCAGGAGATG |
| E08 | GTAACCTTGG |
| E09 | GTGTGCCTGT |
| E10 | TCCGACTAAC |
| E11 | TGACCTGAGA |
| E12 | TTATCGCGTT |
| F01 | AACCGGAAGG |
| F02 | AAGTTAGCGC |
| F03 | AACCTAGTCC |
| F04 | CCTTGTATTC |
| F05 | CGGCTCATCA |
| F06 | CTCAAGGACC |
| F07 | GCATCCGATC |
| F08 | GTAAGAACCT |
| F09 | GTGTGTGTCC |
| F10 | TCCGTTATCT |
| F11 | TGCGCTCATT |
| F12 | TTCAGGAGTA |
| G01 | AACCTCATAG |
| G02 | AATAGCCACA |
| G03 | AACTCTACAC |
| G04 | CGAGATCTCT |
| G05 | CGTAACGGAT |
| G06 | CTCGCAACGT |
| G07 | GGACGATGCT |
| G08 | GTATTGTGGA |
| G09 | GTTCGTCGAA |
| G10 | TCGAAGCATT |
| G11 | TGCGTGCTCA |
| G12 | TTCCATCGAG |
| H01 | AACGTAAGCT |
| H02 | AATCACGCGA |
| H03 | AACTGTGTCA |
| H04 | CGATCCTGTG |
| H05 | CGTAAGATTC |
| H06 | CTCGTGCCTA |
| H07 | GGCATCGTGA |
| H08 | GTCCGCATCA |
| H09 | GTTGAATTGG |
| H10 | TCGAGAGAGC |
| H11 | TGTGACGTGC |
| H12 | TTGGCAATTC |

#### Supplementary references


1. C. J. O. Alexander J. Federation, Donald R. Polaski, A. Fan, C. Y. Lin, J. E., Bradner, Identification of candidate master transcription factors within enhancer-centric transcriptional regulatory networks. *bioRxiv* (2018).
2. M. Huang, Y. Chen, M. Yang, A. Guo, Y. Xu, L. Xu, H. P. Koeffler, DbCoRC: A database of core transcriptional regulatory circuitries modeled by H3K27ac ChIP-seq signals. *Nucleic Acids Res.* **46**, D71–D77 (2018).
3. M. Wu, E. Karadoulama, M. Lloret-Ilinares, J. O. Rouviere, C. S. Vaagensø, M. Moravec, B. Li, J. Wang, G. Wu, M. Gockert, V. Pelechano, T. H. Jensen, A. Sandelin, The RNA exosome shapes the expression of key protein-coding genes. *Nucleic Acids Res.*, 1–20 (2020).
4. J. A. Schofield, E. E. Duffy, L. Kiefer, M. C. Sullivan, M. D. Simon, TimeLapse-seq: Adding a temporal dimension to RNA sequencing through nucleoside recoding. *Nat. Methods*. **15**, 221–225 (2018).
5. M. Muhar, A. Ebert, T. Neumann, C. Umkehrer, J. Jude, C. Wieshofer, P. Rescheneder, J. J. Lipp, V. A. Herzog, B. Reichholf, D. A. Cisneros, T. Hoffmann, M. F. Schlapansky, P. Bhat, A. Von Haeseler, T. Köcher, A. C. Obenauf, J. Popow, S. L. Ameres, J. Zuber, SLAM-seq defines direct gene-regulatory functions of the BRD4-MYC axis. *Science (80-. ).* **360**, 800–805 (2018).
6. S. J. Vervoort, S. A. Welsh, J. R. Devlin, N. Gray, A. Gardini, R. W. Johnstone, The PP2A-Integrator-CDK9 axis fine-tunes transcription and can be targeted therapeutically in cancer. *Cell*, 1–20 (2021).
